# Protosequences in brain organoids model intrinsic brain states

**DOI:** 10.1101/2023.12.29.573646

**Authors:** Tjitse van der Molen, Alex Spaeth, Mattia Chini, Sebastian Hernandez, Gregory A. Kaurala, Hunter E. Schweiger, Cole Duncan, Sawyer McKenna, Jinghui Geng, Max Lim, Julian Bartram, Aditya Dendukuri, Zongren Zhang, Jesus Gonzalez-Ferrer, Kiran Bhaskaran-Nair, Lon J. Blauvelt, Cole R.K. Harder, Linda R. Petzold, Dowlette-Mary Alam El Din, Jason Laird, Maren Schenke, Lena Smirnova, Bradley M. Colquitt, Mohammed A. Mostajo-Radji, Paul K. Hansma, Mircea Teodorescu, Andreas Hierlemann, Keith B. Hengen, Ileana L. Hanganu-Opatz, Kenneth S. Kosik, Tal Sharf

**Author notes:** Correspondence (T.S.).

## Abstract

Neuronal firing sequences are thought to be the basic building blocks of neural coding and information broadcasting within the brain. However, when sequences emerge during neurodevelopment remains unknown. We demonstrate that structured firing sequences are present in spontaneous activity of human and murine brain organoids and *ex vivo* neonatal brain slices from the murine somatosensory cortex. We observed a balance between temporally rigid and flexible firing patterns that are emergent phenomena in human and murine brain organoids and early postnatal murine somatosensory cortex, but not in primary dissociated cortical cultures. Our findings suggest that temporal sequences do not arise in an experience-dependent manner, but are rather constrained by an innate preconfigured architecture established during neurogenesis. These findings highlight the potential for brain organoids to further explore how exogenous inputs can be used to refine neuronal circuits and enable new studies into the genetic mechanisms that govern assembly of functional circuitry during early human brain development.

## Introduction

In the last decades, a growing body of experimental evidence has begun to support the notion that intrinsic activity plays a central role in brain function, challenging the traditional Jamesian view that higher order function is an emergent product of sensory input^1^. More recently, analysis of the mesoscale wiring of the cortex has revealed a connectome dominated by recurrent cortical networks that follow lognormal scaling rules that are conserved across species, where only a minor portion of those connections are devoted to direct transfer of sensory input^2–4^. Within cortical circuitry, both spatial and functional properties of neurons follow lognormal distributions, which include synaptic strengths^5^, size of dendritic spines^6^, diameter of axons^7^ and neuronal firing rates^8^, which together establish the functional architecture of the large-scale brain connectome^2,3,9,10^. The skewed distribution of these parameters is thought to serve as a substrate of neuronal assemblies^11,12^, where groups of strongly interconnected neurons can generate temporally structured spiking activity, which organize into sequences^13–15^. Sequential activity patterns represent discrete and temporally consolidated packets of neuronal activity proposed as the basic building blocks of neural coding and information broadcasting within the brain^16^. In mature brain circuits, spiking sequences are predictive indicators for spatial navigation tasks^17^ and memory formation within the murine hippocampus^18^, and through a phenomena called ‘preplay’ are used to encode novel experience through a pre-existing repertoire of sequence motifs^19–23^. Such sequences emerge prior to explicit experience-dependent navigational representations (e.g. before exploration beyond the nest)^24^. Similarly, in the murine visual cortex, evoked responses closely mirror spontaneous sequential patterns^25^. Similar phenomena have also been reported in the human cortex, where the replay of sequences underlies episodic memory formation and retrieval^26^. Moreover, spiking sequences in the human anterior temporal lobe organize into a temporal backbone. This backbone consists of both rigid and flexible sequence elements that are stable over time and cognitive states^27^, support visual categorization tasks in the human visual cortex and were shown to encode non-redundant information beyond latency and rate encoding^28^. Reliable activation of sequences are also present in the reptilian cortex that share a common primordial ancestral origin with mammals, which suggests a conserved function across phylogeny^14^. However, the emergence of spiking sequences during development is not yet well understood. During the third postnatal week, the murine hippocampus generates spiking sequences that resemble those that will later be produced during navigation in a linear environment^24^. Importantly, these sequences emerge in an experience-independent manner and do not improve upon additional experience in the same postnatal week. Whether similar spiking sequences, potentially representing other forms of experience, exist in other brain areas or at earlier developmental stages remains an open question. The existence of such sequences would provide strong evidence in support of the notion that the brain is in a preconfigured state, where spiking sequences are not experience-dependent but are instead constrained by an innate architecture that is established during neurogenesis^29^. To address this open question, we investigated large scale single-unit activity datasets recorded from different models of brain development: (1) human-derived brain organoids generated by two independent laboratories^30,31^, (2) murine-derived brain organoids of dorsal forebrain identity grown from mouse embryonic stem cells (ESCs), (3) *ex vivo* murine brain slices from the somatosensory cortex and, (4) dissociated two-dimensional primary murine cortical cultures^32,33^ that lack developmental organization.

Brain organoids are three-dimensional human induced pluripotent stem cell (iPSC) derived models of the human brain that recapitulate key facets of the anatomical organization and cellular composition found in the developing brain^34–37^. Neurons within brain organoids form functional synapses^36^ and establish spontaneous network activity^38–40^. These self-organized neuronal systems contain cellular diversity and cytoarchitecture necessary to sustain complex network dynamics^30,41,42^ as evidenced by expression of layer specific excitatory pyramidal neurons and generation and incorporation of inhibitory GABAergic interneurons^43,44^. Brain organoids also generate local field potential oscillations (LFP) that mirror preterm EEG patterns^41^, and have been used to model network dynamics associated with rare genetic disorders^42^. Brain organoids can be readily interfaced with state-of-the-art high-resolution CMOS microelectrode arrays to record neuronal activity and LFP oscillations across 26,400 recording sites^30^. Furthermore, brain organoids are not “connected” to any sensory system, and thus represent an ideal model for studying the emergence of spiking sequences as a truly experience-independent phenomenon.

The *ex vivo* rodent dataset consists of acute neonatal slice recordings from the murine somatosensory cortex. At this developmental stage, the somatosensory cortex is characterized by discontinuous activity, displaying an alternation of activity bursts with periods of electrical quiescence^45^, and highly correlated activity between spike trains^46,47^. Both traits are shared with several other cortical areas^48,49^. Importantly, at this stage, with the exception of the olfaction that controls cognitive maturation^50^, sensory systems are still underdeveloped^51^. Accordingly, whisker-elicited sensory responses are mainly the result of passive stimulations by the dam and the littermates in the first postnatal week, while active whisking only emerges around P10-12^52–54^. Along the same lines, analogously to neural activity in brain organoids, a large portion of neural activity in the developing brain is “spontaneous”, that is, internally generated^55^.

Here, we investigate the spontaneous firing patterns across these different developmental brain models. In all four models, we observed bursting dynamics on the order of 10^2^ milliseconds. These bursts reflect the biophysical time constants for the integration of inputs in neural circuits^56^, which last longer than single-neuron refractoriness and burstiness alone^57^. We identified a subpopulation of neurons within organoids and neonatal brain slices capable of generating and sustaining non-random sequential firing patterns, referred to as backbone sequences^27^. In contrast, two-dimensional primary cultures did not exhibit backbone sequences. Backbone sequence generating neurons populate the tails of right-skewed, lognormal firing rate and functional connectivity distributions. We also reveal that, at a population level, neural activity exhibits a distinct low and high-dimensional neuronal subspace that establishes a partition between sequence generating units and their non-rigid counterparts. In an *in vivo* setting, such firing patterning temporally segregate neuronal populations into strongly correlated, ‘temporally rigid’ components, obedient to population dynamics that reside in a low-dimensional subspace, and weakly correlated, ‘temporally flexible’, firing patterns that are less sensitive to population events^58^. Finally, for stable and flexible computation, theory suggests that neural systems must exist at or near a regime called *criticality*, which is characterized by scale-invariant dynamics across multiple timescales^59–61^. At criticality, complex patterns of activity emerge that neither decay rapidly nor saturate the network, maximizing the system’s computational capacity. Using temporal renormalization group theory^62^, we quantified the distance of each preparation from criticality (*d_2_*). This approach revealed that brain organoids, *ex vivo* brain slices, and two-dimensional primary cultures can generate near-critical dynamics. A subset of each type of preparation exhibited temporal correlations that spanned multiple timescales. In contrast, shuffled data failed to capture scale free features of activity. Given that neuronal circuits within brain organoids establish non-random, sequence generating dynamics that emerge in a truly experience-independent manner and maintain near-critical dynamics, these findings have important implications for understanding organizational principles that govern network preconfiguration and activity patterns that support a neural code^63^.

## Results

### Neuronal firing patterns in human brain organoids generate repetitive and variable burst sequences

We investigated the temporal dynamics of spontaneous neuronal activity generated by brain organoids of prominent forebrain identity^30^ using high-density CMOS-based microelectrode arrays. These arrays contain 26,400 recording sites (electrode pitch of 17.5 µm) of which 1,024 can be selected and recorded simultaneously^64^. Single-unit spike events, visualized as a raster plot (Fig. 1A) reveal the temporal evolution of spontaneous spiking dynamics in an 8-month old organoid slice positioned on the array. Population level burst events are highlighted by sharp increases in the population rate (red line Fig. 1A, see Methods) and persist for several hundred milliseconds, followed by longer periods of relative quiescence lasting up to several seconds. Moreover, we observed that the distribution of single-unit firing rates follows a heavy-tailed and right-skewed distribution, well described by a lognormal function, a feature consistent across multiple brain organoids (*n* = 8 organoids, *R*^2^ = 0.97 ± 0.04, Supplementary Fig. 1A-C). This represents one facet of functional activity conserved across brain regions and states *in vivo*^11^.

**Figure 1.**
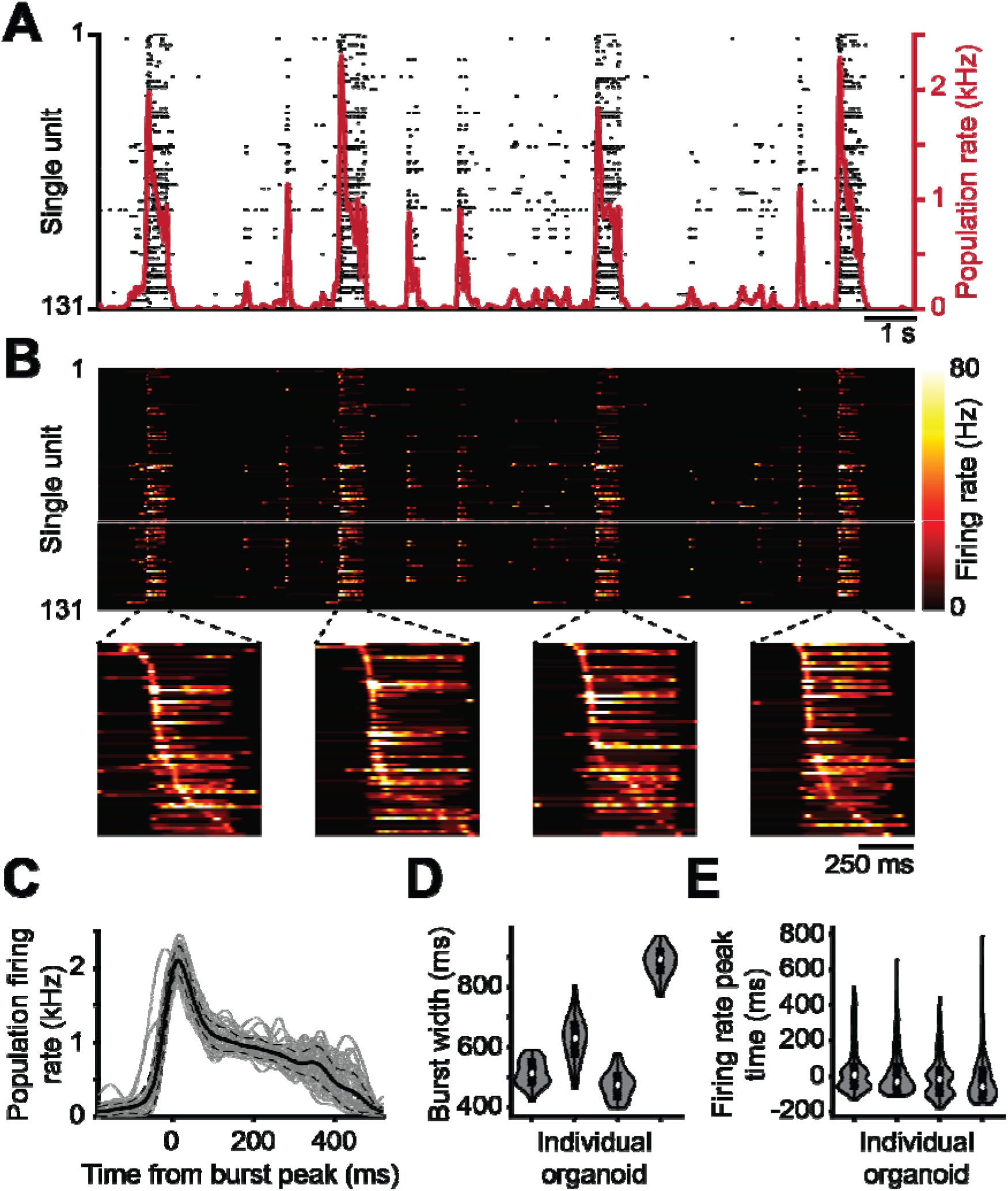
Temporal structure of spontaneous single-unit neuronal firing patterns during population bursts in human brain organoids. (**A**) Raster plot visualization of single-unit spiking (black dots) measured across the surface of a 500 μm thick human brain organoid slice, positioned on top of a microelectrode array. The population firing rate is shown by the red solid line. Population bursts are marked by sharp increases in the population rate. Burst peak events are denoted by local maxima that exceed 4x-RMS fluctuations in the population rate. The shaded gray regions denote the burst duration window as defined by the time interval in which the population rate remains above 10% of its peak value in the burst. (**B**) Top, the instantaneous firing rate of single-unit activity from panel **A**. Bottom, zoomed in view of the trajectory of neuronal firing during population bursts reveals temporal segregation and contiguous tiling of the peak firing rate of single-unit activity. Here the subset of units that fire at least two times during the burst are shown, re-ordered for each burst individually based on the time relative to the burst peak at which the unit has its maximum firing rate during the burst period. (**C**) The population firing rate (gray lines) is plotted relative to the burst peak for 46 burst events measured across a three-minute interval for the same organoid. The mean value is shown by the solid line and the dotted lines represent 1 STD. (**D**) Burst durations are plotted from four different organoids. (**E**) The distribution of single-unit firing rate peak times relative to population burst peaks for the same organoids as in **D**.

To investigate the temporal structure and dynamics of single-unit neuronal firing patterns, we next calculated the instantaneous firing rate from single-unit spike times across all units (Fig. 1B, see Methods for details). Reordering units in time, based on their peak firing rate over the burst width, revealed the presence of sequential activation patterns during individual burst events (Fig. 1B, bottom). Here, we observe bursts with consistent population rate profiles within single organoids (Fig. 1C) and durations that span ≈10^2^ millisecond timescales (Fig. 1D, Supplementary Fig. 2C), a feature consistent across multiple organoids. Furthermore, we observe that a majority of the units display a firing rate peak that is in close temporal proximity to the population rate peak (Fig. 1E). Similar bursting time frames are generated by spontaneous and evoked sequences *in vivo* within the murine^65^ and reptile^14^ cortex, the murine hippocampus during sharp wave ripples^66^ and within the human cortex during memory retrieval^26,27^. Interestingly, the neuronal response time of sensory cortices typically peak with a similar time constant across brain regions and species^67–70^.

Next, we quantified the degree of stereotypy exhibited by single-unit firing patterns during spontaneous bursts in organoids. First, single-units were separated into two populations: backbone units were defined as units that spike at least twice in all bursts (Fig. 2A, above the dashed line); all other units were defined as non-rigid units (Fig. 2A, below the dashed line). Note that the backbone units display high firing rates residing within the tail of a lognormal distribution (Supplementary Fig. 1D). The relative temporal delays between single-unit firing patterns remained consistent across multiple burst events when preserving unit ordering (Fig. 2B) and was preserved when averaging the firing rate of each unit across all burst events (Fig. 2C, see Supplementary Figs. 3A-D for visualizations and firing statistics across organoids). When clustering bursts based on their pairwise correlation matrix^71^, the consistent backbone activity patterns remained similar across all clusters, whereas non-rigid units showed significantly larger variability across clusters, (*P* < 10^−20^ for difference between backbone and non-rigid, linear mixed-effects model; Supplementary Fig. 4). Interestingly, we observed a general decrease in the relative abundance of backbone units compared to earlier developmental time points. This trend was observed in 5 of 7 organoids where we tracked the long term developmental trajectory of the murine and human brain organoids spanning 6 to 8 months/weeks in human and murine organoids respectively (Supplementary Fig. 5A,B). *In vivo*, network synchronization and subsequent maturation is marked by the incorporation of interneurons within maturing excitatory networks^49^, which are also present in our human brain organoids (Supplemental Fig. 6, 7) and murine cortical organoids (Supplemental Fig. 8-10) at these time points.

**Figure 2.**
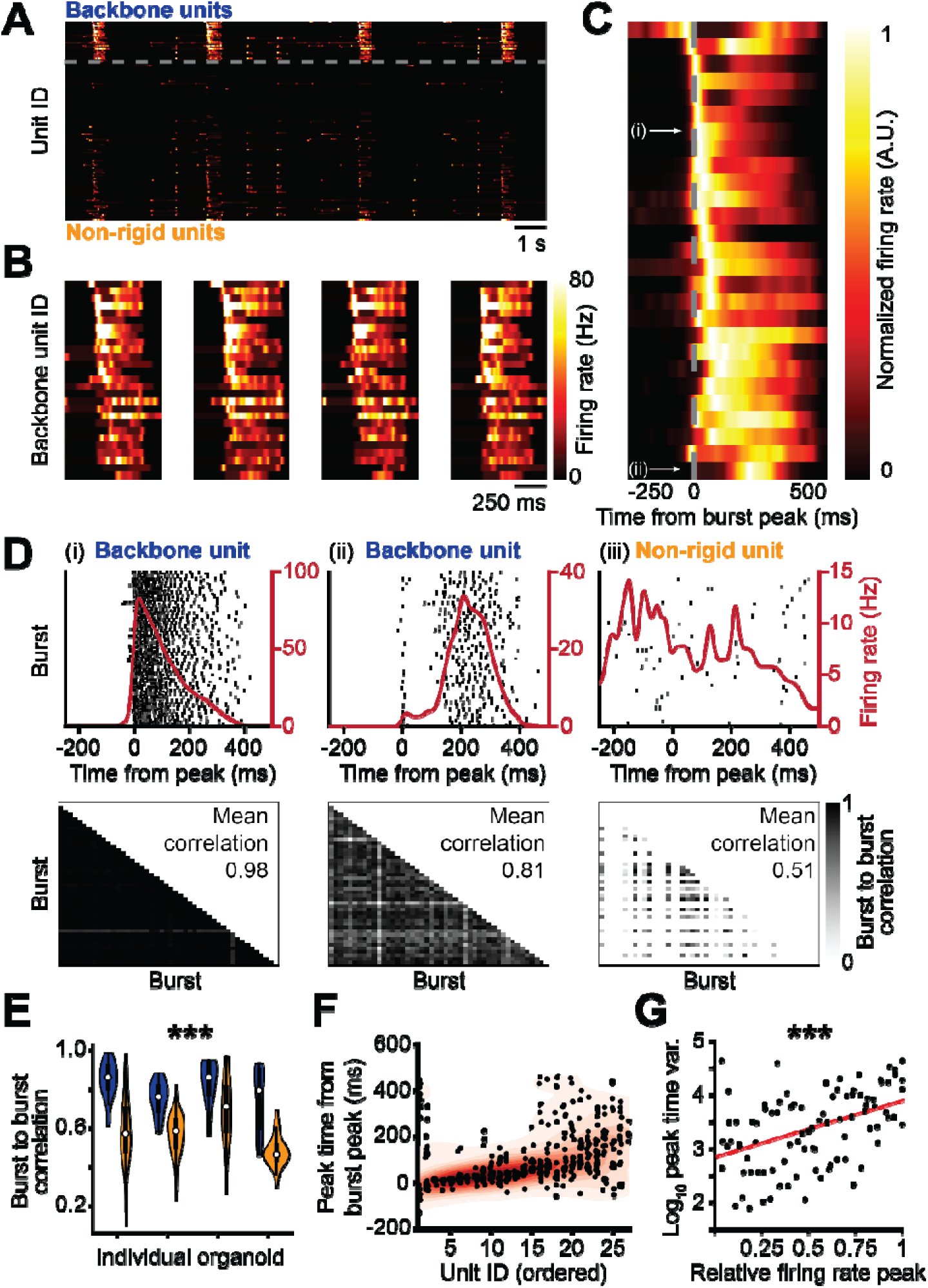
Recurring sequential activation patterns in human brain organoids generate a stereotyped temporal backbone. (**A**) Consistent firing single-units form temporally distant sub-groups (re-ordered from Fig. 1A) and exhibit temporally rigid and non-rigid firing patterns. The rigid backbone units are plotted above the dashed line and are defined by spiking at least two times in every burst epoch. Units that do not meet this criterion (non-rigid) are plotted below the dashed line. In each category, units are ordered based on their median firing rate peak time relative to the burst peak considered over all bursts in the recording. (**B**) Zoomed in view of the units from the upper dashed partition in **A** for the four bursts. The order of each unit is the same for all four plots. Note the similarity in firing pattern for each single-unit over the four different bursts. (**C**) The average burst peak centered firing rate measured across 46 burst events. The burst peak is indicated by the dotted line. The unit order is the same as **A,B**. Note the progressive increase in the firing rate peak time relative to the burst peak, as well as a spread in the active duration for units having their peak activity later in the burst. The average firing rate is normalized per unit to aid in visual clarity. (**D**) Top left, spikes are plotted for a regularly firing backbone unit over a fixed time window relative to population burst peaks. Each row represents spike events in a unique burst event. The average burst peak centered firing rate of the unit is plotted as the red solid line. Top middle, another regular firing backbone unit is plotted with a larger temporal delay and spread relative to the burst peak. Top right, firing patterns of a non-rigid unit, exhibiting poor temporal alignment relative to the burst peak events. Bottom, heatmap visualizations showing the cross-correlation coefficients of the burst peak centered firing rates of the unit shown in the top. The correlation is computed for each pair of bursts, using a maximum lag of 10 ms. The average over all burst pairs is denoted in the top right of the heatmap. (i-ii) highlight units with consistent firing patterns relative to the burst peak and have average correlation scores of 0.98 and 0.81 respectively. They are part of the backbone units as marked in **C**. (iii) illustrates an irregular firing neuron with an average correlation score of 0.51. This unit is not reliably recruited to spike within the burst. The average correlation is computed only over pairs of bursts in which this unit fired at least 2 times. (**E**) Reliably firing backbone units (colored blue) retain higher average burst-to-burst correlation coefficients across all burst instantiations, while non-rigid units (colored yellow) have significantly lower temporal correlation values. This trend is consistently observed across multiple organoids (*P* < 10^-16^, linear mixed-effects model for average burst-to-burst correlation coefficients of backbone units compared to non-rigid units). See Fig. 7B for the results of the statistical comparison of firing rate normalized data between different model types. (**F**) Firing rate peak times relative to the burst peak for each backbone unit as shown and ordered in **C**. The black dots indicate the relative firing rate peak times per burst and the red shading reflects the probability distribution of the firing rate peak times where warming colors indicate higher probability. The probabilities highlight the widening of the distribution towards the end of the sequence. (**G**) The variance of the relative firing rate peak times for the backbone units in each of the 4 presented organoids. The units are ordered based on their median firing rate peak time over all bursts to visualize the significant increase in variability of the firing rate peak times of the units that fire later in the burst (*P* < 10^-5^, linear mixed-effects model for relation between relative firing rate peak and peak time variance).

To further test the impact of inhibitory tone on backbone neurons, we next administered gabazine (10 μM) via bath application to block GABA_A_ receptors (Supplementary Fig. 11). This resulted in a significant increase in both the number of bursts and the fraction of backbone neurons participating in burst events (Supplementary Fig. 11B,C). Additionally, the rank-order statistics of sequence generating neurons increased compared to control conditions, a result consistently observed across 5 cortical organoids (Supplementary Fig. 11D). Meanwhile, the bath application of NBQX (10CµM) and R-CPP (20CµM), which block AMPA and NMDA receptors and thereby inhibit the fast and slow components of excitatory synaptic transmission, abolished bursting dynamics, as previously reported^30^, and completely suppressed neuronal sequences (Supplementary Fig. 11B). Together, these results highlight the importance of GABAergic signaling on spike timing and its impact on temporal rigidity of neuronal sequences.

To quantify the consistency of neuronal firing within spontaneous population bursts, we investigated the activity of single-units relative to the burst peak, similarly to the approach previously used for *in vivo* spontaneous activity^65,72^. Certain units displayed a pronounced peak in their firing rate and narrow temporal jitter when referenced to the burst peak (Fig. 2D, left panel), whereas others displayed increased delays and temporal spread (Fig. 2D, middle panel). A larger fraction of units, however, did not exhibit a clear preference in spike timing relative to the population burst and had more random temporal dynamics (Fig. 2D, right panel). The relative fraction of consistent firing backbone units constitutes 28% ± 14% (mean ± STD) of the total units (*n* = 8 organoids, Supplementary Fig. 2) and represents a subpopulation with significantly higher burst-to-burst correlation scores (Fig. 2E, see Methods). The consistency in the firing patterns of the backbone units was stable across recording intervals that spanned multiple hours (Supplementary Fig. 12), and the enhanced consistency of the backbone units were present at developmental time points that spanned multiple months, which include a significant increase in burst-to-burst-correlations for both murine and human brain organoids (Supplementary Fig. 5C, P < 10^−5^, two-way ANOVA). Moreover, differences in burst-to-burst correlations between backbone and non-rigid units were significantly larger when compared to spike trains that were randomized using a method that preserved both the population and single-unit mean firing rates (Supplementary Fig. 13A,B and Supplementary Fig. 14, see Methods)^58,73^. Further, this randomization destroyed the preservation of sequences across burst events (Supplementary Fig. 13C,D, Supplementary Fig. 15), similar to two-dimensional primary neuronal cultures with inherently randomized network architecture that also did not show sequential firing patterns (Supplementary Fig. 16, see section ‘Comparing sequences across neurodevelopmental models’)^74–76^. In addition, we observed that the variability of the firing rate peak time increased as a function of its average peak time (Fig. 2F), a significant feature preserved across multiple organoids (Fig. 2G, P < 10^−5^, linear mixed-effects model for relation between relative firing rate peak and peak time variance). Together these results suggest that brain organoids are capable of supporting stereotypical sequential activation patterns with increasing variability that mirror the spread of spontaneous activity through local cortical circuits *in vivo*^13,14,26,27^.

### Backbone units are a highly correlated ensemble

To further quantify the firing patterns that emerge during spontaneous burst events, we investigated pairwise correlations between single-unit instantaneous firing rates. Stereotyped activation patterns were reliably generated by backbone units and were preserved across burst events (Fig. 3A). Our analysis of sequential co-activation of these units revealed the preservation of firing rate onsets and peak activity times across all burst events with average peak phase lags of ≈10 milliseconds (Fig. 3B, example units *a* to *b* = 5ms, example units *b* to *c* = 7ms) as well as phase lags crossing several hundred milliseconds among pairs of backbone units that correlated over long durations thanks to the recurring sequential firing patterns (Supplementary Fig. 17). Cross-correlation analysis between all single-unit pairs reveals that units firing within the backbone sub-population form a highly correlated ensemble with non-zero phase lags (Fig. 3C,D). Moreover, backbone units displayed significantly higher correlation coefficients when compared to non-rigid units (Fig. 3E), and occupied the tail of the overall distribution (Fig. 3F), which is well described by a lognormal distribution (Supplementary Fig. 18C-E). Together, these results suggest that a minority population of high firing rate neurons are strongly tuned to population dynamics and function as a stop-watch in the backbone among the more rigid units of the population.

**Figure 3.**
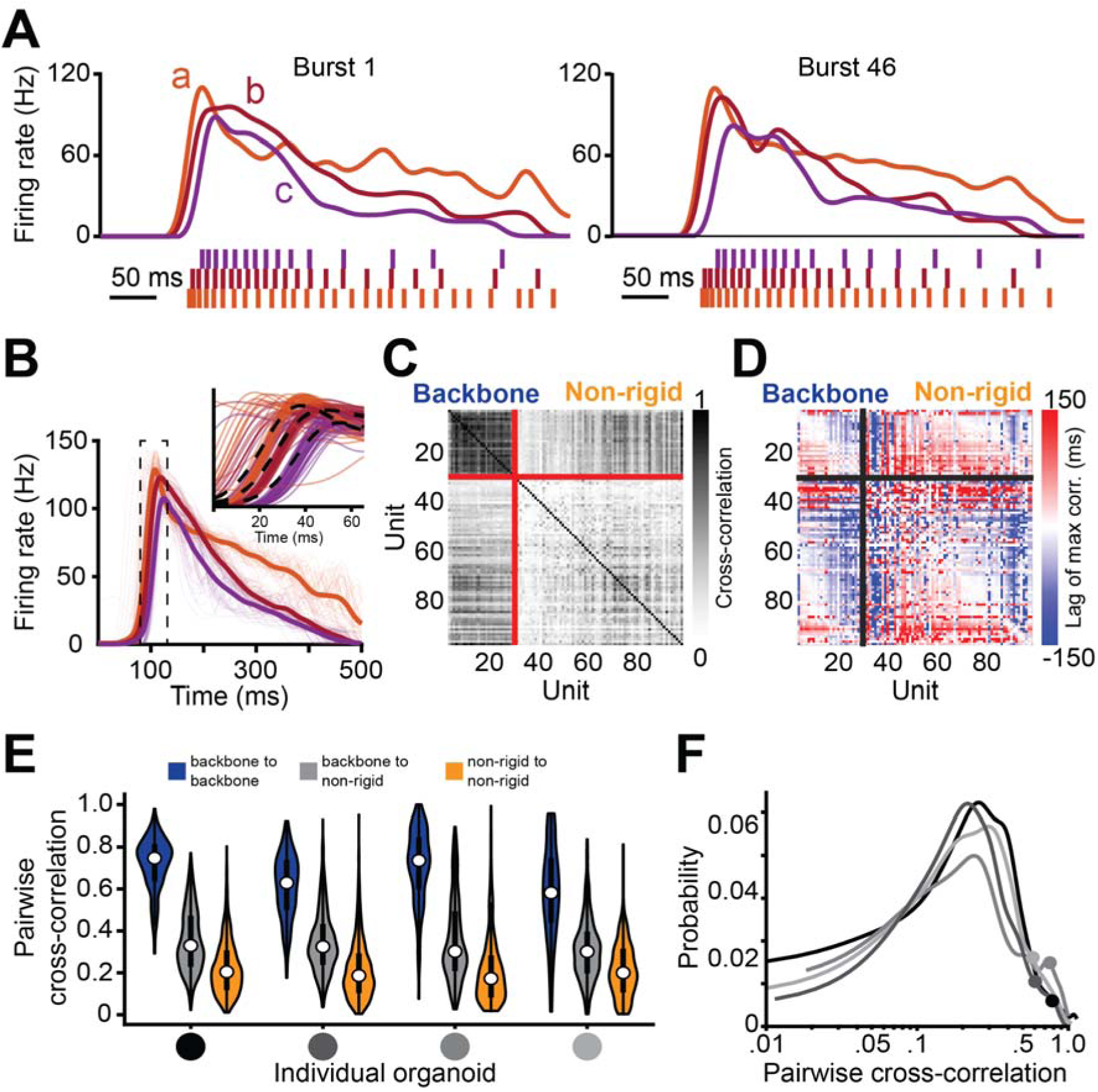
Firing patterns between single backbone units within bursts are nonrandom. **(A)** Spike times and computed firing rates for three representative units are shown for the first and last burst event of the recording, respectively. (**B**) Burst peak centered average firing rates for the three units shown in **A** are calculated over all burst events. The narrow lines indicate the firing rates for each individual burst event. The thick lines (dotted black lines in inset) indicate the average over all bursts. (**C**) Pairwise cross correlation coefficients computed between the instantaneous firing rates of all pairs of units with at least 30 spikes counted over all burst events. A maximum lag time of 350 ms was used. The solid red lines separate the backbone units and the non-rigid units. (**D**) lag times leading to the maximum cross correlation values presented in **C**. Values are clipped at ±150ms. (**E**) Pairwise cross-correlation scores are plotted between unit types. Correlations between backbone units (blue) are significantly higher (*P* < 10^-20^, linear mixed effects model) than the cross-correlations between pairs of backbone and non-rigid units (gray) and pairs of non-rigid units (yellow). See Fig. 7C for the results of the statistical comparison of firing rate normalized data between different model types. (**F**) The histogram of all pairwise cross-correlations follows a skewed, lognormal distribution (*x*-axis is log scale). The pairwise connections between pairs of backbone units populate the tail of this distribution for all organoids as can be seen from the distribution means of the backbone-to-backbone distributions in **E** and marked on each histogram (circles on line).

### Population firing rate vectors preserve timing across burst epochs

In previous sections, we focused our analysis on discrete relationships between pairwise single-unit activity. Next, we asked if single-unit firing rates were temporally structured in time during burst events at the population level. To perform this analysis, we calculated the cosine similarity between instantaneous firing rates across bursts (see Methods). This analysis revealed a peak in the cosine similarity coinciding with the firing rate increase of the backbone units. Subsequently, after a brief plateau, cosine similarity decayed and bottomed out after the burst ended (Fig. 4A-B). The variance of the similarity between bursts displayed an almost opposite trend: it was high during non-bursting periods, rapidly decreased when the similarity peaked, and remained low until burst termination (Fig. 4C).

**Figure 4.**
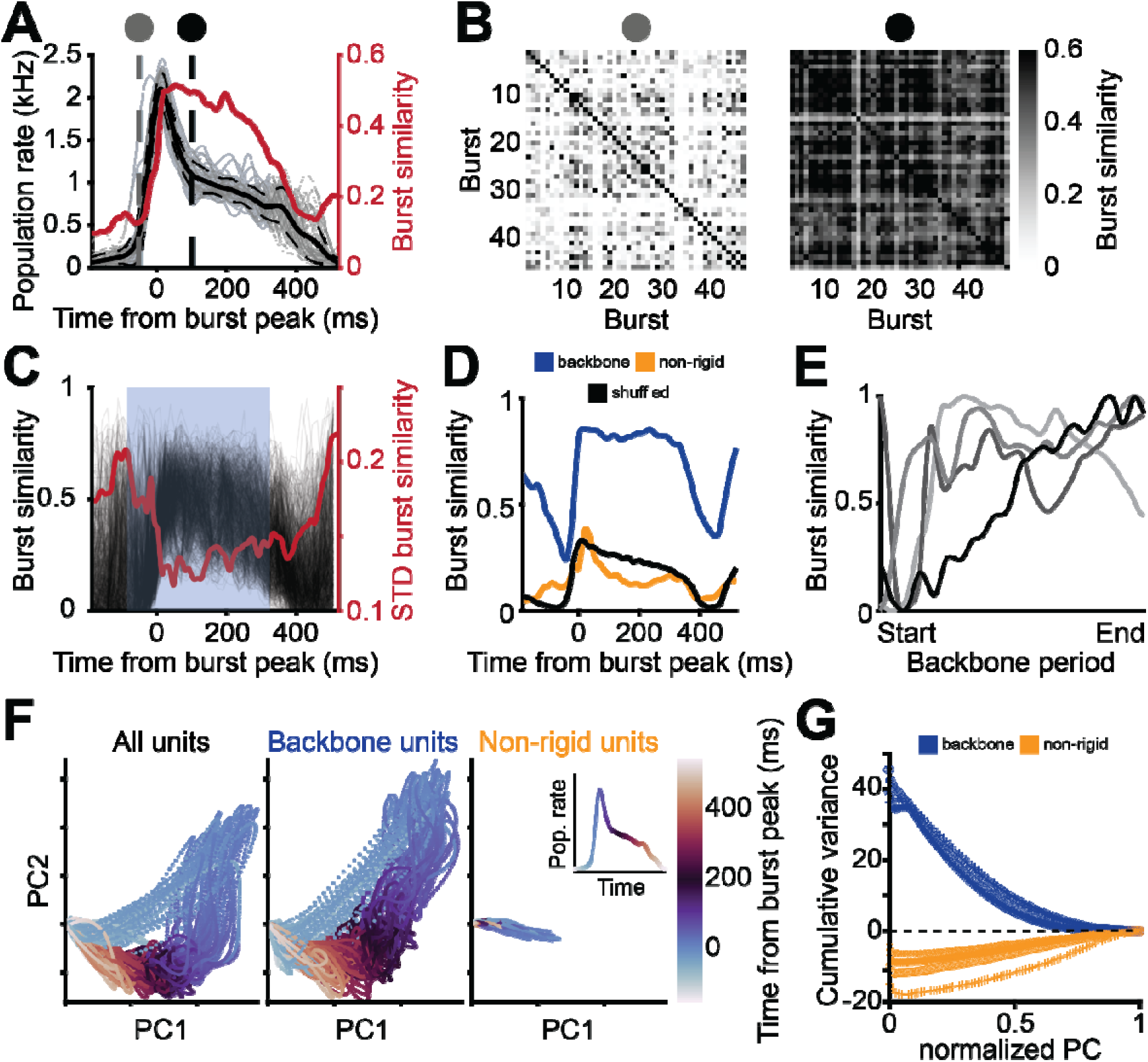
Backbone units provide a stable, low-dimensional reference frame for the population bursts. (**A**) The population firing rate (gray lines) is plotted relative to the burst peak for 46 burst events measured across a three-minute interval for the same organoid. Mean and STD are denoted by the solid black line and the dashed lines on either side of the solid black line, respectively (left axis). The average burst similarity score is plotted in red (right axis, see Methods for more details). The blue box indicates the period from the earliest average firing rate peak time of all backbone units until the last. (**B**) Left, burst similarity for each pair of bursts at 50 ms before the burst peak (indicated by the left dashed line in **A**). Right, burst similarity for each pair of bursts at 100 ms after the burst peak (indicated by the right vertical dashed line in **A**). (**C**) Burst similarity relative to the burst peak for each pair of bursts. Each gray line reflects a burst pair and the red line reflects the standard deviation per recording frame over all burst pairs. The blue box indicates the period from the earliest average firing rate peak time of all backbone units until the last. (**D**) Average burst similarity when only the backbone units or the non-rigid units are considered and for shuffled data. For each frame relative to the burst peak, the difference between the distribution for the backbone units and the non-rigid units was assessed using a paired sample, two-sided t-test which was significant throughout the backbone period (*P* < 10^-20^). (**E**) The average burst similarity throughout the backbone period follows a similar pattern across different organoids, denoted by an increase following the start of the backbone period. (**F**) Left, population activity projected onto its first two principal components. A PCA was performed on the firing rates per single-unit for all units in the recording combined. Only spikes that occurred during bursts were included in the firing rate computations. Each dot represents a recording frame and is colored by the time point relative to the closest burst peak. Note the consistent circular trajectory reflecting the burst manifold. Middle, same as left but only including the backbone units. Note the similarity in the low-dimensional manifold representations between the middle and the left, indicating the strong contribution of the backbone units to the low-dimensional activity of the whole population. Right, same as left but only including the non-rigid units. The low-dimensional manifold is not present anymore and the variance explained by the first to principal components is notably lower, reflecting the higher dimensional activity of the non-rigid units. The inset shows the burst peak centered average population rate colored in the same way as the PCA trajectories to indicate which parts of the burst correspond to the different coloring. (**G**) The difference between the cumulative sum of the variance explained by the selection of the first principal components of the manifold constructed based on only backbone units (blue) or only non-rigid units (yellow) relative to the manifold constructed based on all units. For the comparison, the cumulative sum of the explained variance is interpolated to account for the difference in the total number of principal components possible between the manifold constructed of all units compared to the subsets. The interpolated values are normalized to range from 0 (only first component) to 1 (all components). Positive values indicate that the first principal components of the manifold constructed of the subset of units explains a higher percentage of the variance than the comparable selection of principal components of the manifold constructed of all units while negative values indicate a lower percentage of explained variance. Note that for all organoids, indicated with different markers, the explained variance by the first principal components of the manifold constructed from the backbone units explain a larger amount of the total variance than the manifold constructed for all units while the opposite is true for the non-rigid. This reflects the low-dimensional backbone activity related to their heightened correlations.

To further dissect the composition of firing rate vectors and their contribution to spontaneous burst patterns, we split the population into backbone and non-rigid units. Here, we observed that the backbone ensembles exhibited significantly higher cosine similarity scores (see Fig. 4D, blue line) compared to non-rigid units (yellow line), and shuffled data (black line). To illustrate this trend, we showed that the top 20^th^ percentile of units (based on their average correlation from the matrix shown in Fig. 3C) are sufficient to generate a significantly higher average burst similarity than when considering all units and this difference gradually decreases when a larger percentile of the most correlated units are included (Supplementary Fig. 19A,C). When comparing the lower 20^th^ percentile of units (ranked by their average correlation), we observed a significant decrease of the average burst similarity in the population firing rate vector. When a larger percentile of the least correlated units is included, this difference gradually decreases (Supplementary Fig. 19B,C). Overall, we observed an increase in average burst similarity during the onset of the backbone units that subsequently plateaus for the remainder of their activation period across several organoids (Fig. 4E and Supplementary Fig. 19D for results across *n* = 8 organoids). These results further highlight that the activity of backbone units in organoids provide temporal stability across bursts at the ensemble level.

To quantify the trajectory of firing patterns generated during spontaneous activity in brain organoids we performed principal component analysis (PCA). We observed that the trajectory firing patterns follow conserved trajectories in PC-space that preserve timing relative to the burst peak (Fig. 4F, first two PCs are shown for organoid 1). When dividing the populations into backbone and non-rigid groups (Fig. 4F middle and right panel respectively), we observed that backbone units captured a larger variance for the first two PCs (73%) when compared to the non-rigid sub-population (25%) (Fig. 4G, Supplementary Figs. 20-21). These results highlight that a lower-dimensional subspace is occupied by neurons that fire in backbone ensembles, whereas the non-rigid population exhibit more irregular and, thus, higher-dimensional firing patterns and require more PCs to explain their variance. Of note, the lower-dimensional space is abolished after data randomization preserving both the mean neuron firing rate as well as the population firing rate (Supplementary Fig. 13). This indicates that the trajectories, and non-zero temporal correlation patterns are not a trivial result of the neuron’s mean firing rate.

### Uncovering temporal structure in population bursts with a hidden Markov model

We have previously shown that the distribution of functional connectivity in human brain organoids is well described by a heavy-tailed shape^30^, mirroring scaling rules found in cortical circuits such as the visual^5^ and somatosensory cortices^77^. The prevalence of these circuit motifs are widely believed to give rise to spontaneous activity patterns that spread across most of the cortical mantle, mirroring sensory-evoked responses observed *in vivo*^78^. To further investigate this rich repertoire of spontaneous neuronal firing patterns, we applied a hidden Markov model (HMM).

We trained an HMM to cluster single-unit spiking activity, generated by each brain organoid, into discrete states with shared firing patterns. The data was binned into 30 millisecond intervals (see Methods), reflecting timescales of fast electrophysiological dynamics in the cortex that span ∼10-50 milliseconds^56,79^. We found varying the HMM time bins over this range did not significantly impact the performance of the model (Supplementary Fig. 22). The identified HMM states, shaded by different colors and superimposed on spike raster plots (Fig. 5A), highlight their trajectories in relation to burst peak events. Here, each colored state represents a distinct linear combination of single-unit firing patterns that are coincident across the recorded ensemble. The first fifty firing rate realizations (across all units for a given state) are plotted as heatmap visualizations (Fig. 5B, left panels). Each representative state plot is accompanied by a histogram of the average firing rate per unit (Fig. 5B, right panels). These visualizations reveal the presence of distinct manifolds of firing patterns delineated by both differential gain and attenuation of single-unit firing rates associated with each state (Fig. 5B, red lines). Here a model with 20 hidden states was used for visualization purposes, but we observe similar-performing models across a range of hidden state counts (Supplementary Fig. 23). To further validate that the transition between states was not a result of trivial differences in the mean rate, we ran our HMM analysis after data randomization, preserving both the neuron’s mean firing rate and population rate^58,73^, and observed that the log likelihood was significantly larger for real data (Supplementary Fig. 24). These results demonstrate the presence of distinct combinations of firing patterns, regularly activated during burst events, captured by the HMM. The average change in firing rate (across all units) between the three states (Fig. 5B) further revealed that these discrete transitions represent both increases and decreases in relative firing rate across different sets of units and states. We next utilized the HMM to model transition probability between observed states in our brain organoids. Lower probability states were observed prior to the burst peak with a larger number of states sampled per unit time. However, during and immediately following the burst peak we observed a narrowing of states available per unit time (Fig. 5C). This narrowing effect might establish a sequential arrow of time in state space, where initial states preserve more precise timing relative to following states (Supplementary Fig. 25)^16^. Subsequently, as the burst fades, state observations return to a lower probability. The number of realized states observed during burst events remains similar over the variable range of hidden states (Supplementary Fig. 26), indicating the model’s robustness to discrete patterns of spiking activity and their evolution in time. Next, we show that the more temporally rigid units that reside within backbone sequences also distribute across a larger pool of HMM states compared to non-rigid units, which span a smaller range of states (Fig. 5D), a result consistent across organoids (Supplementary Fig. 27). These differences are further visualized by performing PCA on state vs. unit realizations (Fig. 5E), which suggests that backbone and nonrigid units may be linearly separable by hidden state structure. We quantified this by training linear support vector machine (SVM) classifiers (see Methods) to distinguish backbone units from non-rigid units by their state structure representation (Fig. 5F), and found an accuracy of 83.9% ± 12.0%, significantly higher than the 63.2% ± 10.4% achieved when classifying by firing rate alone. Together these results highlight spontaneous population burst events in brain organoids consisting of ensembles that link together in time to establish neuronal manifolds, which exhibit more complex hidden state dynamics in real than in shuffled data (Supplementary Fig. 28). These manifolds represent a latent multidimensional space that is composed of a temporally rigid and flexible subsets of units. These units form a subspace of states that follow probabilistic trajectories that are Markovian in time, namely future states depend on the system’s current state. Recent theoretical models and experimental observations have proposed that local pairwise correlations are a dominant driver of irreversibility within noisy logical computations that contribute to a local arrow of time, and generate an irreversible Markovian process independent of sensory input^80,81^.

**Figure 5.**
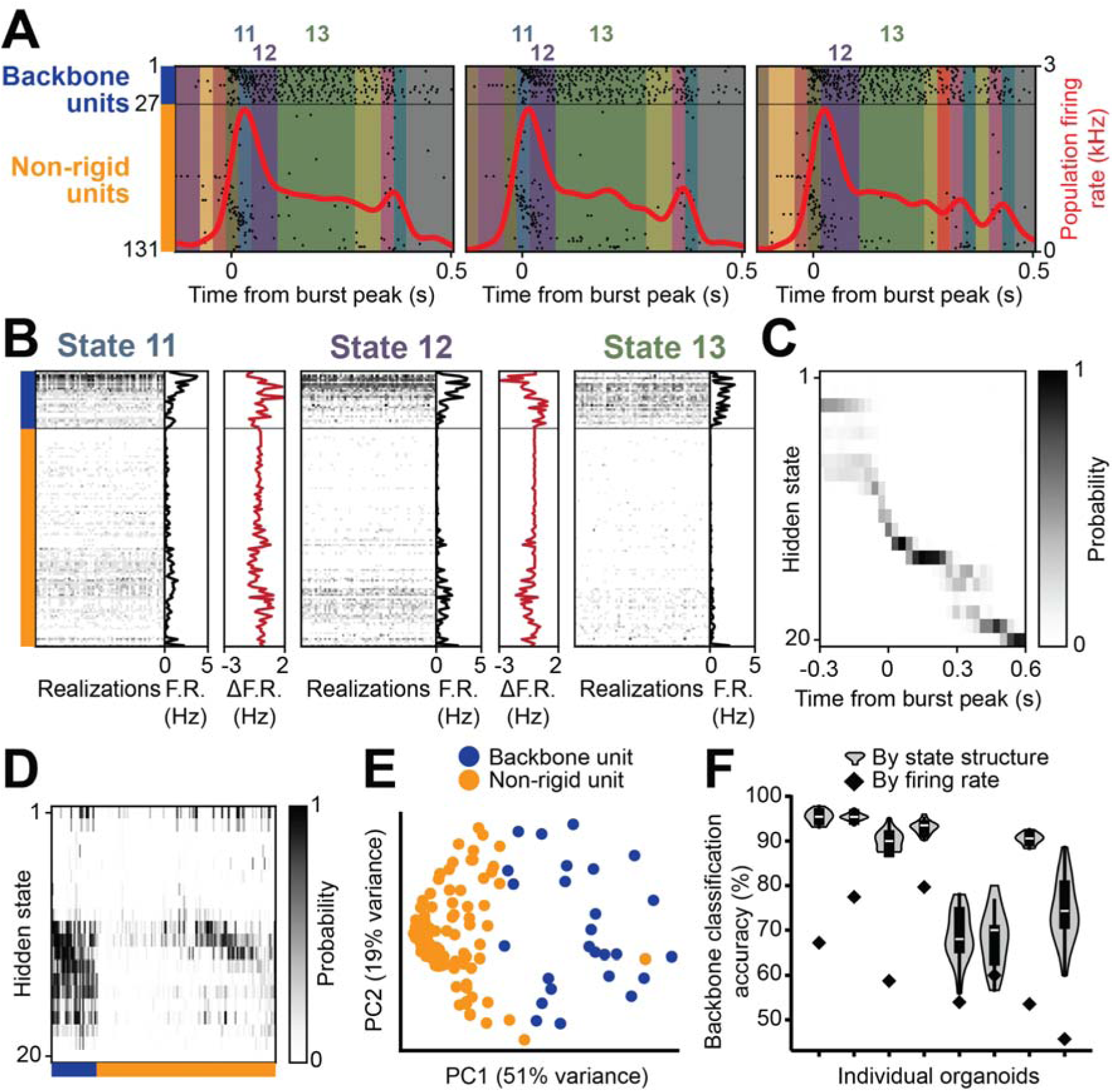
Hidden Markov Models (HMMs) explore stable trajectories and population motifs. (**A**) Repeated sequence of discrete hidden states during three example bursts. The model assigns each 30 ms bin of the spiking data a hidden state, indicated here by the background color. Note the stereotyped trajectory both in state space and in the population rate (red). Spiking events are displayed as a raster, with the 27 backbone units displayed above and the 104 non-rigid units displayed below a separator line. (**B**) Each state represents a stochastic but repeated pattern of activity across all units. Example realizations of three hidden states which occur during the burst trajectory are displayed as heatmaps. The differences between states are highlighted in subpanels which show the overall firing rate of each unit, FR, corresponding to the highlighting of part **A**. Also shown is the difference between these plots for adjacent states, ΔFR, in red. (**C**) The sequence of hidden states follows a stereotyped trajectory across each burst. The probability of each hidden state as a function of time relative to the burst peak is displayed as a heatmap. Note in particular that for the first 0.3 seconds from the burst peak, there is very little variation in the pattern of hidden states, whereas later in the bursts (as well as before the burst begins), the variability is significantly higher. (**D**) Backbone units fire consistently even outside of the burst peak states 11 and 12. The probability that each unit will fire in each of the hidden states of an example HMM is displayed as a heatmap. The backbone units (left) are active in various states. (**E**) Backbone and non-rigid units are almost linearly separable by consistency across states. First two principal components of vectors representing each unit as the sequence of its consistency across states. Backbone and non-rigid units are indicated with color. (**F**) Firing rate is not adequate to identify backbone and non-rigid units. The linear separability of backbone/non-rigid units based on their vectors of consistency scores is shown as a violin plot across all fitted HMMs for each brain organoid. Linear separability based only on the firing rate is marked by a diamond for comparison. In all cases, classification by state structure performs significantly better (Student’s *t*-test, *P* < 0.0001).

### Endogenous spiking sequences in early developing neonatal cortex

We next asked if spiking sequences also emerge in early developing cortical circuits. It has been established that sequential patterns are crucial components for mature brain function^16^ as well as early navigational tasks^24^, however, it is not known if such patterns are present in early brain development before eye opening and exploration occur. To address this open question, we investigated the emergence of sequential firing patterns in acute coronal slices obtained from the developing murine somatosensory cortex. We performed acute extracellular recordings using high-density CMOS MEAs (*n* = 6 slices). Discontinuous single-unit spiking activity alternated with long periods of quiescence and abrupt transitions to synchronized bursts (Fig. 6A and Supplementary Fig. 29 for plots from additional slices).

**Figure 6.**
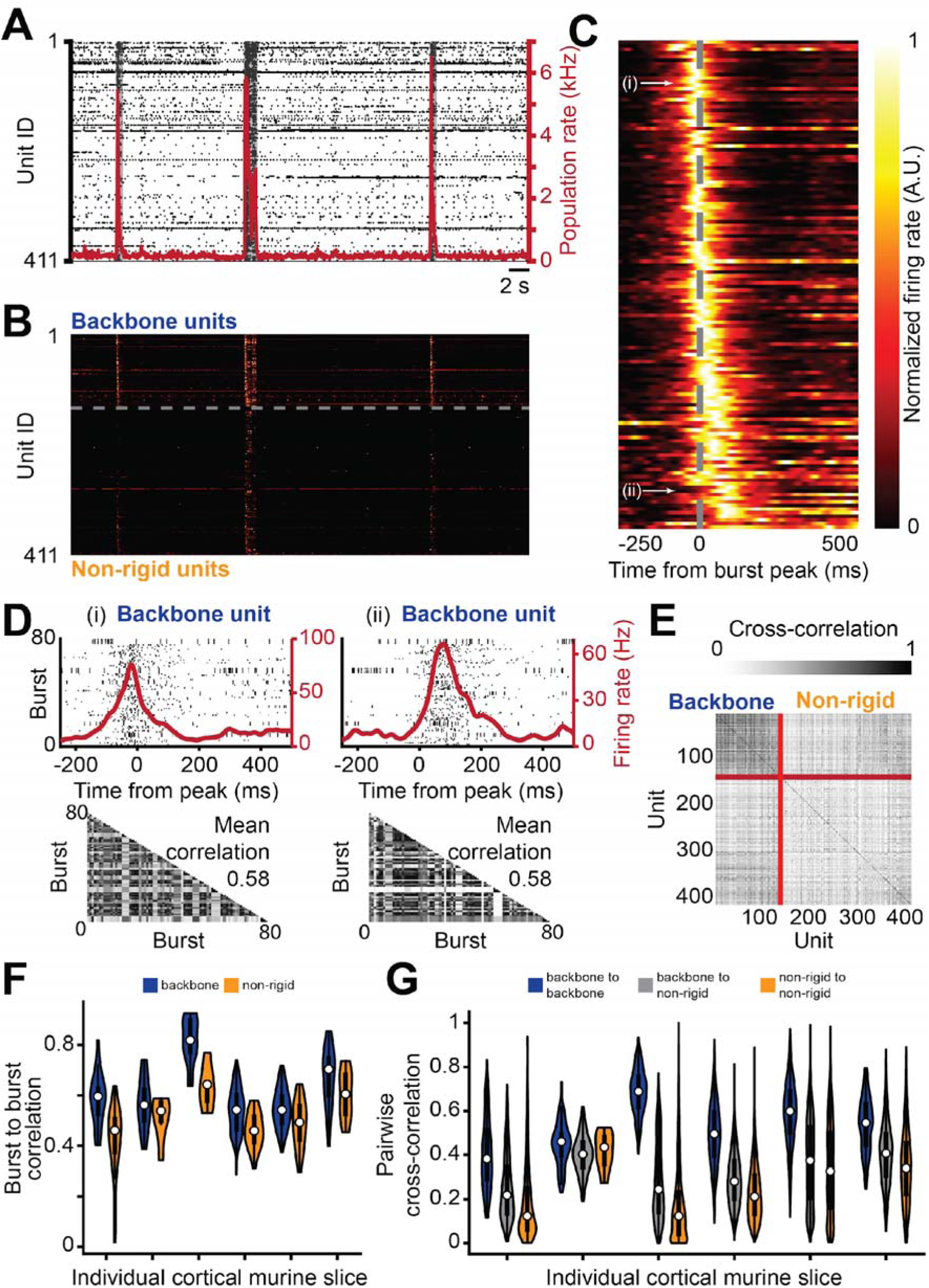
Recurring sequential activation patterns in murine neonatal cortical slices generate a stereotyped temporal backbone. (**A**) Raster plot visualization of single-unit spiking measured across the surface of the somatosensory cortex of a murine neonatal cortical slice dissected at P13, positioned on top of a microelectrode array. The population firing rate is shown by the red solid line. Population bursts are marked by sharp increases in the population rate. Burst peak events are denoted by local maximas that exceed 4x-RMS fluctuations in the population rate. The shaded gray regions denote the burst duration window as defined by the time interval in which the population rate remains above 10% of its peak value in the burst. (**B**) The instantaneous firing rate of single-unit activity from panel **A** after reordering. Backbone units are plotted above the dotted line and non-rigid units are plotted below the dotted line. In each category, units are ordered based on their median firing rate peak time relative to the burst peak, considered over all bursts in the recording. The backbone unit threshold for murine neonatal cortical slices was lowered to at least 2 spikes in at least 70% of all bursts. (**C**) The average burst peak centered firing rate measured across all burst events. The burst peak is indicated by the dotted line. The unit order is the same as **B**. Note the progressive increase in the firing rate peak time relative to the burst peak, as well as a spread in the active duration for units having their peak activity later in the burst. The average firing rate is normalized per unit to aid in visual clarity. (**D**) Top left, spike events are plotted for a regularly firing backbone unit over a fixed time window relative to population burst peaks. Each row represents spike events in a unique burst event. The average burst peak centered firing rate of the unit is plotted as the red solid line. Top right, another regular firing backbone unit is plotted with a larger temporal delay relative to the burst peak events. Bottom, heatmap visualizations showing the correlation coefficients of the burst peak centered firing rates of the unit shown in the top. The correlation is computed for each pair of bursts, using a maximum lag of 10 ms. The average over all burst pairs is denoted in the top right of the heatmap. Both example backbone units are marked in **C**. (**E**) Pairwise cross correlation coefficients over the instantaneous firing rates of all pairs of units with at least 30 spikes counted over all burst events. A maximum lag time of 50 ms was used. The solid red lines separate the backbone units and the non-rigid units. (**F**) The average burst-to-burst correlation per unit grouped in backbone and non-rigid categories for all murine neonatal cortical slices. Burst-to-burst correlations are significantly higher for backbone units compared to non-rigid units (*P* < 10^−20^, linear mixed-effects model). See Fig. 7B for the results of the statistical comparison of firing rate normalized data between different model types. (**G**) The pairwise correlations between all unit pairs for all murine neonatal cortical slices grouped into pairs that connect two backbone units (blue), pairs that connect a backbone unit and a non-rigid unit (gray) and pairs that connect two non-rigid units (yellow). Correlations between backbone units are significantly higher (*P* < 10^−20^, linear mixed-effects model) than the cross-correlations between pairs of backbone and non-rigid units and pairs of non-rigid units. See Fig. 7C for the results of the statistical comparison of firing rate normalized data between different model types.

Next, we quantified the consistency of firing patterns generated by coronal slices of the somatosensory cortex. First, we split single units into backbone and non-rigid units based on their firing recruitment within population bursts, similarly to the protocol used for brain organoid data (Fig. 6B, Supplementary Fig. 2,29, see Methods). The temporal pattern of these backbone units was apparent upon signal averaging of each unit’s instantaneous firing relative to the burst peak events (Fig. 6C), which revealed a sequential firing structure within the subset of units regularly recruited during spontaneous bursts events. Analysis of the single-unit spike times further revealed preservation of spike timing relative to burst peak events with consistent shifts between their peak firing rates in neonatal murine brain slices (Fig. 6D). Analogously to what we observed in brain organoids, we found that backbone units form a more strongly correlated core when compared to their non-rigid counterparts (Fig. 6E-G, Supplementary Fig. 14,18), and that activity in murine neonatal cortical slices generated temporal sequences that span ≈ 10^2^ millisecond time scales. Together, these results highlight that slices of the developing murine somatosensory cortex generate firing patterns composed of both rigid and flexible units that are capable of establishing sequential activation patterns commonly observed in mature cortical circuits across a range of species and brain regions^16^.

### Comparing sequences across neurodevelopmental models

To understand the potential role played by the three dimensionality of neurodevelopment, we investigated the firing patterns generated by two-dimensional murine primary cultures of neurons derived from the cortex, and compared them to what we observed in human brain organoids, murine brain organoids and acute murine neonatal cortical slices (Supplementary Fig. 16). Here, all four neuronal systems have characteristic population bursts with consistent firing units that are recruited during burst epochs (Supplementary Fig. 2). These consistent firing neurons have significantly higher average firing rates compared to the more irregular non-rigid counterparts (Fig. 7A, P < 10^−6^ for organoid, murine slice and primary cultures, linear mixed-effects model, Supplementary Fig. 30 for results per recording). However, on average the primary two-dimensional cultures tend to have higher average firing rate distributions for both the backbone and non-rigid units (*P* < 10^−10^ for comparisons across organoid and murine neonatal cortical slices). Meanwhile, we observed that backbone units for organoids and neonatal cortical slices have higher normalized burst-to-burst single-unit firing rate correlation scores relative to two-dimensional primary cultures (Fig. 7B, see Methods and Supplementary Fig. 14 for unnormalized and normalized results per recording). Furthermore, after normalizing with the randomized data, the average burst-to-burst single-unit firing rate correlations remain significantly larger for backbone units than for non-rigid units in both organoids and neonatal slices (*P* < 10^−10^, *P* = 7×10^−4^, respectively, linear mixed-effects model, backbone unit distribution mean is larger than non-rigid), whereas this difference is not significant for primary cultures (*P* = 0.06, linear mixed-effects model, backbone unit distribution mean is smaller than non-rigid). Despite the strong variability that can occur between organoids grown from different batches^82^, we found that the difference in normalized burst-to-burst correlations between backbone units and non-rigid units were consistent across human and murine organoids (Supplementary Fig. 3E, Supplementary Fig. 31, Supplementary Fig. 14, Supplementary Fig. 6, 7). These neurophysiological features were conserved across whole and sliced organoid models measured in different laboratories (Methods).

**Figure 7.**
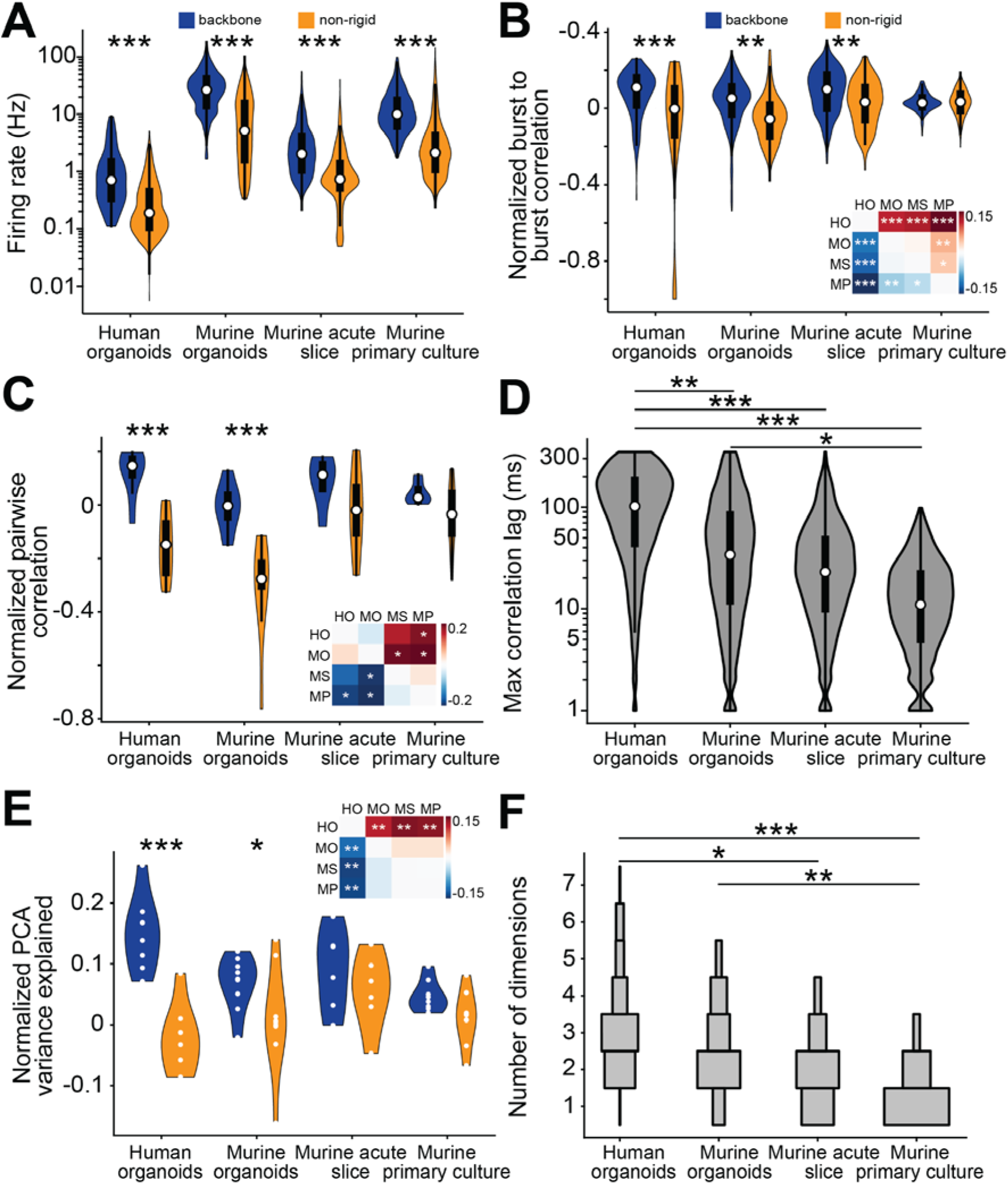
Backbone units provide a stable reference frame in brain organoids and murine neonatal cortical slices but not in murine primary cultures. (**A**) The log of the average firing rate per unit grouped in backbone and non-rigid categories for organoid slices, murine neonatal cortical slices and primary cultures. The human brain organoid group consists of 8 different organoids (4 whole organoids, 4 sliced organoids) with a total of 1048 units, 275 backbone and 773 non-rigid, the murine organoid group (dorsal forebrain identity) consists of 9 different whole organoid with a total of 1179 units, 603 backbone and 576 non-rigid, the murine neonatal slice group consists of 6 different slices from a total of 3 animals (2 unique slices per animal) with a total of 786 units, 296 backbone and 490 non-rigid, the murine primary group consists of 8 different cultures with a total of 1048 units, 277 backbone and 771 non-rigid. Statistical differences between the different model types and between backbone and non-rigid units for the different model types are computed using a linear mixed effects model. The model considers interactions between backbone and non-rigid units and between different model types. The specific recordings within each model type are included as grouping factors in the model. Each model is significantly different from each other (*P* < 10^−10^) and within each model the backbone and non-rigid units are significantly different from each other (*** = *P* <0.001, ** = *P* < 0.01, * = *P* < 0.05). (**B**) The normalized average burst-to-burst correlation per unit grouped the same as in **A**. Normalization was performed by subtracting the average burst-to-burst correlation per unit after shuffling from the original value and the dividing the outcome by the sum of both values (Original-Shuffled)/(Original+Shuffled). Similar to **A**, the statistical differences between the model types and between the backbone and non-rigid units were computed using a linear mixed effects model. Each model is significantly different from each other (*P* < 10^−10^) and for the organoids and murine cortical slices the backbone and non-rigid units are significantly different from each other (*** = *P* <0.001, ** = *P* < 0.01, * = *P* < 0.05). (**C**) The normalized pairwise cross-correlation per unit pair is grouped into backbone-to-backbone (blue), backbone to non-rigid (gray) and non-rigid to non-rigid (yellow) for each model type individually. Normalization was performed by subtracting the average pairwise correlation per unit pair after shuffling from the original value and then dividing the outcome by the sum of both values (Original-Shuffled)/(Original+Shuffled). A linear mixed effects model was used to assess which of the distributions were significantly larger than 0, indicating above random correlation strengths (*** = *P* <0.001, ** = *P* < 0.01, * = *P* < 0.05). Note that even though all three-unit pair types have a *P* value smaller than 10^−10^ for the organoids due to the large *n*, the effect size for backbone-to-backbone unit pairs is notably higher compared to the other two categories. (**D**) The absolute lag times that resulted in the optimal pairwise cross-correlations for pairs of backbone units shown in C. A linear mixed-effects model was used to assess differences between model types, reflecting that the optimal lag times are significantly larger for human and murine brain organoids compared to primary cultures (*** = *P* <0.001, ** = *P* < 0.01, * = *P* < 0.05). These long latency correlations result from consistent backbone sequences. (**E**) The normalized fraction of the variance explained summed over the first three principal components for the PCA manifolds per model type. To account for differences in the total number of principal components per category, the summed explained variance for the first three principal components is divided by the summed explained variance of the first *X* principal components, where *X* is the lowest number of total principal components from the three categories, all units, backbone and non-rigid. This value is computed for the original data and the shuffled data after which the shuffled result is subtracted from the original result to get the final value. The dots within the violins represent each individual recording. A linear mixed-effects model was used to assess differences between model types (*** = *P* <0.001, ** = *P* < 0.01, * = *P* < 0.05). (**F**) Hidden Markov models explore a higher-dimensional space for human and murine brain organoids than for primary cultures. For each fitted HMM, the dimensionality of representation was estimated by counting the number of principal components of the HMM observations that were required to explain at least 75% of variance. A linear-mixed effects model was used to assess differences between model types (*** = *P* <0.001, ** = *P* < 0.01, * = *P* < 0.05). No normalization with shuffled data was applied since shuffled models always required only 1 principal component. See Supplementary Fig. 32 for the results of other percentage thresholds.

We next focused our analysis on pairwise cross correlations between single-unit firing rates among the backbone and non-rigid units (Fig. 7C, Supplementary Fig. 18). We observed significant increase in the pairwise correlations between backbone-to-backbone (blue) units when referenced to randomized data preserving both population and mean single-unit firing rates^58,73^ (see Methods) for both the organoid and murine neonatal slice recordings (*P* < 10^−10^, linear mixed-effects model), an effect not observed in two-dimensional murine primary cortical cultures (*P* > 0.99, linear mixed-effects model). The differences in firing rate, burst-to-burst correlation and pairwise correlation were present in organoids at all observed developmental time points (Supplementary Fig. 5B-D). Furthermore, we found that higher normalized pairwise correlations between backbone unit pairs were consistently observed in both whole and slice recordings from unguided human brain organoids. These organoids were generated using different cell lines and different protocols in separate laboratories^30,31^ (Supplementary Fig. 3F). This trend was also present in mouse brain organoids of dorsal forebrain identity (Supplementary Fig. 8, 18). Interestingly, the temporal trajectory of backbone units for the two-dimensional primary cultures exhibited synchronous activity centered about the burst peak (Supplementary Fig. 16C), effectively abolishing sequential structure, and were identical to randomized organoid data (Supplementary Fig. 13C). As a result, both human and murine organoids showed significantly larger cross correlation lag times among backbone unit pairs compared to primary cultures (Fig. 7D, P < 0.001 and *P* < 0.05 respectively), which highlights their ability to sustain correlated neuronal activity with time constants that span order 10^2^ ms found in cortical and subcortical circuits with the capacity to support sequential firing patterns^16,56,63,70^.

Additionally, we performed principal component analysis (PCA) on the single-unit firing rates to quantify further the variance explained by the backbone and non-rigid single-units across these developmental models. The backbone units in the organoids explain a significantly higher fraction of the total variance than the other model types and unit types, as reflected by the percent variance explained by the first three principal components (Fig. 7E, Supplementary Fig. 21, *P* < 0.001, linear mixed-effects model). Together these findings highlight the finding that organoids generate sequential patterns that reside in a lower dimensional subspace (explained by fewer PCs) that is embedded in a higher dimensional background (requiring larger number PCs) of comparatively more irregular spiking patterns. Interestingly, we observe that backbone units are present in neonatal slices from the somatosensory cortex with the intrinsic capacity to generate sequences that span timescales (order 10^2^ milliseconds) observed in mature cortical circuits^13,14,26^, and occur with less stereotypy compared to organoids. Meanwhile, recurring sequential activation patterns of backbone units are not sustained in two-dimensional primary cultures.

We next asked if our analysis using hidden Markov models (HMMs) would enable us to quantify the firing patterns that are present in brain organoids, brain slices and two-dimensional primary cultures. First of all, we noted that in all models, the non-rigid units are predominantly Poisson, and backbone units are predominantly non-Poisson (Supplementary Fig. 33). In an *in vivo* setting, it has been well established that Poisson randomness is not a universal feature of spiking patterns in the cortex^57^, where architectonically defined brain regions generate homologous firing patterns that differ systematically across brain regions and less across species, a feature consistent from mice to cats and monkeys^83^. To further quantify the complexity across the different neuronal model systems we calculated their dimensionality, which we defined as the number of principal components of the HMM observations required to explain a variable fraction, θ, of the total variance. Here, the hidden states range from 10-30 and θ spans the range from 0.5 to 1, (see Methods and Supplementary Fig. 32). At θ = 0.75, human organoids are separable from slice data (*P* = 0.04 according to a linear mixed-effects model with Poisson observations); however, two-dimensional primary cultures exhibit substantially lower-dimensional HMM observations than both human (*P* < 0.0001) and murine (*P* = 0.003) brain organoids (Fig. 7F), with similar trends existing for a range of values of θ (Supplementary Fig. 32). Importantly, we observed that within organoids, hidden states correspond to clusters of population activity patterns which are distinguished across multiple dimensions, whereas after randomization, the HMM captures only a one-dimensional space which scales with the population rate (Supplementary Fig. 28). Furthermore, the number of realized states does not strongly depend on the number of hidden states (Supplementary Fig. 26). However, the rate at which states are traversed per unit time differs significantly across models (Supplementary Fig. 34), with the three-dimensional models exhibiting a lower state traversal rate than the two-dimensional primary cultures.

### Firing pattern stability

Despite the similarities in neuronal firing patterns and the establishment of stable sequences observed in both organoids and neonatal slices, two-dimensional counterparts are dominated by Poisson-like irregularity, which precludes the generation of sustained temporal patterns and confines their state transitions, as defined by a HMM, to a lower-dimensional space (Fig. 7F). However, all models can generate complex neuronal firing patterns. Many complex systems, including brains across phylogeny, show signs of criticality^61^. Criticality is a dynamical state that comprises internally-generated multi-scale, marginally-stable dynamics that maximize general features of information processing, such as dynamic range, information transmission, susceptibility, and robustness^84^. Criticality directly accounts for a system’s capacity for complex function, and is a homeostatic endpoint in the brain^60^ that is, in the intact brain, maintained by sleep^85^. While measurement of criticality traditionally requires extended sampling of a system’s activity, recent progress solves this problem by applying renormalization group theory^86^ - to the temporal features of neural data. We assessed each system’s proximity to criticality using temporal renormalization group theory to quantify how close neural dynamics are to exhibiting perfect temporal scale-invariance^62^. At criticality, general features beneficial to complex computation are simultaneously optimized, including information transmission, information storage, dynamic range, entropy, and susceptibility^60,84^. The distance metric *d_2_* measures how far a system’s dynamics are from criticality, with values between 0.0-0.1 indicating near-critical dynamics that span many timescales without a characteristic scale, while larger values indicate deviation from criticality. We found that a subset of each preparation type generated activity well-described by the autoregressive model framework and exhibited near-critical dynamics (see Supplemental Fig. 35 and Methods for additional details). Shuffling spike times consistently abolished temporal criticality across all preparations, producing significantly larger *d_2_* values (*P* < 10^−10^ compared to intact data). Murine organoid preparations showed particularly good agreement between empirical measurements and model predictions. Taken together, these data suggest that all preparations are capable of generating near-critical dynamics when properly captured by the autoregressive framework. This indicates that the capacity for scale-invariant dynamics may be a fundamental property of neural circuits that emerges during development, independent of precise circuit architecture or environmental context.

## Discussion

The advent of high-density extracellular recordings has facilitated the detection of non-random firing patterns that assemble into temporally precise sequences that are believed to form a basis for broadcasting and computing information in the brain^16^. Whether such sequences are emergent features, present during early brain development, a stage that is dominated by spontaneous activity with the potential to encode informational content, remains unclear. It has been hypothesized that sequences are ‘preconfigured’ and represent an innate architecture independent of external experience^29^. However, experimental evidence in support of this notion is still sparse, largely due to experimental inaccessibility^63^.

To address this open question, we leveraged state-of-the-art high density extracellular recordings from three-dimensional stem cell derived models of the human brain, known as brain organoids, which represent an intrinsically self-organized neuronal system that recapitulates key facets of early brain development^34–37^ and the establishment of functional circuits^30,39,42^. Crucially, brain organoids are not exposed to sensory information, and are thus an ideal model to study whether the emergence of sequences is truly experience-independent. Here, our analysis of single-unit firing patterns within human and murine brain organoids revealed the presence of a subset of high-firing rate neurons capable of generating firing patterns that assemble into temporally precise spiking sequences. A larger subpopulation conversely exhibits lower firing rates and less regular firing patterns (Fig. 2), where temporally rigid sequences project back onto a minority of strong functional connections (Fig. 3), which we previously reported are emergent properties of brain organoids^30^. In a seminal paper, Hopfield demonstrated that emergent computational properties from simple properties of many cells, rather than complex circuits, are capable of generalization, time sequence retention, error correction and time-evolution of the state of the system^87^. Therefore, it’s not surprising that the temporal structure of spontaneous and evoked patterns of cortical circuits are similar^88,89^, since such representations are drawn from a functionally connected neuronal pool with right-skewed, lognormal scaling rules^5^.

Units that fire within the recurrent neuronal firing sequences exhibit varying degrees of temporal precision. The subpopulation of neurons that fire at the beginning of the population bursts are the most constrained, whereas units firing later in the sequences are more temporally flexible (Fig. 2F,G). An analogous organization of spiking activity is present also in the somatosensory and auditory cortex of adult rats^65^, and in the three-layered turtle cortex^14^. In the hippocampus, experience-dependent replay has been shown to emerge from spontaneous, experience-independent preconfigured sequences^24^. The balance between temporally correlated and irregular spiking neuronal populations is an important feature of information processing in the brain. For example, large-scale extracellular field recordings from neurons in the mouse visual cortex and monkey brain have revealed a low-dimensional subspace of neurons that are entrained to population firing dynamics and represent a fixed attribute insensitive to external stimulus^58^. Consistent with this observation, also in brain organoids, we observe a low-dimensional subspace, consisting of backbone units, that resides within a higher dimensional space spanned by more weakly correlated and irregularly firing units (Fig. 4). Moreover, we show that the stochastic firing patterns of randomized organoid data closely mirror the firing patterns of two-dimensional dissociated cortical cultures, which exhibit culture-wide synchronization events that cannot sustain sequential activity patterns (Supplementary Fig. 13,16). Such an effect is likely the result of highly redundant and interconnected two-dimensional network configurations commonly observed *in vitro*^74,76^.

Together, these findings highlight that the self-organized generative process of neurogenesis and synaptogenesis within brain organoids may serve as a faithful *in vitro* model of the human brain, and do not simply generate random networks. Our results support the hypothesis that construction of complex networks capable of recapitulating *in vivo* neural dynamics requires morphogenesis in three-dimensions. In fact, recent experiments have demonstrated that functional circuits in human brain organoids, when interfaced with a machine, can function as a reservoir for computing, capable of speech recognition and nonlinear equation prediction^90^.

In an *in vivo* setting, a minority pool of strongly correlated neurons have been proposed to serve as a fast-acting system that resides within a more weakly coupled background, believed to function as a pre-configured and optimally tuned brain state^11,91^. The delicate balance between rigid and flexible spiking components and the establishment of sequential spiking patterns are an emergent feature of the three-dimensional cytoarchitecture, which is further supported by our analysis of spontaneous activity from neonatal slices of the mouse somatosensory cortex (Fig. 6). Recent work has further established that a neurophysiological backbone is likely organized during neurogenesis where common pyramidal neuronal progenitors establish high firing rate subnetworks, which are functionally connected and co-activated across brain states^29^. The presented organoid data shows that highly correlated backbone units increase in burst-to-burst correlation from earlier organoid developmental time points relative to later time points (Supplementary Fig. 5), a feature conserved across human and murine brain organoids. This transition also coincides with the functional incorporation of inhibitory interneurons in our brain organoids^30,92^, and reflects an excitatory-inhibitory balance shifting towards inhibition that is observed in *in vivo* cortical circuits^49^, while preserving right-skewed functional organizational rules^93^. Further, we demonstrate that the temporal rigidity of neuronal sequences is strongly modulated by inhibition of GABAergic input (Supplementary Fig. 11), which leads to a sharp increase in burst synchronization - an effect consistent with the inhibitory role of GABA. This finding highlights the critical role of interneuron signaling in maintaining network homeostasis and microcircuit function observed in mature brain circuits^94,95^, suggesting that it may also play a key role in shaping early neural dynamics prior to sensory input. While previous studies have suggested that GABA exerts a paradoxical excitatory effect during early brain development, with a transition to inhibition occurring after postnatal week two in the murine neocortex^96,97^, a growing body of evidence points to inhibitory effects of GABAergic signaling occurring much earlier in development^49,98–100^. Our findings align with this emerging perspective, indicating that interneurons and their inhibitory effects are key contributors to the temporal structure of neuronal sequences during the initial stages of brain development. The brain organoid model thus provides a unique platform for future research to investigate how interactions between brain regions and specific cell types drive the assembly of functional microcircuits^101^.

Brain organoids represent a self-organized neurodevelopmental system that operates as a truly closed system devoid of external input, and yet is capable of generating a rich repertoire of activity patterns that resemble the temporal dynamics of the early developing brain. Embedded within these firing patterns we observed an overlap between the pool of non-Poisson-like firing patterns generated by brain organoids and neonatal brain slices. However, within primary cortical cultures, with an inherently randomized cytoarchitecture, we observed a shift to lower dimensional state transitions as captured by a hidden Markov model (Fig. 7F). In an *in vivo* setting, brain regions are defined by a balance between a repertoire of firing patterns that span irregular Poisson dynamics to clock-like regularity, which depend on local circuit architectures and are believed to be critical components underlying higher order brain function^57^. In fact, the ‘backbone’ of consistently firing neurons, which form a minority pool within strongly connected networks, can predict as much as 80% accuracy during motor control in humans^102^. Backbone units have been proposed to function as an ‘ansatz’, or initial estimate for matching behavior to external environmental inputs^11^. We posit that the highly correlated, non-Poisson neuronal components may serve as basis for the emergence of temporal sequences in early brain development. Early sequences might function as an internal reference frame for larger scale population dynamics found in mature circuits^16^, which will later be calibrated through interplay between sensory input and motor output^63^. In fact, recent work in the postnatal murine brain has revealed that preconfigured sequence motifs (spontaneously generated in sleep and rest states) emerge before explicit sequential experience and are not improved by sequential experience during the third postnatal week after navigational beyond the nest^24^. We further demonstrate that neuronal firing patterns within brain organoids and neonatal brain slices generate strong non-random pairwise correlations with varying degrees of temporal jitter (relative to population events) with non-zero phase lags. These ensembles link together to establish a manifold of trajectories that are identifiable using a hidden Markov model, with a core of these probabilistic state transitions consisting of units with strong pairwise interactions (Fig. 5C,D). These subsets of units may function as an ‘irreversible’ set of ‘noisy logic elements’ that define a local arrow of time, which has recently shown to emerge in retinal circuits, where neuronal activity remains irreversible even when their inputs are not^80^.

In summary, our analysis of spontaneous activity generated by stem cell derived human and murine brain organoids, demonstrates that structured spiking sequences can emerge when completely devoid of sensory experience and motor output, supporting the pre-configured brain hypothesis. These results are in line with recent work showing how pharmacologically abolishing all central nervous system activity during the development of the larval zebrafish did not alter an oculomotor behavior^103^, suggesting that complex sensory-motor systems are hard-wired by activity-independent mechanisms. Our findings recall the philosophy of Immanuel Kant in *Critique of Pure Reason*^104^, who posited an *a priori* construction of a space-time map that in modern terms, could serve as a ‘scaffold’ to enable the brain to interact with and make sense of the world. In conclusion, brain organoids provide a heuristic platform for exploring how exogenous inputs refine self-organized neuronal circuits imbued with the innate capacity to process information and compute^90^, while also facilitating new studies into the genetic mechanisms governing the assembly of functional circuitry during early human brain development^105–109^.

## Methods

### Human brain organoid slice recordings and pre-processing

The human brain organoids presented in Fig. 1-4 were grown and prepared for extracellular field recordings as described in Sharf *et al*.^30^. Organoids with less than 20 active units were not considered as significant to reliably separate a backbone and non-rigid population. Briefly, brain organoids were grown based on methods developed by Lancaster *et al*.^35^ and were of predominant forebrain identity based on single cell RNA sequencing analysis^30^. The recordings were made using complementary metal-oxide-semiconductor (CMOS) micro-electrode array (MEA) technology (MaxOne, Maxwell Biosystems, Zurich, Switzerland). The arrays contain 26,400 recording electrodes of which a subset of 1,024 electrodes can be selected for simultaneous recording^64^. With a diameter of 7.5 µm each and a 17.5 µm pitch, the electrodes cover a total sensing area of 3.85 mm x 2.1 mm. The electrode selection was made based on automatic activity scans (tiled blocks of 1,020 electrodes) to identify the spatial distribution of electrical activity across the surface of the organoid. The 1,020 most active electrodes were chosen with a minimum spacing distance of at least two electrodes (2 x 17.5 µm), providing sufficient electrode redundancy per neuron to enable accurate identification of single units by spike sorting^110^, while simultaneously sampling network activity across the whole organoid surface interfacing the MEA. Measurements were made in a culture incubator (5% CO_2_ at 37 °C) with a sampling rate of 20 kHz for all recordings and saved in HDF5 file format. The raw extracellular recordings were band-pass filtered between 300-6000 Hz and subsequently spike sorted using the Kilosort2 algorithm^111^ through a custom Python pipeline. The spike sorting output was then further curated by removing units with an ISI violation threshold^112^ above 0.3, an average firing rate below 0.05 Hz and/or a signal to noise ratio (SNR) below 5.

### Whole human brain organoid recordings

Additional recordings were obtained from whole human brain organoids (Fig. 7), which were grown from human iPSCs and differentiated into organoids based on a previously established protocol^113^. Briefly, the NIBSC8 iPSC line was cultured in mTESR Plus medium on vitronectin. Neural differentiation was induced with Neural Induction medium via SMAD inhibition (Gibco) and organoids were differentiated under gyratory shaking (88 rpm, 50 mm orbit) for up to 8 weeks in Neurobasal Plus medium supplemented with 1x B27-Plus, 10 ng/ml human recombinant GDNF (GeminiBio™), 10 ng/ml human recombinant BDNF (GeminiBio™), 1% Pen/Strep/Glutamine (Gibco, Thermo Fisher Scientific). Half changes of medium were performed 3-times a week.

Organoids electrophysiology data was sourced from Alam El Din *et al*.^31^ which made use of high-density MEAs integrated into 6-well configuration (MaxTwo, Maxwell Biosystems, Zurich, Switzerland) with the same electrode configuration per well as the MaxOne system described in the previous section. Briefly, whole organoids attached to MEAs at 9.5 weeks old and grown for 32 days. Prior to plating, the HD-MEA chips were coated with 0.07% Poly(ethyleneimine) solution (Sigma Aldrich) diluted in a 1x borate buffer (Thermofisher) for 1 hour at 37°C and 5% CO_2_. Following three washes with water to remove the Poly(ethyleneimine) solution the chips were dried for an hour in the hood. 0.04 mg/ml Mouse Laminin (Sigma Aldrich) was then added and incubated for 1 hour at 37°C and 5% CO_2_. After the incubation, the laminin was removed, and the organoids were plated on the MEA. Recordings were performed using the same methods in the previous section.

### Bulk RNA sequencing of brain organoids

Total RNA was isolated from 8-week old organoids via a Quick-RNA™ Microprep Kit (Zymo Research). The RNA integrity was assessed using the Agilent TapeStation 4200 (Agilent Technologies). A total of 500 ng of RNA was employed to generate mRNA libraries following the TruSeq Stranded mRNA Library Prep Kit (Illumina) protocol. This procedure involved isolating Poly-A mRNA, fragmenting it, synthesizing the first and second cDNA strands, adding adenylated 3’ ends, and ligating TruSeq RNA Combinatorial Dual Indexes (Illumina). The ligated fragments were then amplified for 15 cycles, purified, and quantified using the Qubit 3.0 Fluorometer (Thermo Fisher Scientific). The size and quality of the libraries were evaluated with the Agilent TapeStation 4200. Finally, the libraries were normalized, pooled, and sequenced on the NovaSeq X Plus (Illumina) using 150 bp paired-end reads, achieving approximately 100 million reads per sample.

### Bulk RNA sequencing analysis

Raw FASTQ files were processed using the nf-core RNA-seq pipeline which leverages Nextflow workflows (version 23.10.1) (https://doi.org/10.5281/zenodo.1400710114). Adapters and poor quality sequences were trimmed using “Trim-Galore!”^115^. Reads were aligned to the GRCh38 genome using ENSEMBL gene annotations (release 111) using STAR and transcripts were quantified with Salmon^116–118^. Post processing of reads was accomplished using SAMtools (sorting and indexing alignments)^119^, picard (duplicate identification)^120^, and BEDtools (genome coverage assessment)^121^. Read and mapping quality were assessed using RSeQC^122^, Qualimap^123^, dupRadar^124^, and Preseq^125^. Gene length corrected and scaled counts from Salmon were normalized using limma voom^126^. RNA-seq data from the four samples are deposited in the NCBI Gene Expression Omnibus.

### Immunohistochemistry and microscopy of human brain organoids

Organoid samples were stained following methods described in Romero *et al*.^113^ Briefly, after fixation in 2% PFA, organoids were permeabilized with 0.1% Triton X and blocked with 100% BlockAid^TM^. Organoids were stained with primary antibodies (Table below) at 4°C on a shaker for 48 hours followed by staining with secondary antibodies for another 24 hours.

Nuclei were stained with Hoechst 33342 trihydrochloride (Invitrogen Molecular Probes) at a concentration of 1:10,000. <u>Organoids were imaged on a FV3000RS Olympus or a Zeiss</u> <u>LSM800 confocal microscope.</u>

| Antibody | Host | Clone | Company | Cat # | Dilution |
| --- | --- | --- | --- | --- | --- |
| Gephyrin | Mouse | Monoclonal | Synaptic Systems | 147 011 | 1:1000 |
| MAP2 | Chicken | Polyclonal | Invitrogen | PA1-10005 | 1:5000 |
| GFAP | Rabbit | Polyclonal | Dako | Z0334 | 1:400 |
| O4 | mouse | Monoclonal | R&D | MAB1326 | 1:200 |

### Mouse ESC maintenance

Three mouse embryonic stem cell lines were used: C57BL/6, E14TG2a (129/Ola), and KH2 (129/SvJ × C57BL/6 hybrid). Mycoplasma testing using MycoAlert (Lonza # LT07-318) confirmed lack of contamination.

ESCs were maintained on recombinant human protein vitronectin (Thermo Fisher Scientific A14700) coated plates using mESC maintenance media containing Glasgow Minimum Essential Medium (Thermo Fisher Scientific 11710035), embryonic stem cell-qualified fetal bovine serum (Thermo Fisher Scientific 10439001), 0.1 mM MEM nonessential amino acids (Thermo Fisher Scientific 11140050), 1 mM sodium pyruvate (Millipore Sigma S8636), 2 mM glutamax supplement (Thermo Fisher Scientific 35050061), 0.1 mM 2-mercaptoethanol (Millipore Sigma M3148), and 0.05 mg/ml primocin (Invitrogen ant-pm-05). mESC maintenance media were supplemented with 1000 units/ml of recombinant mouse leukemia inhibitory factor (Millipore Sigma ESG1107). Media was changed every day.

Vitronectin coating was incubated for 15 min at a concentration of 0.5 μg/ml dissolved in 1× PBS pH 7.4 (Thermo Fisher Scientific 70011044). Dissociation and cell passages were done using ReLeSR passaging reagent (Stem Cell Technologies 05872) according to the manufacturer’s instructions. Cell freezing was done in mFreSR cryopreservation medium (Stem Cell Technologies 05855) according to the manufacturer’s instructions.

### Mouse embryonic stem cell derived cortical organoids

Organoids were generated from three distinct mouse embryonic stem cell lines: C57BL/6, E14TG2a (129/Ola), and KH2 (129/SvJ x C57BL/6 hybrid). Embryonic stem cells (ESCs) were dissociated into single cells using TrypLE Express Enzyme (Thermo Fisher Scientific, #12604021) for 5 minutes at 37°C. After dissociation, the cells were re-aggregated in lipidure-coated 96-well V-bottom plates at a density of 3,000 cells per well in 150 µL of mESC maintenance medium, supplemented with 10 µM Rho Kinase Inhibitor (Y-27632, Tocris #1254) and 1,000 units/mL Recombinant Mouse Leukemia Inhibitory Factor (Millipore Sigma, #ESG1107). Following 24 hours of re-aggregation, the medium was replaced with cortical differentiation medium, composed of DMEM/F12 with GlutaMAX (Thermo Fisher Scientific, #10565018), 10% Knockout Serum Replacement (Thermo Fisher Scientific, #10828028), 0.1 mM MEM Non-Essential Amino Acids (Thermo Fisher Scientific, #11140050), 1 mM Sodium Pyruvate (Millipore Sigma, #S8636), 1X N-2 Supplement (Thermo Fisher Scientific, #17502048), 2X B-27 minus Vitamin A (Thermo Fisher Scientific, #12587010), 0.1 mM 2-Mercaptoethanol (Millipore Sigma, #M3148), and 0.05 mg/mL Primocin (Invitrogen, #ant-pm-05). The medium was further supplemented with 10 µM Rho Kinase Inhibitor (Y-27632), 5 µM WNT inhibitor (XAV939, StemCell Technologies #100-1052), and 5 µM TGF-β inhibitor (SB431542, Tocris #1614).

Daily medium changes were performed, with N-2 and B-27 supplements added post-filtration to preserve their hydrophobic components. On Day 5, organoids were transferred to ultra-low adhesion plates (Millipore Sigma, #CLS3471), where the medium was replaced with fresh neuronal differentiation medium. The plates were then placed on an orbital shaker set to 68 rpm to prevent organoid fusion.

From Day 6 to Day 12, the neuronal differentiation medium consisted of Neurobasal-A (Thermo Fisher Scientific, #10565018), BrainPhys Neuronal Medium (Stem Cell Technologies, #05790), 1X B-27 Supplement without Vitamin A (Thermo Fisher Scientific, #12587010), 1X N-2 Supplement (Thermo Fisher Scientific, #17502048), 0.1 mM MEM Non-Essential Amino Acids (Thermo Fisher Scientific, #11140050), and 0.05 mg/mL Primocin (Invitrogen, #ant-pm-05), with the fresh addition of 200 µM Ascorbic Acid. Organoids were cultured under 5% CO2, with medium changes performed every 2-3 days. From Day 15 onward, the organoids were maintained in a neural differentiation medium containing BrainPhys Neuronal Medium, 1X B-27 Plus Supplement (Thermo Fisher Scientific, #A3582801), 1X N-2 Supplement, 1X Chemically Defined Lipid Concentrate (Thermo Fisher Scientific, #11905031), 5µg/mL Heparin (Sigma Aldrich, #H3149), and 0.05 mg/mL Primocin. Additionally, 200 µM Ascorbic Acid was included until Day 25. The medium was refreshed every 2-3 days, and the shaker speed was maintained at 68 rpm to minimize organoid fusion. To ensure optimal growth conditions, 16 organoids per well were consistently maintained.

### Single-cell RNA sequencing of cortical organoids

#### PIPseq library preparation

Murine cortical organoid library prep was carried out following the methods described in Clarke *et al*.^127^. Ten murine cortical organoids per cell line (3 cell lines defined above) were pooled together and dissociated using Worthington Papain Dissociation System (Worthington # LK003150) according to the manufacturer’s instructions. Briefly, 20 units of papain per ml, 1mM L-cysteine and 0.5mM EDTA were resuspended in Earle’s Balanced Salt Solution (EBSS). The enzyme solution was activated by incubating for 30 minutes at 37° C. After activation, we included 200 units of DNase I per ml. We transferred the tissue into the papain and DNase I solution and incubated for 30 minutes at 37° C. Every 10 minutes the samples were agitated by gently shaking the tube. The tissue was mechanically dissociated using flamed glass Pasteur Pipets (Fisher Scientific # 13-678-6B). The tissue was centrifuged at 300 RCF for 3 minutes. The supernatant was removed and approximately 1mL of 1X PBS containing 0.1% Bovine Serum Albumin (Millipore Sigma # A3311) was added and the cells were resuspended, then put through a 40 µm cell strainer (Corning # 431750) into a 6 well ultra low-adhesion plate (Millipore Sigma # CLS3471). Cells were then manually counted on a hemocytometer and 3.3K cells from each of the three dissociated organoid cell lines were pooled to make a total of 10K cells. These pooled cells were then combined into a single sequencing reaction using the PiPseq T2 kit and carried out according to the manufacturer’s instructions.

#### PIPseq data analysis

Samples were pooled and sequenced on the AVITI PE75 Flowcell ∼900M reads at the UCDavis Technologies Core. The sequencing data was then processed using the PIPseeker pipeline (PIPseeker v3.3, Fluent BioSciences), using the mouse genome GRCm39 (GENCODE vM29 2022.04, Ensembl 106)^128^. We used the default parameters to process the FASTQ files, perform mapping, transcript counting and cell calling.

Sensitivity 5 matrices were used for downstream analysis using R package Seurat (version 5.1.0)^129^. Souporcell was used for genotype demultiplexing and doublet identification^130^. Doublets were also identified using the R package DoubletFinder v2.0.4^131^.

Cells were filtered based on doublet identification, with greater than 20% mitochondrial genes, less than the 5th percentile unique genes detected, and greater than 50,000 total RNA counts. The raw gene count matrices were normalized with SCTransform function while regressing out mitochondrial genes identified the top 3,000 highly variable genes. Principal component analysis was performed and clusters were identified by the FindNeighbors function using 40 principal components (chosen based on visual inspection of the elbow plot). The function FindClusters was run with resolutions of 2, 1, and 0.5 with leiden clustering. The clusters were then embedded and visualized with Uniform Manifold Approximation and Projection (UMAP)^132^ and the resolution of 0.5 was chosen based on visual inspection of marker genes to represent the best approximal cell type. Annotation of the dataset was performed by marker gene expression from the FindMarkers function by the Wilcoxon Rank Sum test and reference to canonical marker genes from the Allen Brain Atlas Whole Cortex and Hippocampus^133^ visualized using the UCSC Cell Browser^134^.

#### Immunohistochemistry and microscopy

Organoids were collected and fixed in room temperature 4% Paraformaldehyde (PFA) (Thermo Fisher Scientific # 28908) and cryopreserved in 30% Sucrose (Millipore Sigma # S8501). They were then embedded in a solution containing 50% of Tissue-Tek O.C.T. Compound (Sakura # 4583) and 50% of 30% sucrose dissolved in 1X Phosphate-buffered saline (PBS) pH 7.4 (Thermo Fisher Scientific # 70011044). Organoids were then sectioned to 20 µm using a cryostat (Leica Biosystems # CM3050) directly onto glass slides. After 2 washes of 5 minutes in 1X PBS the sections and 1 wash in deionized water (Chem world #CW-DW-2G) sections were incubated in blocking solution 5% v/v donkey serum (Millipore Sigma # D9663), and 0.1% Triton X-100 (Millipore Sigma # X100) for 1 hour. The sections were then incubated in primary antibodies overnight at 4° C. They were then washed 3 times for 10 minutes in PBS and incubated in secondary antibodies diluted in blocking solution for 90 minutes at room temperature. They were then washed 3 times for 10 minutes in PBS and coverslipped with Fluoromount-G Mounting Medium (Thermo Fisher Scientific # 00-4958-02). Additional histology was performed on sections on fixed tissue that were not cryopreserved (Supplementary Fig. 10). Organoid samples were fixed in 4% paraformaldehyde (Thermo Fisher Scientific # 28908) overnight at 4 °C. After rinsing, samples were embedded in 4% low melting point agarose (invitrogen # 16520-050) in PBS and sectioned (50 µm) using a VT1000s vibratome (Leica, Lumberton, NJ). Sections were then incubated in an initial blocking solution consisting of 5% v/v donkey serum (Millipore Sigma # D9663), BSA 1% (Millipore Sigma # A7906), and 0.5% Triton X-100 (Millipore Sigma # X100) for 1 hour in 4°C. The initial blocking solution was then carefully removed, and sections were then placed in an antibody blocking solution with primary antibodies composed of 2% v/v donkey serum (Millipore Sigma # D9663), and 0.1% Triton X-100 (Millipore Sigma # X100) overnight. Sections were removed from the antibody blocking solution and were washed four times in 1X PBS for 15 minutes at room temperature. Sections were then placed with secondary antibodies diluted in antibody blocking solution, as previously described, at room temperature for 30 minutes. Another wash in 1X PBS was performed, followed by Hoechst 33342 nucleic acid counterstain for 15 minutes at room temperature. Three final washes were conducted and sections were mounted on glass slides using Fluoromount-G Mounting Medium (Fisher Scientific OB100-01).

Primary antibodies used were: rabbit anti-Map2 (Proteintech # 17490-1-AP, 1:2000); mouse anti-Pax6 (BD Biosciences # 561462, 1:100); rabbit anti-Nkx2.1 (Abcam # ab76013, 1:400); rat anti-Ctip2 (Abcam # ab18465, 1:250); rabbit anti-Brn2 (Thermofisher # PA530124, 1:400); anti-Gaba (Thermo Fisher Scientific # PA5-32241, 1:375). Secondary antibodies were of the Alexa series (Thermo Fisher Scientific), used at a concentration of 1:750. Nuclear counterstain was performed using 1.0 µg/ml Hoechst 33342 (Thermofisher # H1399).

Imaging was done using an inverted confocal microscope (Zeiss 880) and Zen Blue software (Zeiss). Images were processed using Zen Black (Zeiss) and ImageJ software (NIH).

#### Synaptic Blocker experiments

Preparation of stock solutions to block components of slow and fast synaptic transmission: the AMPA receptor blocker NBQX (Abcam) was solubilized in DMSO, the NMDA receptor blocker R-CPP (Abcam) and the GABA receptor blocker Gabazine (Abcam) were prepared in ultrapure distilled water (Life Tech) at 1000x the desired working concentration. The working concentrations were 10, 20 and 10 μM for NBQX, R-CPP and Gabazine respectively. Recordings were made from murine brain organoids with gabazene 15 minutes after addition of the compound. Subsequent recordings were performed after the addition of NBQX and R-CPP. Data was spike sorted and analyzed following the methods described above.

#### Neonatal murine brain-slice preparation

All experiments involving murine neonatal acute slice recordings were approved by the Basel-Stadt veterinary office according to Swiss federal laws on animal welfare. Briefly, mouse pups (P12-14; both sexes; C57BL/6JRj from Janvier Labs) were decapitated under isoflurane anesthesia, followed by brain dissection in ice-cold artificial CSF (aCSF) bubbled with carbogen gas (95% O_2_, 5% CO_2_). To promote self-sustained cortical activity^135^, the following aCSF recipe was used (in mM): 126 NaCl, 3.5 KCl, 1.25 NaH_2_PO_4_, 1 MgSO_4_, 2 CaCl_2_, 26 NaHCO_3_, and 10 glucose, at approximately pH 7.3 when bubbled with carbogen. Coronal brain slices (370 μm) were prepared using a vibratome (VT1200S, Leica, Wetzlar, Germany). Slices were subsequently transferred to a chamber submerged in carbogenated aCSF and stored at room temperature until use.

#### Acute recordings from neonatal murine brain slices

For recordings, a brain slice containing somatosensory cortex was transferred from the storage chamber onto the sensing area of the CMOS MEA and fixated with a customized MaxOne Tissue Holder (MaxWell Biosystems, Zurich, Switzerland). The slice was perfused with heated aCSF (32-34 °C). Sparse, rectangular electrode configurations were selected to find active regions of the somatosensory cortex, with a sparsity of two or three to allow for a good spike-sorting performance.

#### Primary planar culture preparation

The presented primary neuronal recordings (Pr) were performed and spike sorted as described in Yuan *et al*.^32^ for Pr1-4 and as described in Bartram *et al*.^33^ for Pr5-10. Briefly, neuronal cultures according to Yuan *et al*. were prepared from embryonic day 18 Wistar rat cortices and plated at a density of 3,000 cells/mm^2^ onto high-density CMOS MEAs (MaxOne, Maxwell Biosystems) and maintained in a cell culture incubator (5% CO_2_ at 37 °C). The recordings were made at 20 days *in vitro*. The recordings can be obtained here: https://www.research-collection.ethz.ch/handle/20.500.11850/431730.

#### Comparing data from different sources

The recording durations of all recordings coming from the same source were kept consistent. The recording durations per data source were selected so that each recording contained around 40 bursts (38 ± 4 bursts, mean ± SE), using the first *x* minutes of the recording to get to this value.

#### Single-unit firing rate and CV2 calculations scores

All of the following analyses were performed using custom MATLAB scripts. MATLAB version R2018b was used. The firing rate of each individual spike-sorted unit with at least 30 detected spikes in the recording was computed by obtaining the inter spike interval between each spike event and applying a Gaussian smoothing with a 50 ms kernel to its inverse. A lognormal distribution was fitted to the distribution of firing rates averaged over the whole recording period for each unit. The goodness of the fit was assessed using the *R*^2^ metric. In addition, for the same selection of units, the CV2 score of the spiking activity was computed per unit as described by Holt *et al*.^136^ as a measure of spiking variability. The same CV2 computations were performed on 100 different shuffled the spike matrices and the results from the original spike matrices were z-score normalized using the mean and standard deviation over all shuffled datasets.

#### Population rate calculations

The population firing rate was computed by summing spikes over all units per frame followed by smoothing with a 20 ms sliding square window and a subsequent 100 ms sliding Gaussian kernel. For the detection of the burst start, end and peak, population activity bursts were defined when the population-averaged spike rate exceeded 4x its RMS value (using the built in MATLAB function findpeaks with min_dist = 700 ms. For recordings with long duration bursts, min_dist was increased up to 2000 ms in order to prevent peaks in the tail of the burst from being detected as separate burst instances). The burst start and end times were determined to be the first time points where the multi-unit activity fell below 10% of the detected peak value, before and after the burst peaks respectively. The actual burst peak time was then obtained by recomputing the population firing rate using a 5 ms square window and a 5 ms Gaussian kernel and finding the frame with the highest value between the burst start and end time. For murine primary planar cultures, a 20 ms square window, a 50 ms Gaussian kernel and a 3 x RMS threshold were used for population peak detection and a 20% threshold for the burst start and end time detection. These values were chosen to account for stronger jitteriness of the population activity and more abundant inter-burst activity.

#### Firing rate sequences and burst backbones

For each individual unit, the firing rate centered by the burst peak was averaged from −250 ms to 500 ms relative to the burst peak. In addition, the time relative to the population burst peak at which this unit had a peak in its firing rate within the burst start and end window was selected. The median and variance of the firing rate peak times was computed per unit over all bursts in which this unit fired at least two action potentials. The median values were used for reordering the units for different plotting purposes and the variance was used to fit a linear mixed-effects model to study the relationship between the effect of the relative position of the peak (from 0 to 1) on the variance of the peak.

Units that fired at least two action potentials in all the bursts in a recording were defined as backbone units. For murine organoids and cortical slices, a threshold of 80% and 90% of bursts was used respectively since only a small fraction of units had at least two action potentials in all bursts (Supplementary Fig. 2A). A Backbone unit sequence was defined by ordering all backbone units based on their median firing rate peak time. For each sample, the burst backbone period was defined as the average firing rate peak time of the earliest backbone unit until the average firing rate peak time of the latest backbone unit in the sequence. For different plotting purposes, the backbone period was rescaled from 0 to 1 and data per organoid were overlaid and averaged over the rescaled backbone period for comparison.

#### Firing rate peak rank order correlations

For each burst, the firing rate peak time of all backbone units was taken and a rank was assigned to each unit. Next a Spearman rank order correlation was computed between every pair of bursts to assess the similarity in the firing rate peak sequences of the backbone units (using the built in MATLAB function corr with “Type” = “Spearman”). The same computations were performed on 100 different shuffled the spike matrices and the results from the original spike matrices were normalized using the (A-B)/(A+B) strategy, where A is the measured value and B is the averaged value computed over the 100 shuffled datasets.

#### Burst-to-burst firing rate correlations

For each unit, the firing rate was recomputed after removing all spikes that fell outside of the burst windows. Next, this firing rate was selected from −250 ms to 500 ms relative to each individual burst peak and a cross correlation was computed for the unit firing rate between each pair of bursts for each individual unit (using the built in MATLAB function *xcorr* with maxlag = 10ms and normalization = “coeff”). Only bursts with at least 2 detected action potentials and units with at least 2 spikes in at least 30% of all bursts were considered for this analysis. Afterwards the average over the maximum correlations for all the burst pairs with at least 2 detected action potentials was computed per unit, yielding the burst-to-burst correlation. The same computations were performed on 100 different shuffled spike matrices and the results from the original spike matrices were normalized using the (A-B)/(A+B) strategy, where A is the measured value and B is the averaged value computed over the 100 shuffled datasets.

In a separate analysis, burst-to-burst correlations were computed between bursts from two recordings from the same organoid slice at four-hour intervals. Average burst-to-burst correlations were computed for pairs of bursts within each of the two same recordings, as well as for pairs of bursts where one burst came from the recording at zero hours and the other burst from the recording at four hours.

#### Pairwise firing rate correlations

Using the same firing rates computed after removing spikes outside burst windows, cross-correlations were computed between each pair of units (using the built in MATLAB function *xcorr* with maxlag = 350ms and normalization = “coeff”, a maxlag of 350ms was chosen since the median backbone period over all samples except the murine primary cultures was 348ms). The rate for the whole recording was used. The maximum correlation values for each unit pair were compared between pairs of backbone units, pairs of one backbone and one non-rigid unit and pairs of non-rigid units. The same computations were performed on 100 different shuffled spike matrices and the results from the original spike matrices were normalized using the (A-B)/(A+B) strategy, where A is the measured value and B is the averaged value computed over the 100 shuffled datasets. In addition, for all pairs of backbone units, the lag time corresponding to the maximum correlation value was compared to the average absolute lag time over all shuffled datasets for the same unit pair.

#### Burst clustering

For all the detected bursts, the firing rate per unit was selected for a window of −250 ms until 500 ms relative to the burst peak. Similar to Segev *et al*.^71^, the pairwise correlations in the firing rates were computed for all unit pairs, using the firing rates for this single burst window. Subsequently, the bursts were clustered by performing a k-means++ clustering on the pairwise correlation matrices. The optimal number of clusters was selected using the elbow method. The clustering results were assessed by projecting the pairwise correlation matrices per burst onto the first two principal components, labeled by their cluster and cluster separability was confirmed. Next, to assess variability in the firing of a unit between different burst clusters, the average firing rate per unit was computed for each burst cluster and the CV score (STD/mean) was computed to quantify the firing rate differences between burst clusters per unit. This difference was compared between backbone units and non-rigid units using the statistical analysis as described in *Methods: statistical analyses for model comparisons*.

#### Burst similarity score

At every frame relative to the burst peak, a vector containing the firing rates for each unit was obtained. For every pair of bursts, the cosine similarity was computed between the vectors from the two different bursts. This yielded a matrix with pairwise burst similarity values at every frame relative to the burst peak. The average of this matrix was defined to be the burst similarity score for that relative frame. This score was computed for every frame in the period from the earliest burst start time - relative to the burst peak over all bursts - until the latest burst end time - relative to the burst peak over all bursts. The same computations were performed on the spike matrices after shuffling.

Besides computing the burst similarity score over all units, burst similarity scores were also computed for only a subset of units. In the first case, these subsets consisted of all backbone units and all non-rigid units respectively. Subsequently, at each frame the burst similarity score distribution for all burst pairs was compared using a paired sample, two-sided t-test (using the built in MATLAB function t-test). In the second case, these subsets consisted of units with an average correlation value (*Methods: Pairwise firing rate correlations*) in the top/bottom *i*^th^ percentile where *i* ranged from 20 to 95. Subsequently, the difference between the top and bottom *i*^th^ percentile was quantified for this range as the sum of the burst similarity score over all frames in the backbone period. This was done to assess the burst similarity based only on highly/lowly correlated units. Similarly, the burst similarity score distribution for all burst pairs was compared between the top/bottom 20 percent of units and all units using a paired sample, two-sided t-test.

#### PCA manifold analysis

The spike rate matrix of an organoid with *n* units can be interpreted as a set of points in *n* dimensional space, where each axis holds the spike-rate trajectory of a specific unit. The principal components (PC) of this system are the directions in this space that capture the majority of the dataset’s variance. A dimensionality reduction is achieved by linearly projecting the dataset onto these PCs. This transformation collapses the *n* dimensional system to *p* dimensions where *p* < *n*, while preserving the dominant patterns exhibited by the system. For this analysis, the PCs are computed by the Eigen-decomposition of the covariance matrix computed as follows:

Prior to the dimensionality reduction step, the firing rate data was normalized for each unit individually using the *z*-score method, which centers the data around zero mean and unit standard deviation. The dimensionality reduction was performed on three separate selections of units: all units, backbone units only and non-rigid units only.

The cumulative sum of the variance explained per PC was computed for the PCs ordered from high variance to low. For each recording, the results for all units were subtracted from the results for backbone units only and non-rigid units only. Negative values mean that the cumulative sum of the variance explained by all units is larger than for the subset of units and positive values mean that the cumulative sum of the variance explained by all units is smaller.

Furthermore, the sum of the variance explained by the first three PCs was computed and divided by the summed explained variance of the first *X* principal components, where *X* is the lowest number of total PCs from the three selections, all units, backbone and non-rigid. This was done to account for differences in the total number of PCs per selection. This value was computed for the original data and for 100 different shuffled spike matrices and the results from the original spike matrices were normalized using the (A-B)/(A+B) strategy, where A is the measured value and B is the averaged value computed over the 100 shuffled datasets. These scores were compared between the three selections and between the different model types as described in *Methods: statistical analyses for model comparisons*.

#### Randomized recording

Randomization of single-unit spike times were performed based on the methods of Okun *et al*.^58,73^ to preserve each neuron’s mean firing rate as well as the population averaged firing rate distribution. This is necessary to avoid trivial differences that would arise simply by changes in the mean firing rate of a neuron. Briefly, whenever analyses were performed on a randomized recording, the randomization was done as follows (unless stated otherwise): Two separate units, A and B, were selected and two separate frames, 1 and 2, were selected where A but not B fired in frame 1 and B but not A fires in frame 2. Next, the spikes from unit A and B were switched between frames 1 and 2. The resulting spike matrix still has an equal number of spikes per unit (same average firing rate) and an equal number of spikes per frame (same population rate). This shuffling procedure was performed 5x as many times as there were spikes in the spike matrix, resulting in each spike getting shuffled 10x on average. This method was applied to produce 100 different shuffled spike matrices per original recording.

#### Statistical analyses for model comparisons

Statistical modeling was carried out in the R environment. Nested data were analyzed with linear mixed-effects models (*lmer* function of the lme4 R package^137^) with “organoid” or “unit ID” as random effect. Non-nested data were analyzed with linear models (*lm* function). Right-skewed and heavy-tailed data were log-transformed and analyzed with a linear model. Statistical significance for linear mixed-effects models was computed with the lmerTest R package^138^ and the summary (type III sums of squares) R function. Statistical significance for linear models was computed with the summary R function. When possible, model selection was performed according to experimental design. When this was not possible, models were compared using the *compare_performance* function of the performance R package^139^, and model choice was based on a holistic comparison of AIC, BIC, RMSE and R2. Model output was plotted with the *plot_model* (type=’pred’) function of the sjPlot R package^140^. 95% confidence intervals were computed using the *confint* R function. Post hoc analysis was carried out using the *emmeans* and *emtrends* functions of the *emmeans* R package.

#### Hidden Markov model analysis

Hidden Markov models (HMMs) have been widely used in computational biology, ranging from protein modeling^141^ to determining evolutionarily conserved genomic elements across species^142^. More recently, this approach has been utilized to characterize the firing patterns of neuronal ensembles of specific brain states during motor function^143^, deciphering neural codes of sleep^144^ and uncovering temporal structure in hippocampal outputs^145^.

A hidden Markov model is a statistical characterization of a discrete-time random process in terms of a discrete “hidden” state, which cannot be directly observed, but which changes the probability distribution of the observations. At each time step, the value of the hidden state depends only on the previous hidden state. The observation distribution and transition probabilities together make up the parametrization of the HMM. These parameters are fitted using the Expectation-Maximization (EM) algorithm to assign the parameters of the observation distribution and transition probabilities in order to co-optimize the posterior likelihood of the observations and the sequence of hidden states.

For neuronal spiking data, HMMs are typically applied to time-binned spike matrices, where at each time step, the observation is a population activity vector consisting of the number of spikes produced by each unit within that time bin. If the bin size is significantly larger than the refractory period of the neuron, the resulting observation distribution is per-neuron independent Poisson with a parameter λ, such that the probability that the unit fires *n* times in a given time bin is given by *p*(*n*) = λ*^k^*e^-λ^/*k*! Our analysis is carried out in the Python programming language, version 3.11, using the implementation of an HMM with Poisson observations provided by the package SSM.

We validated that HMMs capture information from the spiking data via 5-fold cross-validation by comparing the posterior log likelihood of a held-out validation set to that of the randomized surrogate data for the held-out region^145^. Each “fold” consists of fitting the model parameters to a random 80% subset of the data and evaluating the fitted model on the remaining 20%. Log likelihood is always greater for the real than the surrogate data (Supplementary Fig. 24), indicating that the HMM is modeling transitions using information present in the real data but not the surrogate.

#### Hidden Markov Model Hyperparameters

Although the parameters of the observation and state transition distributions are selected by EM, there are two hyperparameters as well: the bin size *T* for converting spiking data into a discrete-time observation vector, and the number of states *K* for the HMM itself. Both must be chosen independently of the EM fitting process, so we treat them as hyperparameters and evaluate performance for a range of values using the 5-fold cross-validation method described above. We performed this validation across all 8 human brain organoid recordings. We first chose a default bin size of 30 ms based on the characteristic time scale of bursts. Under this condition, fit performance is insensitive to the number of hidden states above 10 (Supplementary Fig. 23), so all further analysis is conducted across models with the number of hidden states ranging from 10 to 30. We next evaluated performance across bin sizes of 10, 20, 30, 50, 70, and 100 ms, for numbers of hidden states ranging from 10 to 50 (Supplementary Fig 22). Performance is also relatively insensitive to bin size near 30 ms, but significantly larger or smaller bin sizes do exhibit somewhat worse performance. The analysis reported on in the main paper uses a fixed bin size of 30 ms, but the number of hidden states ranges from 10 to 30.

#### Hidden State Trajectories

The ability of a HMM to capture the stereotyped dynamics of bursts in human brain organoids is explored in Figure 5A-C. Given a fitted HMM, we estimate the most probable sequence of hidden states for a given time-binned spike matrix using the forward-backward algorithm, a standard technique for maximum likelihood estimation of hidden state. Then, this sequence of hidden states is registered relative to the peaks of all the bursts in the recording, and trimmed into fixed-length subsequences corresponding to a fixed time window surrounding each peak. For visualization purposes, the parameters were set to 300 ms before and 600 ms after. We then computed the empirical probability distribution over hidden states as a function of time relative to the burst peak by counting how many times each state appears at each position across all subsequences, divided by the total number of subsequences. Results are shown for one representative HMM (with 20 hidden states) in the first organoid we analyzed (with 30ms time bins).

We also measured the rate of hidden state traversal during the burst backbone time period (*Methods: firing rate sequences and burst backbones*). We split the maximum-likelihood state sequence into subsequences corresponding to one backbone period around each burst peak, and calculated the total number of hidden states divided by the duration of the burst backbone period. The distribution of these state traversal rates was significantly different for all three biological models (Supplementary Fig. 34), but much more similar between murine slices and human brain organoids than between either model and murine primary cultures.

#### Consistency of units within a Hidden State

We refer to each time bin of the spike matrix in which a given hidden state is most probable as a “realization” of that state. We then computed the consistency of each unit in each hidden state as the fraction of realizations of that hidden state in which the unit fires at least once. This procedure yielded an array of consistency scores with one row for each of the model’s *K* hidden states, and one column for each unit (Fig. 5D, Supplementary Fig. 27).

We view the columns of this array as vectors in a *K*-dimensional space; Figure 5E visualizes a concrete example of this space, using PCA dimensionality reduction to demonstrate that backbone and non-rigid units are almost linearly separable. We measured the degree of this approximate linear separability by determining an optimal decision boundary within the original space using a linear support vector machine (SVM).

Linear separability was quantified as the accuracy of the linear classifier according to leave-one-out cross-validation: for each unit, the model was fitted to all other units, and its accuracy was evaluated on the single held out unit. The single-unit accuracy (always either 0 or 1) was then averaged across all units. Then, as a null hypothesis to control for the fact that backbone units fire more overall than non-rigid units, we calculated the linear separability of backbone and non-rigid units based solely on their overall firing rate. This does not depend on the fitted HMM, so the null hypothesis has only one score per organoid (the linear separability across all units), rather than the distribution across models produced by the SVM-based linear separability metric.

#### Dimensionality of a Fitted HMM

To evaluate the ability of HMMs to represent non-trivial activity manifolds beyond simple variations in population firing rate, we estimated the dimensionality of the observation model in each fitted HMM. The population observation matrix is an array of shape [number of hidden states] × [number of units], where each entry represents the parameter λ of the Poisson distribution estimated for the firing of that unit in that state. We performed a singular value decomposition (SVD) on this matrix so that a principal component analysis with any desired number of dimensions *d* could be acquired by projecting only the first *d* components of the SVD. Furthermore, this same transformation can be applied to the time-binned spike matrix itself in order to yield a dimension-reduced version for visualization purposes (Supplementary Fig. 28).

We defined the “dimensionality” of the trained HMM for a given dataset to be the number of principal components required to meet a given threshold θ in the percent explained variance on the HMM states themselves. The statistical significance of this finding was calculated using a generalized linear mixed-effects model (*glmer* from the lme4 R package) with Poisson-family observations and a logarithmic linkage function. This differs from other results using a standard linear mixed-effects model due to the dimension being a discrete quantity, which cannot reasonably be approximated as Gaussian. For the same reason, normalization against the randomized baseline was not performed in this analysis; the original integer dimensionality values were modeled.

#### Non-Poisson Units

Under the null hypothesis, which was originally assumed in fitting HMMs to our data in the first place, each unit produces Poisson firing at a rate which depends on the hidden state. For a Poisson unit, for each hidden state of each fitted HMM, the number of firings of the unit in each realization of the hidden state should follow a Poisson distribution. This distribution has mean and variance both equal to its single parameter λ, so we calculated the mean number of firings of a unit across all realizations of a hidden state, and performed a one-sided chi-square test to identify whether the variance in firings for that unit was less than under the Poisson null hypothesis at the *P* = 1% significance level. This value was then averaged across all hidden states (weighted by the number of realizations of each state) to yield an overall measurement of the conditional deviation from Poisson statistics given the hidden state information (Supplementary Fig. 33D).

#### *d_2_* calculation

To assess temporal scale-invariance in network dynamics, we calculated the distance metric *d_2_* following methods developed by Sooter *et al*.^62^. For each recording, we first created a population activity time series by counting all spikes across all neurons in 30 ms time bins. This time series was z-scored (mean subtracted and normalized by standard deviation). We then fit an order-20 autoregressive (AR) model to the z-scored time series using the Yule-Walker method. The AR model describes how current activity depends on past activity through a history kernel that spans 20 time steps, capturing temporal correlations across multiple timescales. For time-shuffled controls, we randomly shuffled the spike times of each neuron independently while maintaining their firing rates, then performed the same binning, z-scoring, and AR model fitting procedures.

The AR model coefficients were used to calculate *d_2_*, which quantifies the Euclidean distance from the model parameters to the β=2 critical fixed point in the temporal renormalization group framework. This fixed point represents a type of criticality characterized by power-law temporal correlations with exponent β=2. Values of *d_2_* near zero indicate dynamics close to criticality, characterized by scale-invariant temporal fluctuations that persist across a broad range of timescales. In contrast, larger values of *d_2_* indicate deviation from criticality, reflecting dynamics dominated by a characteristic timescale or lacking long-range temporal correlations. Time-shuffled controls, which destroy temporal correlations while preserving basic firing statistics, typically yield *d_2_* values greater than 0.2. The geometric interpretation of *d_2_* is the minimum distance between the AR model coefficients and a hyperplane in coefficient space that contains all models that flow to the β=2 fixed point under the temporal renormalization group transformation.

Not all neural activity patterns are well-described by autoregressive models. For example, highly bursty dynamics with long silent periods punctuated by brief population-wide activation can violate the AR model’s assumptions about temporal continuity. Therefore, before calculating *d_2_*, we assessed the quality of the AR model fit for each preparation. We compared the avalanche size distributions, avalanche duration distributions, and power spectra of the original data with those generated by simulating the best-fit AR model. Preparations were excluded from *d_2_* analysis if the AR model failed to capture these key statistical features of the empirical data, specifically if the model-generated distributions deviated from the empirical distributions by more than one order of magnitude. This approach excluded 5/8 human organoid preparations, 2/9 mouse organoid preparations, 2/6 acute slice preparations, and 0/7 2D culture preparations from *d_2_* analysis, ensuring that our assessment of criticality was based only on preparations whose temporal structure was accurately captured by the AR framework.

## Data Availability

The data supporting the findings of this study are available within the article and its supplementary information. Raw and curated electrophysiology recordings can be found here https://dandiarchive.org/dandiset/000732

## Code Availability

Spike sorting was performed in Python 3.6 using SpikeInterface 0.13.0 and previously published^110^, which can be found at https://github.com/SpikeInterface/spikeinterface. Custom code for electrophysiology analysis is available at https://github.com/braingeneers/Protosequences

## Acknowledgments

The authors would like to thank members of the *Braingeneers* consortium for helpful discussions and David Haussler for insightful comments. We would also like to thank members of the UC Santa Cruz Genomics Institute for help with computing resources, in particular David Parks for assistance with archiving the neurophysiology data. This study was funded by the National Science Foundation (NSF) awards CNS-1730158, ACI-1540112, ACI-1541349, OAC-1826967, OAC-2112167, CNS-2100237, CNS-2120019, the University of California Office of the President, and the University of California San Diego’s California Institute for Telecommunications and Information Technology/Qualcomm *Institute,* Schmidt Futures Foundation SF857 (M.T.), German Research Foundation FOR5159 TP1 (437610067) (I.L.H.-O.), European Research Council (ERC) Advanced Grant 694829 ‘neuroXscales’(A.H.), Swiss National Science Foundation project 205320_188910/1 (A.H.), NIH T32 ES007141 and International Foundation for Ethical REsearch (D.M.A.E.D.), Hopkins Discovery and Johns Hopkins SURPASS (L.S.), John Douglas French Alzheimer’s Foundation (K.S.K.), NIH BRAIN Initiative R01NS118442 (K.B.H.).

## Author contributions

T.S. designed, conceived and supervised the study; M.C., I.L. H.-O., K.B.H., and K.S.K. offered numerous suggestions and comments; T.V.D.M., A.S. and M.C. performed computational analysis and statistics on electrophysiology recordings; J.B. performed extracellular recordings on acute brain slices under the supervision of A.H.; S H., G.A.K., H.E.S., C.D. and S.M. cultured murine brain organoids and performed electrophysiology measurements under supervision of T.S. and M.A.M-R.; S. H., G.A.K., H.E.S. performed single-cell RNA sequencing and immunohistochemistry of murine organoids under the supervision of M.A.M-R., B.M.C. and T.S.; C.R.K.H. performed additional immunohistochemistry and fluorescence microscopy under supervision of T.S.; S. H., H.E.S. and J.G-F. performed analysis on single cell RNA sequencing data from murine organoids under the supervision of M.A.M-R. and B.M.C.; D-M.A.D., J.L, M.S. performed electrophysiology measurements and bulk RNA sequencing of additional human brain organoids under the supervision of L.S.; A.D. and Z.Z. performed additional electrophysiology analysis under the supervision of T.V.D.M., L.R.P. and P.K.H.; K.B-N. performed computational analysis under the supervision of K.B.H.; L.J.B. archived neurophysiology data sets; T.S. wrote the first draft of the manuscript, M.C., I.L.H.-O., K.B.H., T.V.D.M. and K.S.K. provided valuable edits to subsequent drafts, and all authors discussed the results and commented on the manuscript.

