## Supplementary Information for "Protosequences in brain organoids model intrinsic brain states"

**This PDF file includes:**

Supplementary Figures 1 to 35

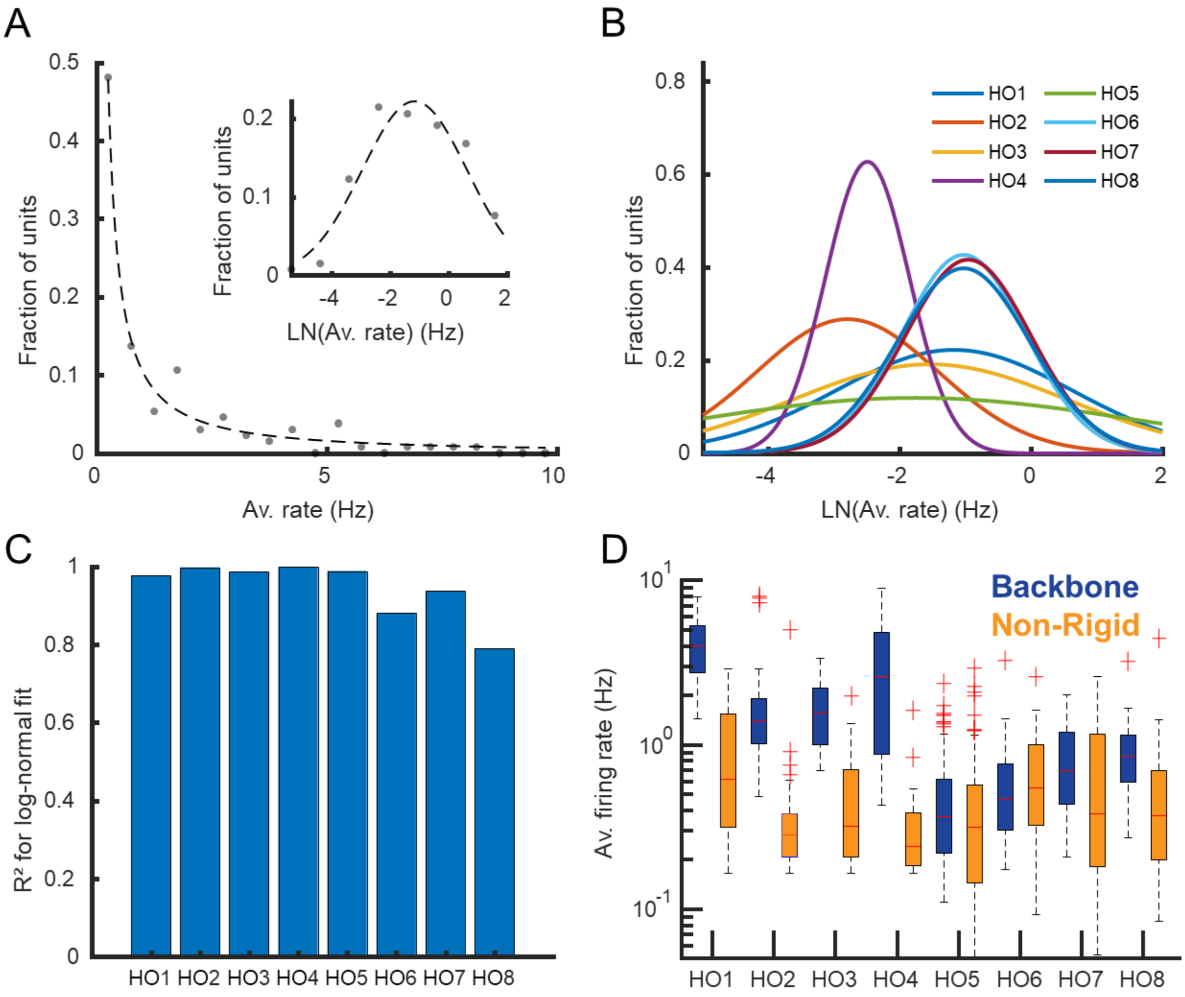


**Supplementary Figure 1. Average firing rate distributions are skewed with backbone units in the tail.** (**A**) A histogram of the distribution of average firing rates for all units in organoid 1. The majority of units have low average firing rates while a long tail in the distribution contains a small subset of units with high average firing rates. A lognormal distribution is fitted to the histogram. The inset shows the histogram for the logarithm of the average firing rates of the same organoid. A normal distribution is fitted to the histogram. (**B**) Normal distributions fit to the logarithm of the average firing rate per unit for the 8 different organoids. (**C**) The R-squared values for the fitted normal distributions shown in **B**. $R^2 = 0.97 \pm 0.04$ (mean±STD) across the 8 human brain organoids. Backbone neurons alone are not well described by a lognormal distribution. $R^2$ values are 0.45±0.30 across the 8 human brain organoids. (**D**) The distribution of average firing rates per organoid for backbone and non-rigid units separated. The backbone units populate the tail of the skewed average firing rate distributions in all organoids. See **Fig. 7A** for statistical comparisons.

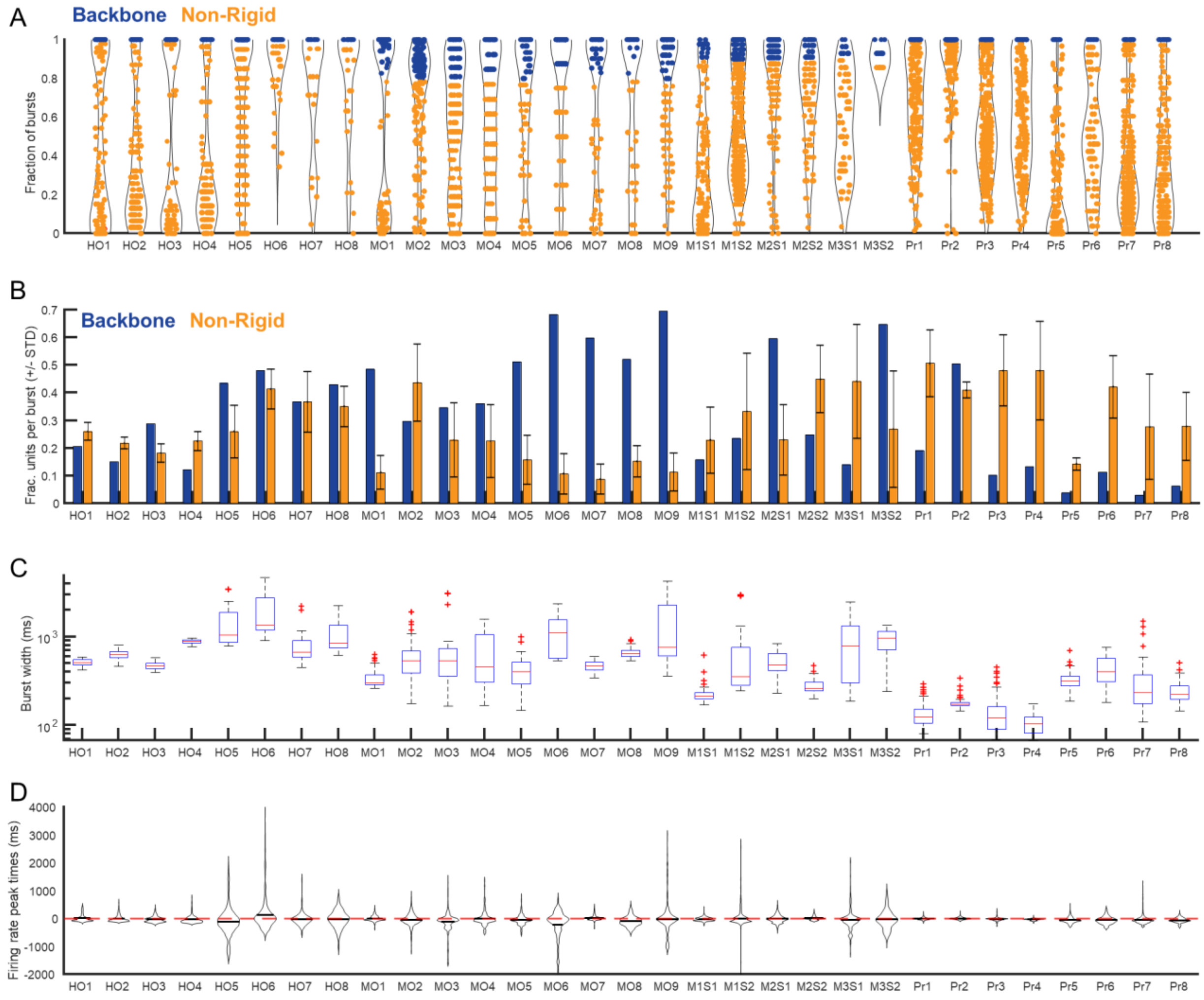


**Supplementary Figure 2: Backbone units during bursts.** (**A**) Fraction of bursts in which a unit has at least two spikes. Every dot represents a unit. Blue dots are the backbone units and have at least two spikes in all bursts. Red dots are the non-rigid units that spike at least two spikes in less than all bursts. The backbone unit threshold for murine organoids and murine neonatal slices was lowered to at least 2 spikes in at least 80% and 90% of all bursts respectively. (**B**) Fraction of backbone units and non-rigid units that are active per burst. Backbone units are active in all bursts and so the fraction of backbone units is the same for each burst. The fraction of non-rigid units that fire at least two spikes in all bursts varies, depicted by the error bars that represent the standard deviation over all bursts. (**C**) Burst durations are plotted for all different recordings. (**D**) The distribution of single-unit firing rate peak times relative to population burst peaks for all different recordings.

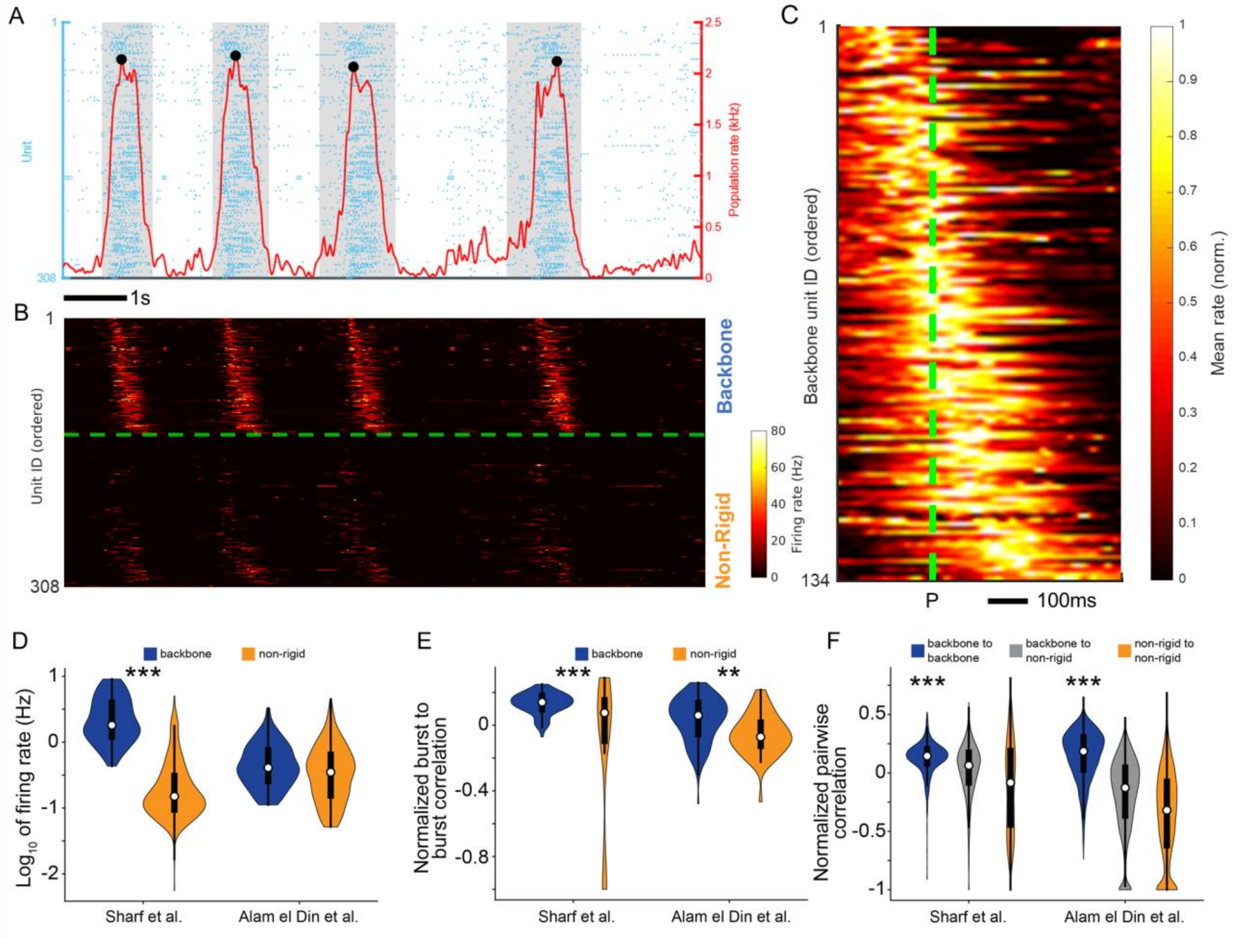


**Supplementary Figure 3: Consistent firing patterns in human brain organoids.**
(**A**) Raster plot visualization of single-unit spiking (blue dots) measured across the surface of a human brain organoid slice from Alam El Din *et al*. (HO5), positioned on top of the same Maxwell Biosystems microelectrode array as used for the organoid recordings included in the main figures. The population firing rate is shown by the red solid line. Population bursts are marked by sharp increases in the population rate. Burst peak events are denoted by local maxima (black dots) that exceed 4x-RMS fluctuations in the population rate. The shaded gray regions denote the burst duration window as defined by the time interval in which the population rate remains above 10% of its peak value in the burst. (**B**) The instantaneous firing rate of single-unit activity from panel A after reordering. The backbone units are plotted above the dashed line while non-rigid units are plotted below the dashed line. In each category, units are ordered based on their median firing rate peak time relative to the burst peak, considered over all bursts in the recording. (**C**) The average burst peak centered firing rate measured across all burst events for the example recording of which part is shown in A. The burst peak is indicated by the dotted line. The unit order is the same as **B**. Note the progressive increase in the firing rate peak time relative to the burst peak, as well as a spread in the active duration for units having their peak activity later in the burst. The average firing rate is normalized per unit to aid in visual clarity. (**D**) The distributions of the log of the average firing rate per unit, separated for backbone and non-rigid units. All units from the 4 Sharf recordings are pooled together and all units from the 4 Alam El Din recordings are pooled together. (**E**) The distributions of the average burst to burst

correlations per unit after average rate normalization, separated for backbone and non-rigid units. All units from the 4 Sharf recordings are pooled together and all units from the 4 Alam El Din recordings are pooled together. Note the significant difference between backbone and non-rigid units present for both sets of recordings (*** = $P < 0.001$, ** = $P < 0.01$, * = $P < 0.05$, linear mixed-effect model). (**F**) The distributions of the pairwise correlations per unit pair after average rate normalization, separated for backbone pairs, backbone and non-rigid combinations and non-rigid pairs. All unit pairs from the 4 Sharf recordings are pooled together and all unit pairs from the 4 Alam El Din recordings are pooled together. Note that the normalized correlation for backbone pairs are significantly larger than 0 for both sets of recordings (*** = $P < 0.001$, ** = $P < 0.01$, * = $P < 0.05$, linear mixed-effect model).

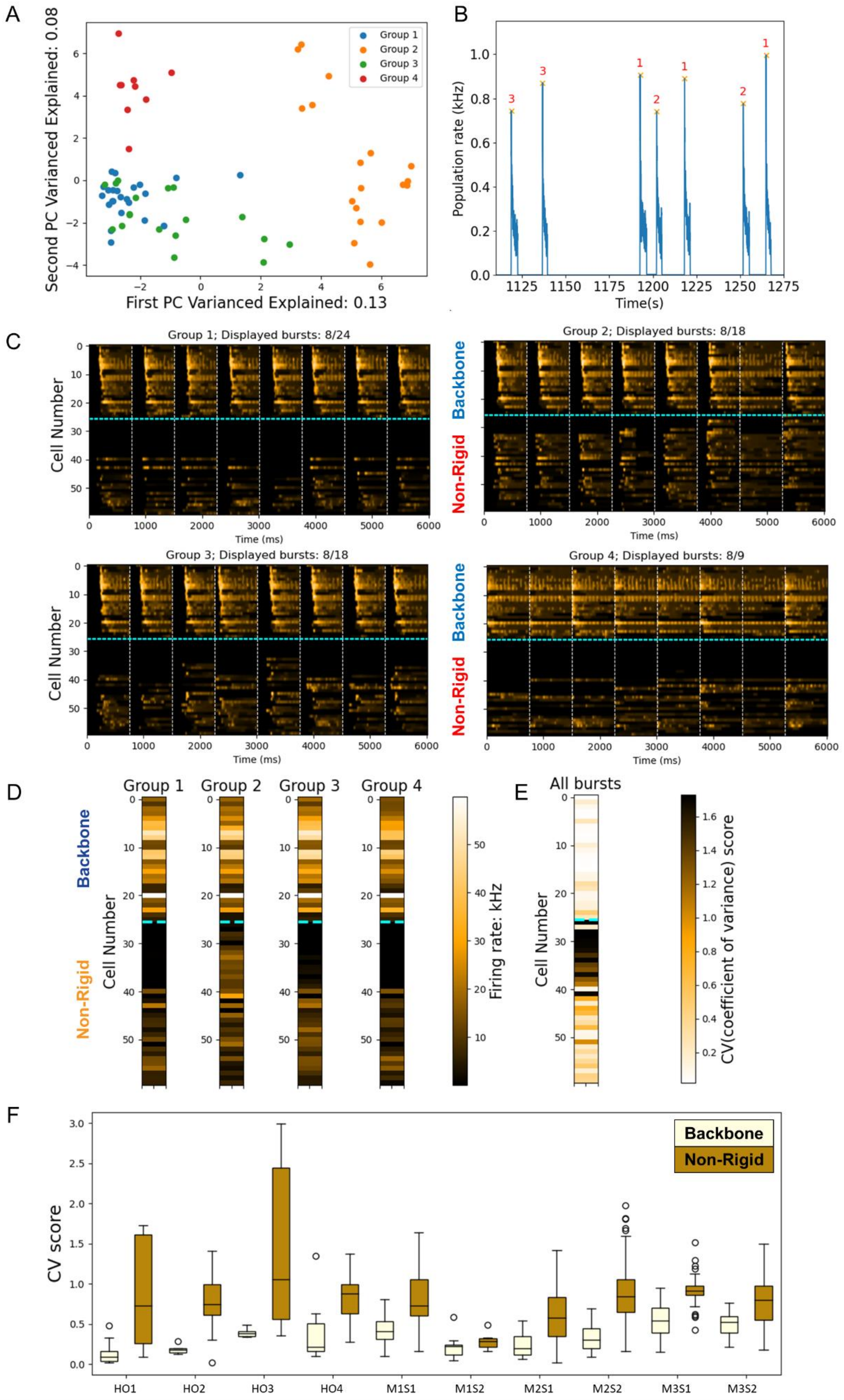

A
B
C
D
E
F
First PC Varianced Explained: 0.13
Second PC Varianced Explained: 0.08
Group 1
Group 2
Group 3
Group 4
Population rate (kHz)
Time(s)
Group 1; Displayed bursts: 8/24
Group 2; Displayed bursts: 8/18
Group 3; Displayed bursts: 8/18
Group 4; Displayed bursts: 8/9
Cell Number
Time (ms)
Backbone
Non-Rigid
Firing rate: kHz
All bursts
CV(coefficient of variance) score
CV score
HO1
HO2
HO3
HO4
M1S1
M1S2
M2S1
M2S2
M3S1
M3S2

**Supplementary Figure 4. Burst clustering reveals between-cluster variability for non-rigid units but not for backbone units.**
(**A**) Pairwise firing rate correlations per burst (computed over a window ranging from -250 ms until 500 ms relative to the burst peak) projected onto the first two principal components, labeled by the identified clusters show a clear separation between different burst clusters. The results for example recording Or5 are shown. (**B**) The population rate for a snippet of the recording for Or5 covering several bursts labeled by their cluster. (**C**) Firing rates per unit for 8 different example bursts per cluster. (**D**) The average firing rate per unit for the different detected burst clusters. (**E**) A selection of non-rigid units is most variable in their activity between the different burst clusters as reflected by a higher CV score for their firing rate in the different burst clusters (CV score are computed per row for the 4 columns shown in **D**. (**F**) The CV scores for the firing rate over the different burst clusters is significantly higher for non-rigid units compared to backbone units. ($P \leq 10^{-20}$ for difference between backbone and non-rigid, linear mixed-effect model)

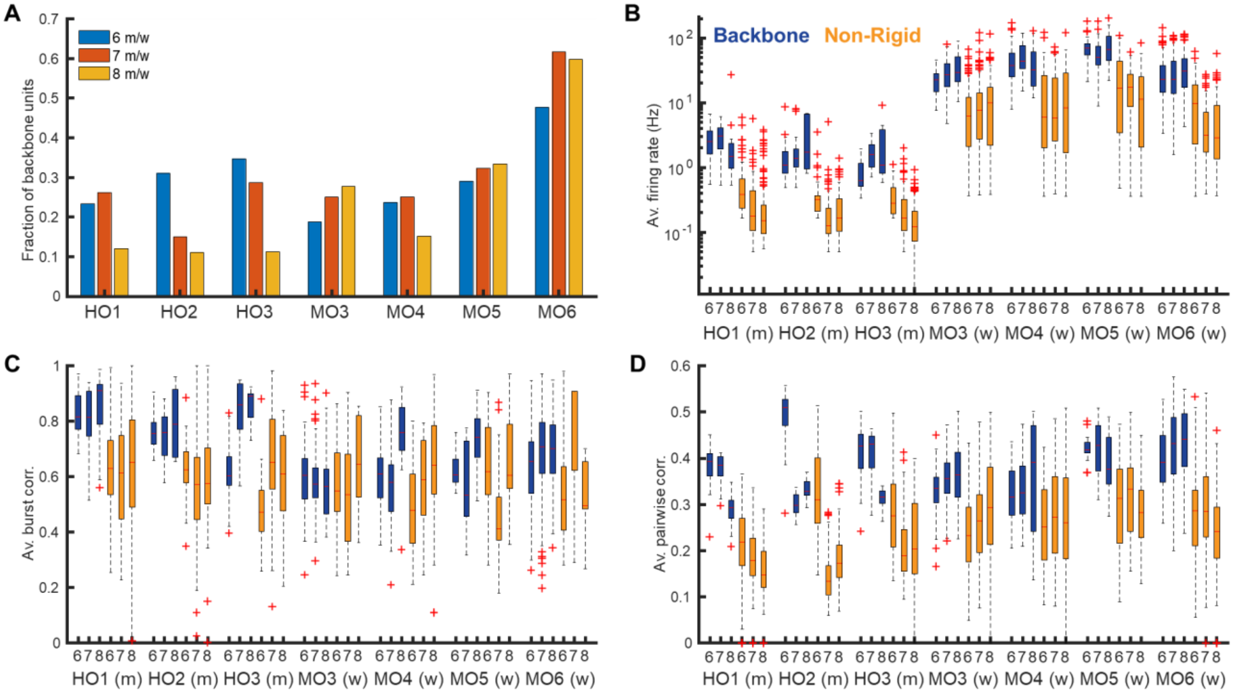


**Supplementary Figure 5. Backbone units during development.** (**A**) The fraction of backbone units decreases when the organoid matures. Human organoid (HO) ages are reflected in months whereas murine organoid (MO) ages are reflected in weeks. (**B**) At all recorded ages, the average firing rate per unit is on average higher for the backbone units than for non-rigid units. (**C**) At all recorded ages, the average burst to burst correlation per unit is on average higher for the backbone units than for non-rigid units. Furthermore, the average burst to burst correlation increases over the course of development (2-way ANOVA, impact of age on burst to burst correlation HO: P = 4.14e-6, MO: P = 9.45e-9). (**D**) At all recorded ages, the average pairwise correlation between pairs of backbone units is on average higher than for pairs of non-rigid units.

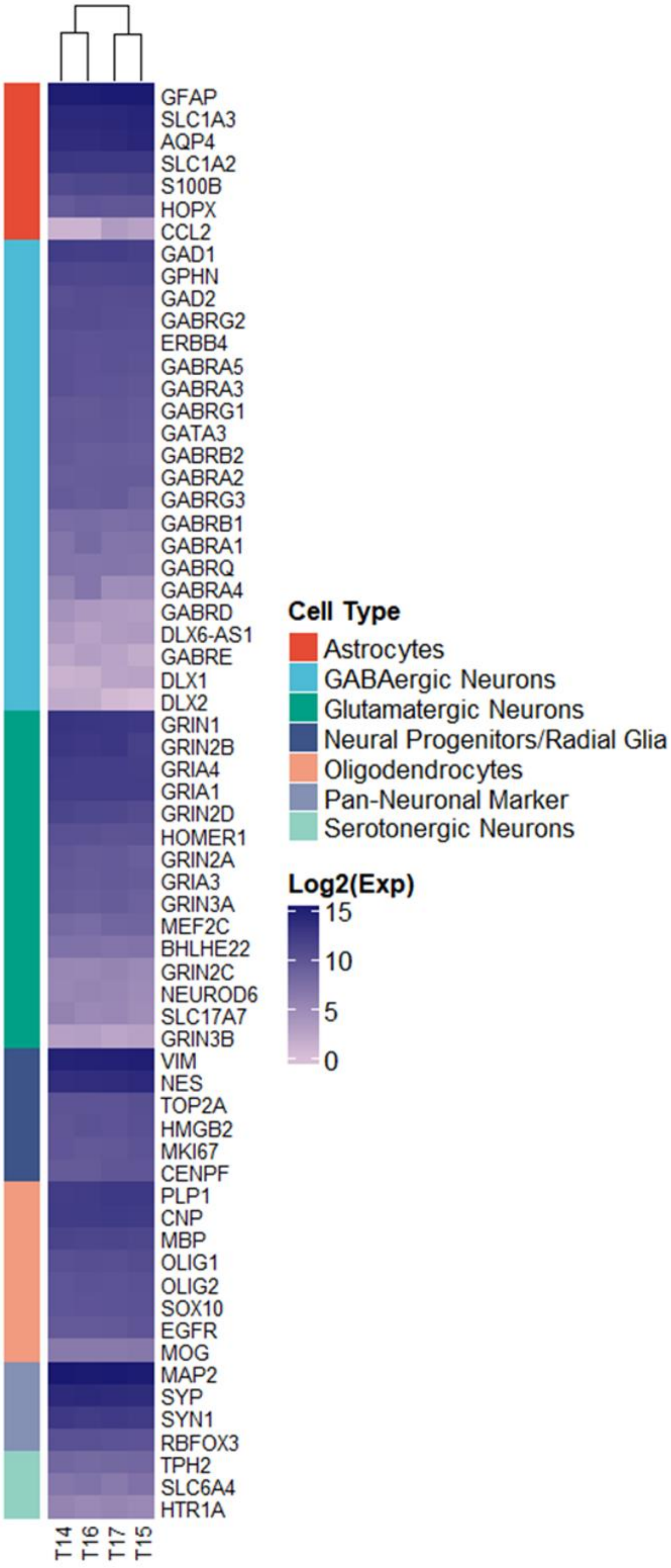
GFAP
SLC1A3
AQP4
SLC1A2
S100B
HOPX
CCL2
GAD1
GPHN
GAD2
GABRG2
ERBB4
GABRA5
GABRA3
GABRG1
GATA3
GABRB2
GABRA2
GABRG3
GABRB1
GABRA1
GABRQ
GABRA4
GABRD
DLX6-AS1
GABRE
DLX1
DLX2
GRIN1
GRIN2B
GRIA4
GRIA1
GRIN2D
HOMER1
GRIN2A
GRIA3
GRIN3A
MEF2C
BHLHE22
GRIN2C
NEUROD6
SLC17A7
GRIN3B
VIM
NES
TOP2A
HMGB2
MKI67
CENPF
PLP1
CNP
MBP
OLIG1
OLIG2
SOX10
EGFR
MOG
MAP2
SYP
SYN1
RBFOX3
TPH2
SLC6A4
HTR1A
T14
T16
T17
T15
Cell Type
Astrocytes
GABAergic Neurons
Glutamatergic Neurons
Neural Progenitors/Radial Glia
Oligodendrocytes
Pan-Neuronal Marker
Serotonergic Neurons
Log2(Exp)
15
10
5
0

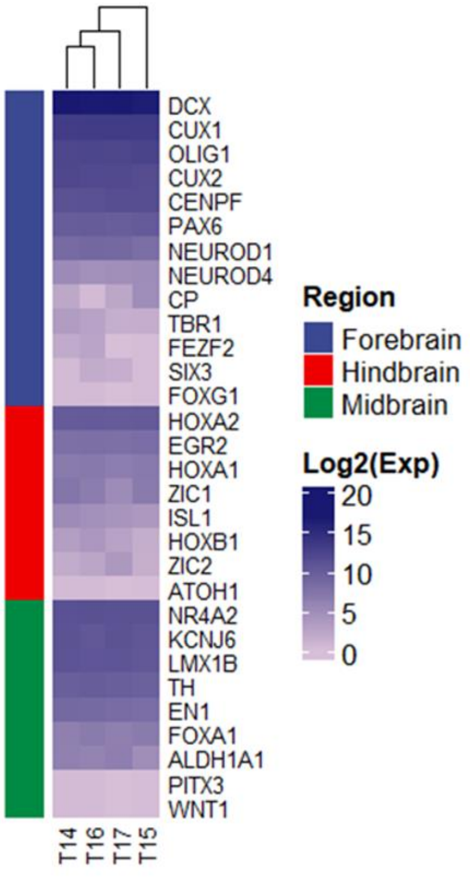
DCX
CUX1
OLIG1
CUX2
CENPF
PAX6
NEUROD1
NEUROD4
CP
TBR1
FEZF2
SIX3
FOXG1
HOXA2
EGR2
HOXA1
ZIC1
ISL1
HOXB1
ZIC2
ATOH1
NR4A2
KCNJ6
LMX1B
TH
EN1
FOXA1
ALDH1A1
PITX3
WNT1
T14
T16
T17
T15
Region
Forebrain
Hindbrain
Midbrain
Log2(Exp)
20
15
10
5
0

**Supplementary Figure 6.** Marker gene expression representing (A) main cell lineages and (B) regions of developing brain in 8-week-old brain organoids. Heatmap represents the Log2 of relative gene expression extracted from limma-voom normalized bulk RNA-seq (data sourced from El Din *et al*.).

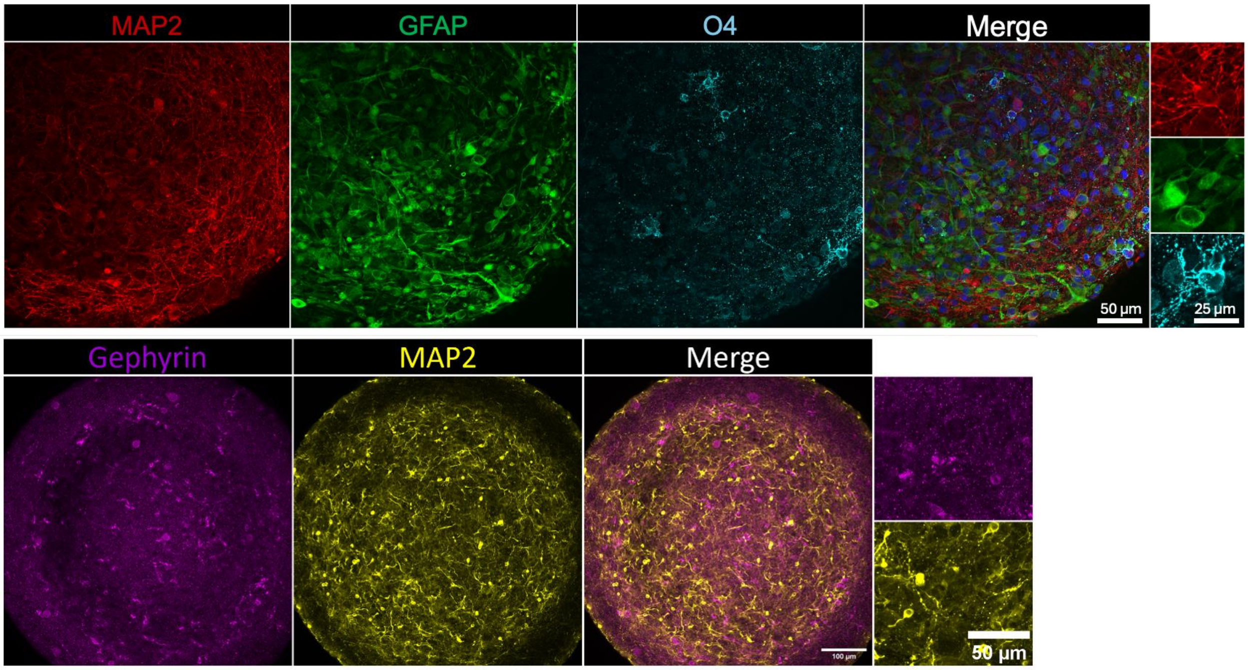


**Supplementary Figure 7.** Immunohistochemistry validation of human brain organoid cell types. Top row, somatodendritic marker MAP2 (red), astrocyte marker GFAP (green), oligodendrocyte marker O4 (turquoise). Nuclei are stained with Hoechst33342 (blue). Bottom row, inhibitory synaptic marker Gephryn (purple), dendritic marker MAP2 (yellow).

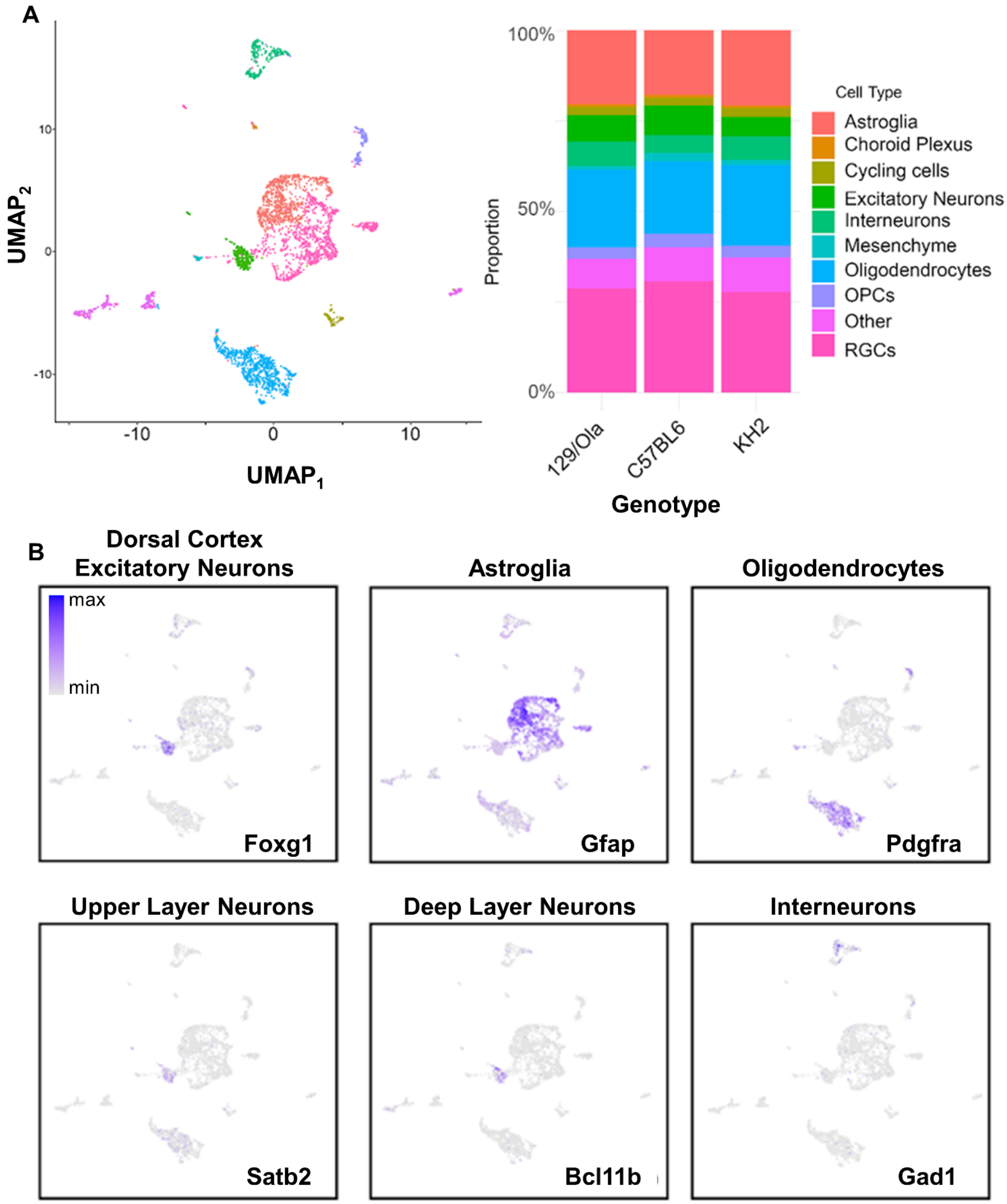


**Supplementary Figure 8. Mouse ESC cortical organoid identification of cell types using single-cell sequencing.** (A) Left, single-cell sequencing UMAP plot colored by cell type cluster. Right, relative proportions (fractions) of cells per cluster. Clusters are identified from 30-day-old organoids as determined by PIPseq. A total of 4031 cells were sequenced from three separate cell lines (C57/BL6, E14, and KH2 ESC, see Methods). (B) Selection of canonical markers used to identify cell types in single-cell data. UMAP plots are colored by normalized and scaled expression values. Expression patterns of brain-region specific marker genes reveal predominant forebrain identity of the murine brain organoids used in this study. (C) Murine organoids express multiple genes encoding subunits of AMPA- and NMDA-types of glutamate receptors as well as GABA receptor subunits and various glial cell type markers. UMAP plots are colored by expression level of known genes encoding different receptor subunits. Not detected in single-cell data were GABRA6 and GABRP.

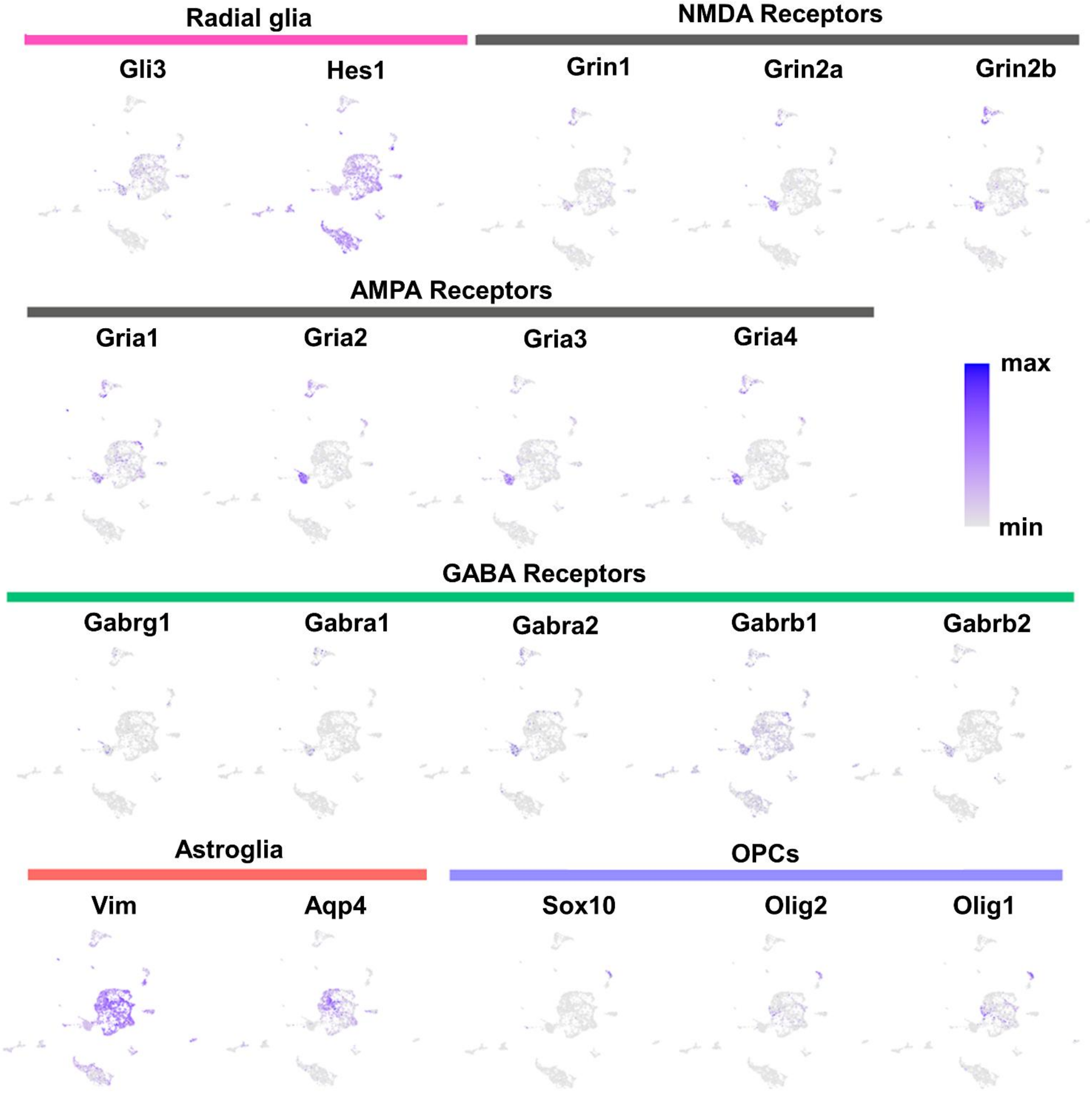


**Supplementary Figure 9. Cellular identities and neurotransmitter profiles of Mouse ESC cortical organoid determined by single-cell transcriptomics**. Murine organoids express multiple genes encoding subunits of AMPA- and NMDA-types of glutamate receptors as well as GABA receptor subunits and various glial cell type markers. UMAP plots are colored by expression level of known genes encoding different receptor subunits. Not detected in single-cell data were GABRA6 and GABRP.

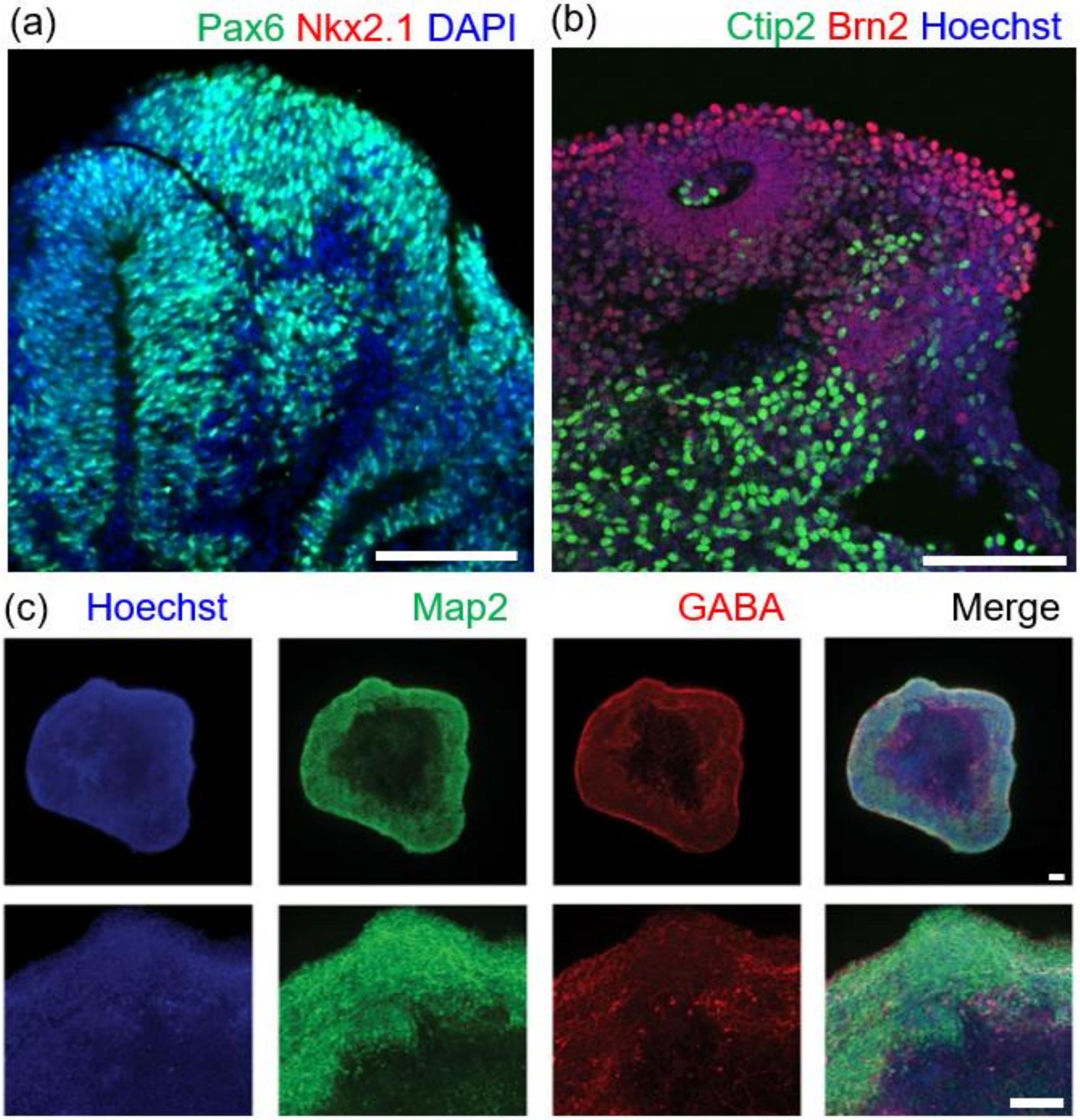


**Supplementary Figure 10. Immunohistochemistry validation of mouse ESC dorsal forebrain organoid cellular identities**. (a) Pax6-positive dorsal forebrain neural progenitors (green) shown in an 10-day organoid mouse cortical organoid counterstained with DAPI (cell nuclei, blue). Ventral forebrain marker Nkx2.1 is shown in red. (b) Ctip2-positive deep layer cortical neurons (green) in a 30-day cortical mouse organoid. Brn2-positive upper layer cortical neurons (red) are shown with Hoechst (cell nuclei, blue). (c) A subpopulation of GABA producing interneurons (red) are shown among MAP2-positive neurons (green) from a 30-day cortical mouse organoid. Scale bar 100 µm.

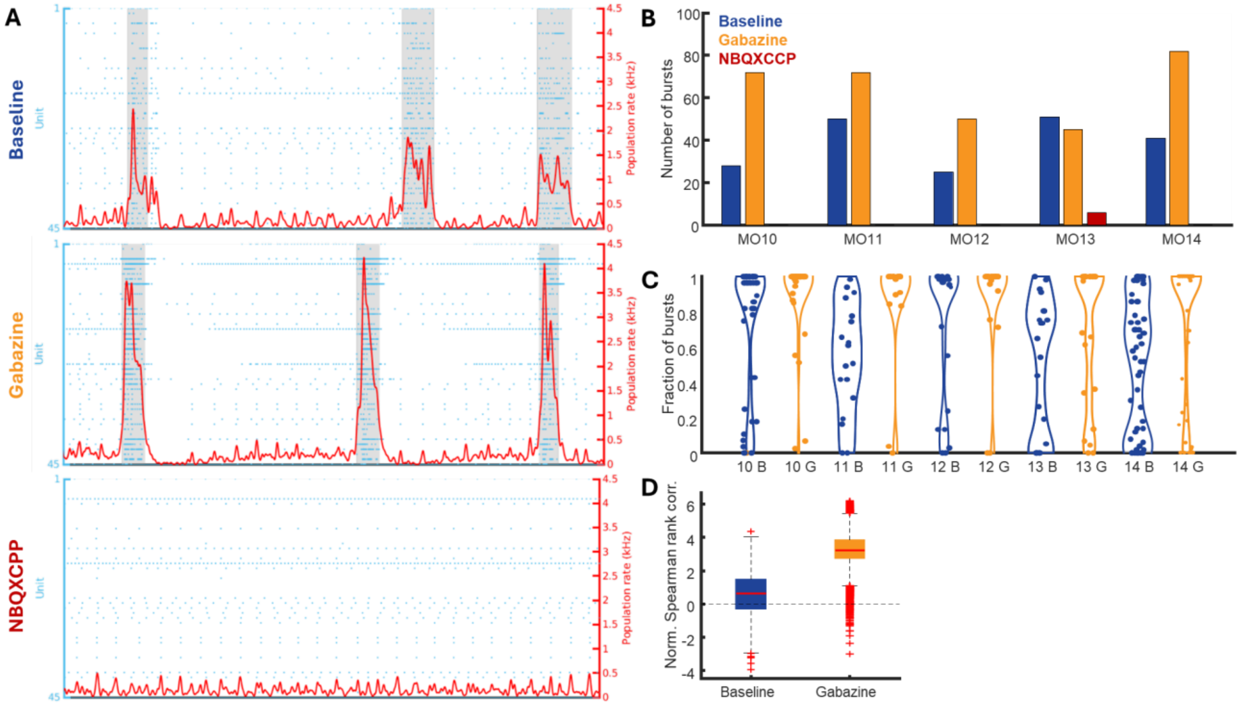


**Supplementary Figure 11: Pharmacology experiments reveal impact of excitatory and inhibitory signaling on population bursts and backbone sequences.** (**A**) Raster plot visualization of single-unit spiking (blue dots) measured across the surface of a murine organoid (MO10), positioned on top of the same type of microelectrode array as used for the recordings included in the main figures. The population firing rate is shown by the red solid line. Population bursts are marked by sharp increases in the population rate. The shaded gray regions denote the burst duration window as defined by the time interval in which the population rate remains above 10% of its peak value in the burst. Top: baseline recording. Middle: recording of the same organoid using the same electrode configuration, after treatment with 10 µM gabazine to inhibit inhibitory signaling by blocking $GABA_A$ receptors. Bottom: recording of the same organoid slice using the same electrode configuration, after blocking AMPA and NMDA receptors with bath application of NBQX (10 µM) and R-CPP (20 µM) to inhibit components of excitatory synaptic transmission. **(B)** Number of detected population bursts for 5 different murine organoids (MO10-14) under baseline conditions, after treatment with gabazine and after treatment with NBQX and R-CPP. Note that bursting disappears after NBQX and R-CPP treatment reflected as a significant decrease in bursting compared to baseline conditions ($P < 0.001$, linear mixed-effect model). Meanwhile, the number of bursts increase after gabazine treatment ($P < 0.05$, linear mixed-effect model). **(C)** Fraction of bursts in which a unit fires at least 2 spikes for the 5 different murine organoids under baseline conditions compared to the gabazine treatment. The fraction of bursts in which units are active increases significantly after gabazine treatment ($P < 10^{-7}$, linear mixed-effect model). **(D)** Normalized Spearman rank order correlations comparing the sequential order of backbone sequences for all burst pairs of 5 different murine organoids (MO10-14) under baseline conditions and after treatment with gabazine. Correlation scores are z-scored relative to shuffled spike matrices such that values above 0 indicate a more consistent backbone sequence than the shuffled data. There is a significant increase in backbone sequence order similarity after treatment with gabazine ($P < 10^{-20}$, linear mixed-effect model).

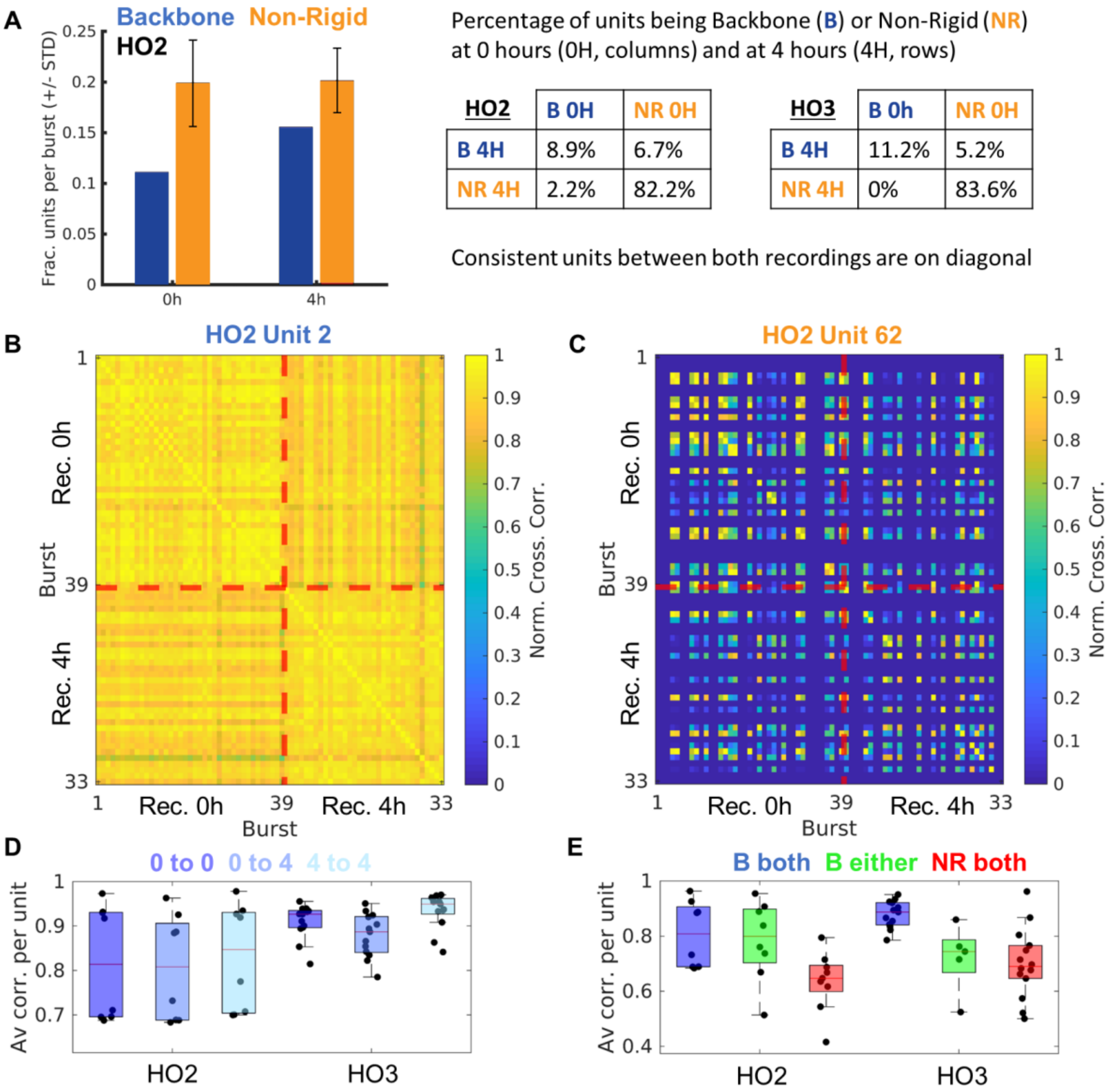


**Supplementary Figure 12: Backbone unit activity is stable over multiple hours.** (**A**) Left, fraction of backbone units and non-rigid units that fire at least 2 spikes per burst. Backbone units fire at least 2 spikes in all bursts and so the fraction of backbone units is the same for each burst. The fraction of non-rigid units that fire at least 2 spikes varies per burst, depicted by the error bars that represent the standard deviation over all bursts. The results are shown for two different recordings of the same organoid slice, that were made at a 4-hour interval. This data was recorded from a subset of human organoids used in this study (human organoid 2 (HO2) and organoid 3 (HO3) from Figs.1,3,5). Right, the majority of units that was detected as a backbone unit at time point 0-hour was also detected as backbone unit at time point 4-hour (top left) for both organoids. Meanwhile, the majority of units that were detected as non-rigid units at time point 0-hour were also detected as non-rigid units at time point 4-hour (bottom right) for both organoids. The percentage of backbone and non-rigid units are shown in the table on the right.(**B**) Pairwise burst to burst correlations for a representative example backbone unit highlight that the firing patterns of this unit are consistent between bursts in the 0-hour recording (top left), between bursts in the 4-hour recording (bottom right), and between bursts in the 0-hour and 4-hour recording (bottom left). (**C**) Pairwise burst-to-burst correlations for a representative example non-rigid unit highlight that the firing patterns of this unit are inconsistent between bursts in the 0-hour recording (top left), between bursts in the 4-hour recording (bottom right) and between bursts in the 0-hour and 4-hour recording (bottom left). (**D**) The average burst to burst correlation per backbone unit, computed between bursts that occurred both in the 0-hour recording (0 to 0), between bursts that occurred in the 0-hour and 4-hour recording (0 to 4), and between bursts that occurred both in the 4-hour recording (4 to 4) reflect the consistent firing pattern of backbone units over multiple hour time spans. (**E**) Comparing the average burst-to-burst correlations computed between burst pairs of which one burst occurred in the 0-hour recording and the other in the 4-hour recording show that the backbone units (**B**) are more consistent in their firing patterns over multiple hours than the non-rigid units (NR).

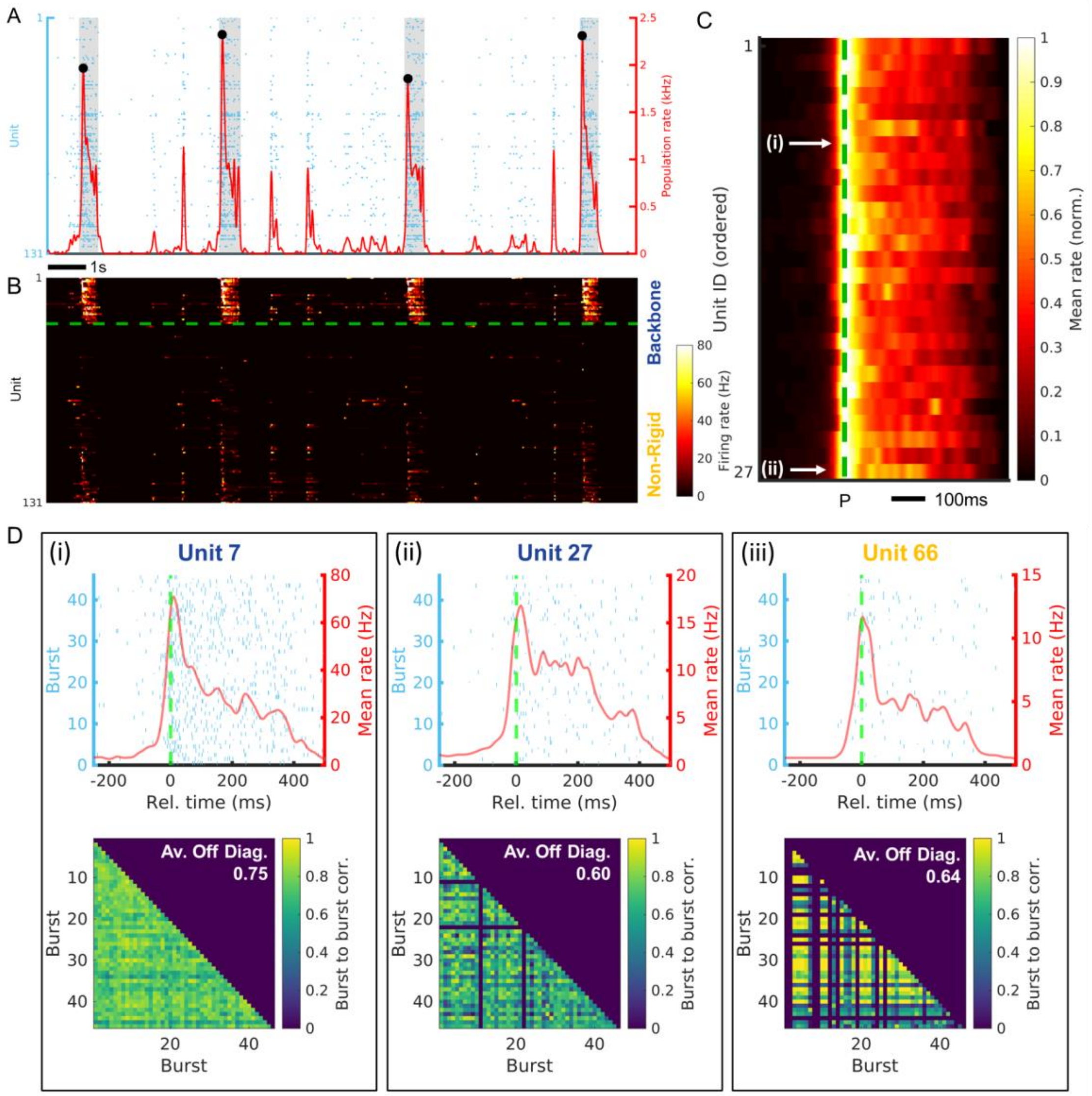


**Supplementary Figure 13. Sequential activations and burst-to-burst similarity are not present after shuffling.** (**A**) Same raster plot visualization as **Figure 1A** after shuffling. The population firing rate remains the same after shuffling and is shown by the red solid line.

Population bursts exist in the same frames after shuffling and are denoted by local maxima (black dots) that exceed 4x-RMS fluctuations in the population rate. The burst duration windows remain the same after shuffling and are marked by the shaded gray regions, which denote the interval in which the population rate remains above 10% of its peak value in the burst. The average firing rate per unit remains the same after shuffling. (**B**) Same instantaneous firing rate visualization as **Fig. 2B**. The same ordering is used as in **Fig. 2B**. (**C**) Same average burst peak centered firing rate visualization as Figure 2C after shuffling. The burst peak is indicated by the dotted line. The unit order is the same as Figure 2C. Note that the progressive increase in the firing rate peak time relative to the burst peak, as well as a spread in the active duration for units having their peak activity later in the burst are not present anymore after shuffling. The average firing rate is normalized per unit to aid in visual clarity. (**D**) Same burst-peak-centered spike times and pairwise burst-to-burst correlations as in **Fig. 2D** after shuffling. For (i) and (ii), the consistent firing patterns relative to the burst peak as exemplified in **Fig. 2D** are not present anymore and the average burst-to-burst correlation scores have decreased from 0.96 to 0.75 and from 0.82 to 0.60 respectively. Meanwhile, the average burst to burst correlation for the non-rigid unit exemplified in (iii) has increased from 0.51 to 0.64.

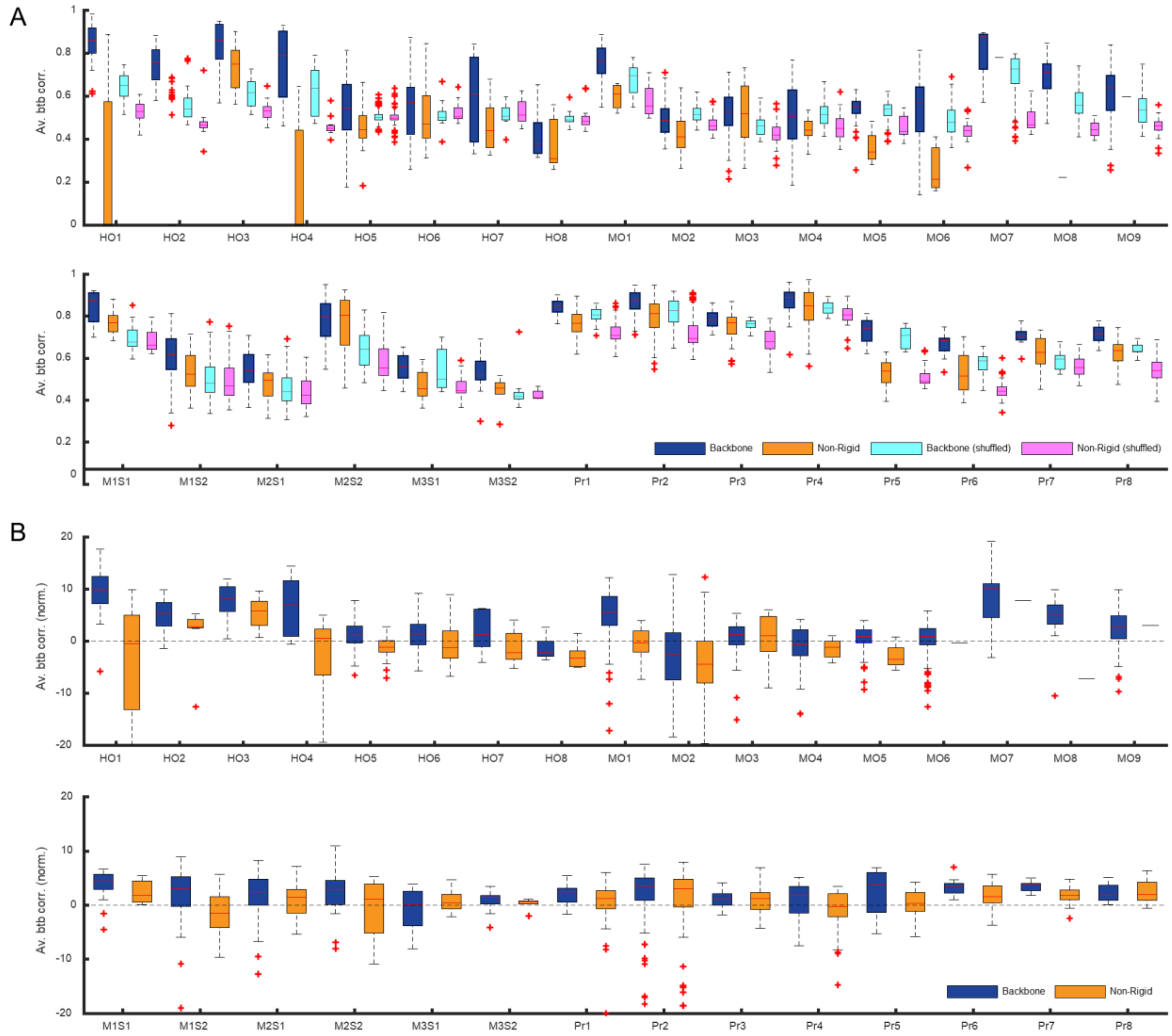


**Supplementary Figure 14. Backbone units have a higher average burst-to-burst correlation after firing rate normalization for organoids and murine neonatal slices.** (**A**) The distribution of burst-to-burst correlations per analyzed recording for backbone and non-rigid units separated. The results for applying the same analysis on shuffled data are included in magenta and cyan. The data is shuffled in a way that maintains the same average firing rate per unit and the same population rate per recording frame. In all cases, the backbone units have on average a higher correlation, also after shuffling. This indicates that the average firing rate differences by themselves affect the burst-to-burst correlation. HO = Human Organoid, MO = Murine Organoid, MS = Murine Slice, Pr = Murine Primary culture. (**B**) The distribution of burst-to-burst correlations per analyzed recording for backbone and non-rigid units separated after subtracting the score from the shuffled recording from the original score. In all organoid recordings and most murine neonatal slice recordings, the average burst-to-burst correlation after average rate normalization is still higher for the backbone units than for the non-rigid units. Meanwhile, this difference is not present anymore after normalization for the murine primary cultures. See **Fig. 7B** for the results of the statistical comparison.

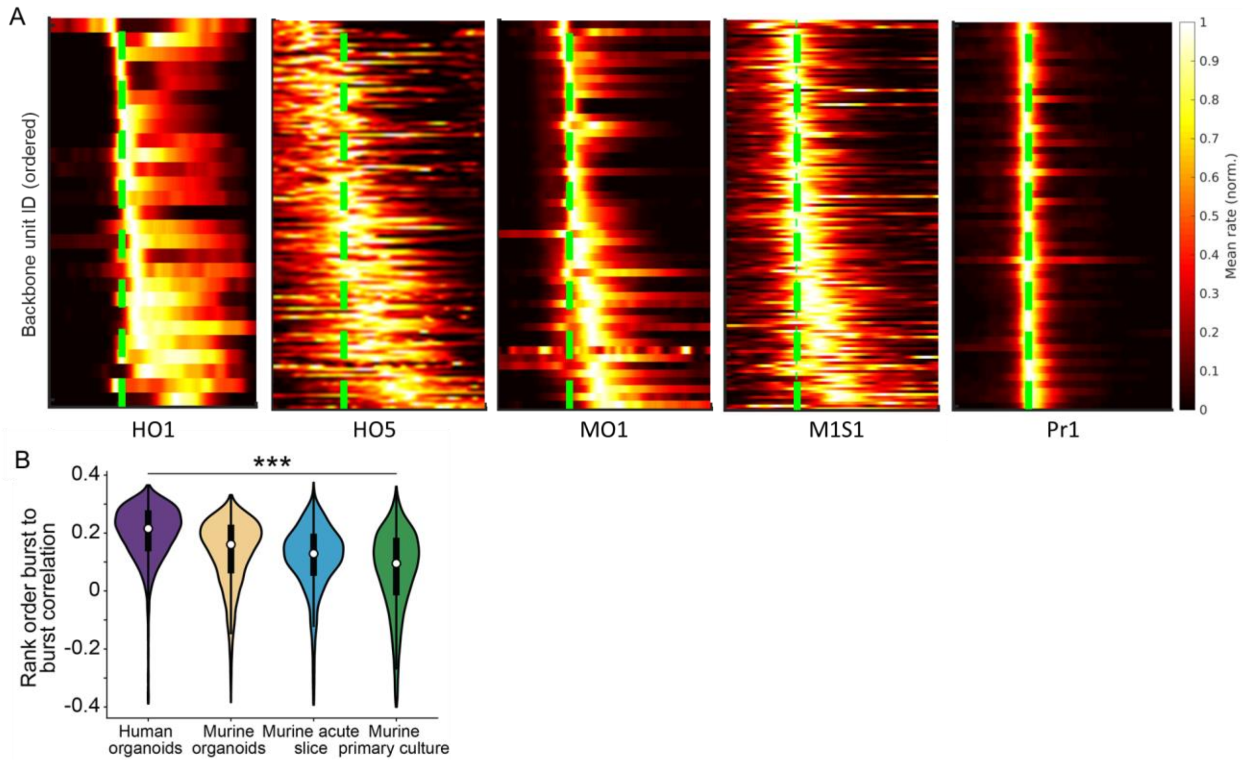


**Supplementary Figure 15. Rank order correlations. (A)** examples of sequential firing patterns visible in the burst peak centered average firing rates of backbone units in human organoids from Sharf *et al*. (HO1), human organoids from Alam El Din *et al*. (HO5), murine organoids (MO1) and murine brain slices (M1S1) but not in murine primary cultures (Pr1). **(B)** Normalized Spearman rank order correlations comparing the sequential order of backbone sequences for all burst pairs over the recordings for the 4 different model types. Correlation scores are normalized relative to shuffled spike matrices such that values above 0 indicate a more consistent backbone sequence than the shuffled data (Main effect of sample type P=0.004, Human organoids vs Murine primary cultures pairwise comparison P=0.0004, linear mixed-effect model with sample as grouping variable).

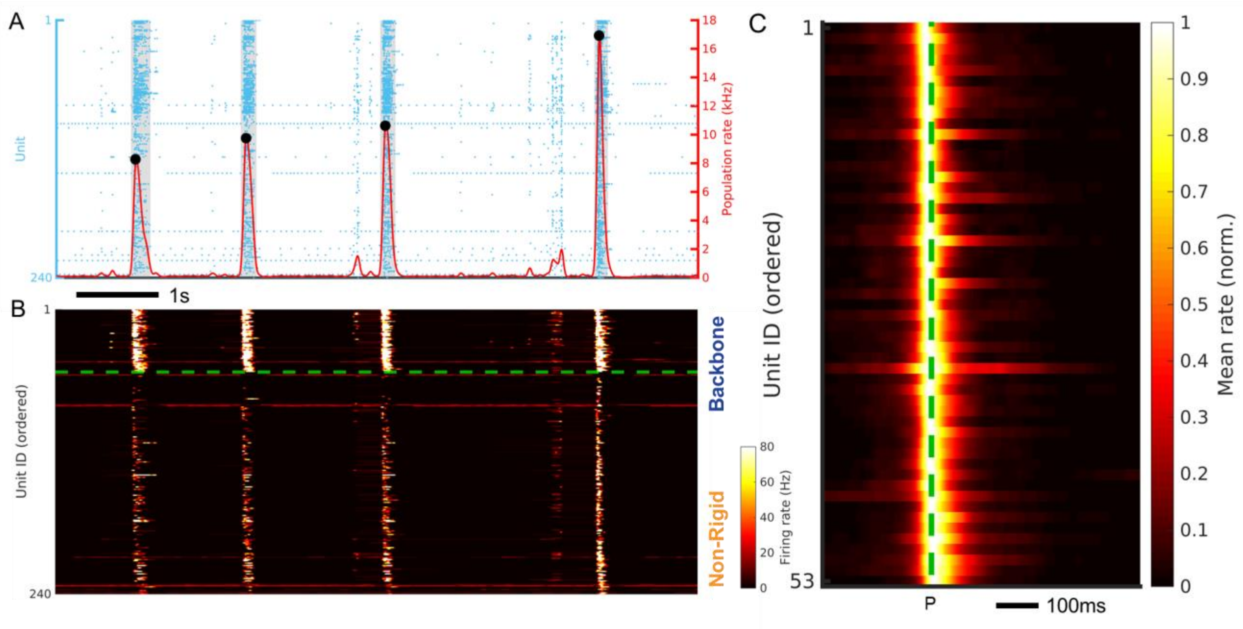


**Supplementary Figure 16: Intrinsic activity in murine primary cultures resembles the organoid results after shuffling.**

(**A**) Raster plot visualization of single-unit spiking (blue dots) measured across a 2D murine primary culture from Yuan *et al*. (Pr1) recorded on a high-density microelectrode array. The population firing rate is shown by the red solid line. Population bursts are marked by sharp increases in the population rate. Burst peak events are denoted by local maxima (black dots) that exceed 4x-RMS fluctuations in the population rate. The shaded gray regions denote the burst duration window as defined by the time interval in which the population rate remains above 10% of its peak value in the burst. (**B**) The instantaneous firing rate of single-unit activity from panel A after reordering. The backbone units are plotted above the dashed line while non-rigid units are plotted below the dashed line. In each category, units are ordered based on their median firing rate peak time relative to the burst peak, considered over all bursts in the recording. (**C**) The average burst peak centered firing rate measured across all burst events for the example recording of which part is shown in **A**. The burst peak is indicated by the dotted line. The unit order is the same as **B**. Note that the progressive increase in the firing rate peak time relative to the burst peak, as well as a spread in the active duration for units having their peak activity later in the burst are not present in the murine primary recording, similar to the organoid data after shuffling as shown in **Supplementary Fig. 13**. The average firing rate is normalized per unit to aid in visual clarity.

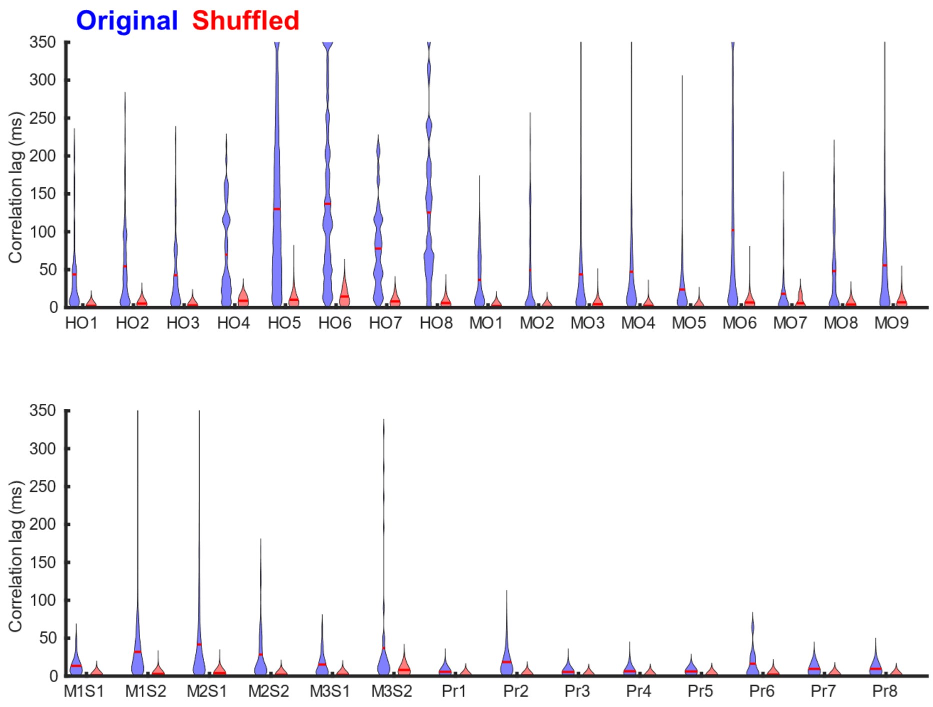


**Supplementary Figure 17. Correlation lag times.** Lag times corresponding to the maximum correlation when applying a cross correlation to the single unit firing rates over all pairs of backbone units. The maximum lag time was set to 350ms, the median duration of the backbone period over all 3D models. The same computations were performed on 100 different shuffled spike matrices and the average absolute correlation lags over all 100 shuffled datasets are reported as well. See **Fig. 7D** for the results of the statistical comparison.

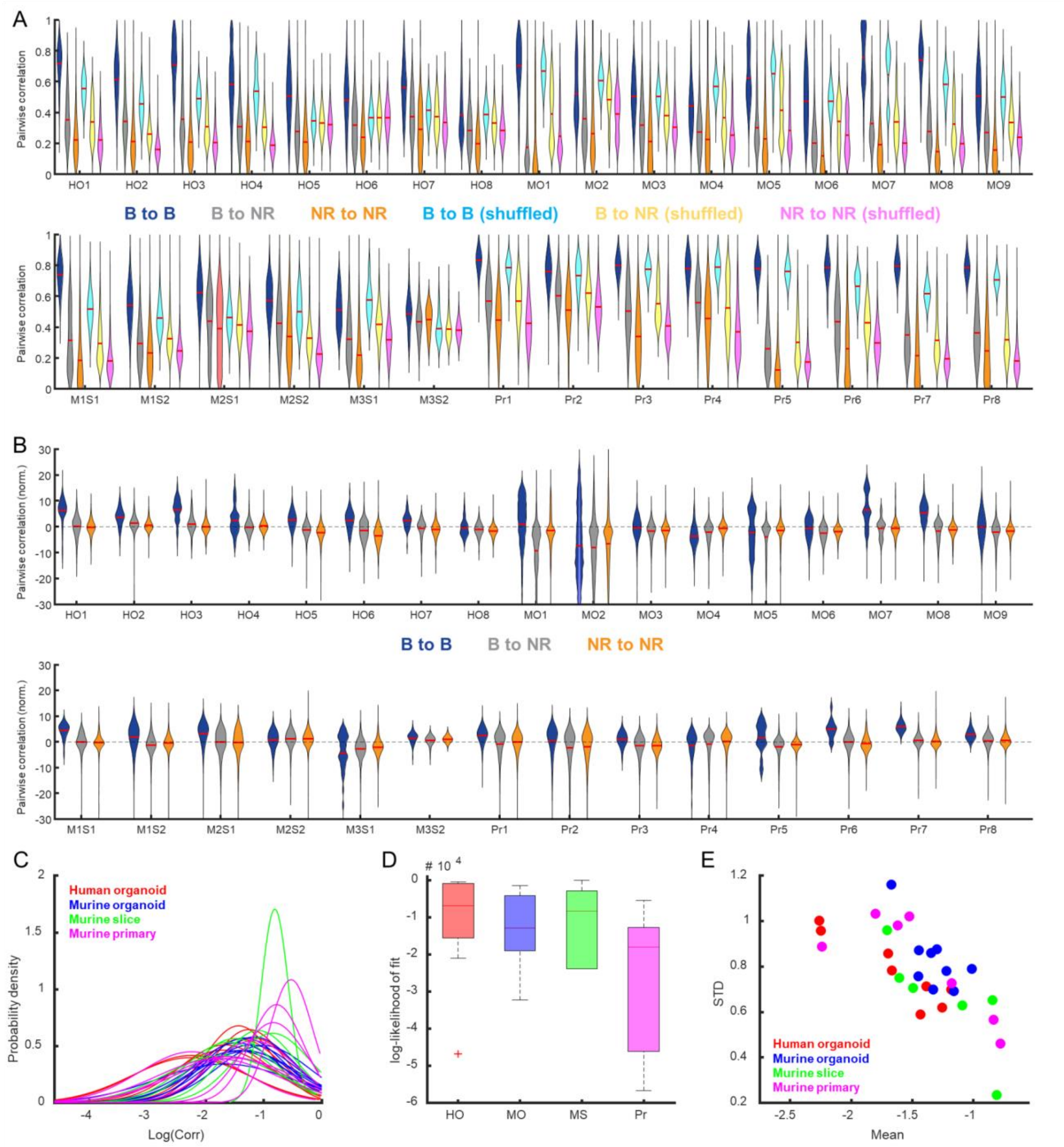


**Supplementary Figure 18. Backbone unit pairs have a higher pairwise correlation after firing rate normalization for organoids and murine neonatal slices.** (**A**) The distribution of pairwise correlations per analyzed recording for backbone unit pairs, backbone and non-rigid unit combinations and non-rigid unit pairs separated. The results for applying the same analysis on shuffled data are included in magenta, yellow and cyan. The data is shuffled in a way that maintains the same average firing rate per unit and the same population rate per recording frame. In all cases, the backbone unit pairs have on average a higher pairwise correlation compared to other unit type pairs, also after shuffling. This indicates that the average firing rate differences by themselves affect the pairwise correlations. B = Backbone, NR = Non-Rigid, HO = Human

Organoid, MO = Murine Organoid, MS = Murine Slice, Pr = Murine Primary culture. (**B**) The distribution of pairwise correlations per analyzed recording for backbone unit pairs, backbone and non-rigid unit combinations and non-rigid unit pairs separated after subtracting the score from the shuffled recording from the original score. In all organoid recordings and most murine neonatal slice recordings, the pairwise correlations after average rate normalization are still higher for the backbone unit pairs than for the other unit pairs. Meanwhile, this difference is not present anymore after normalization for the murine primary cultures. See **Fig. 7C** for the results of the statistical comparison. (**C**) Normal distributions fitted to the distributions of the log of the pairwise correlations. (D) Log-likelihood for the fits in **C**. (**E**) The mean and STD for the fits in **C**.

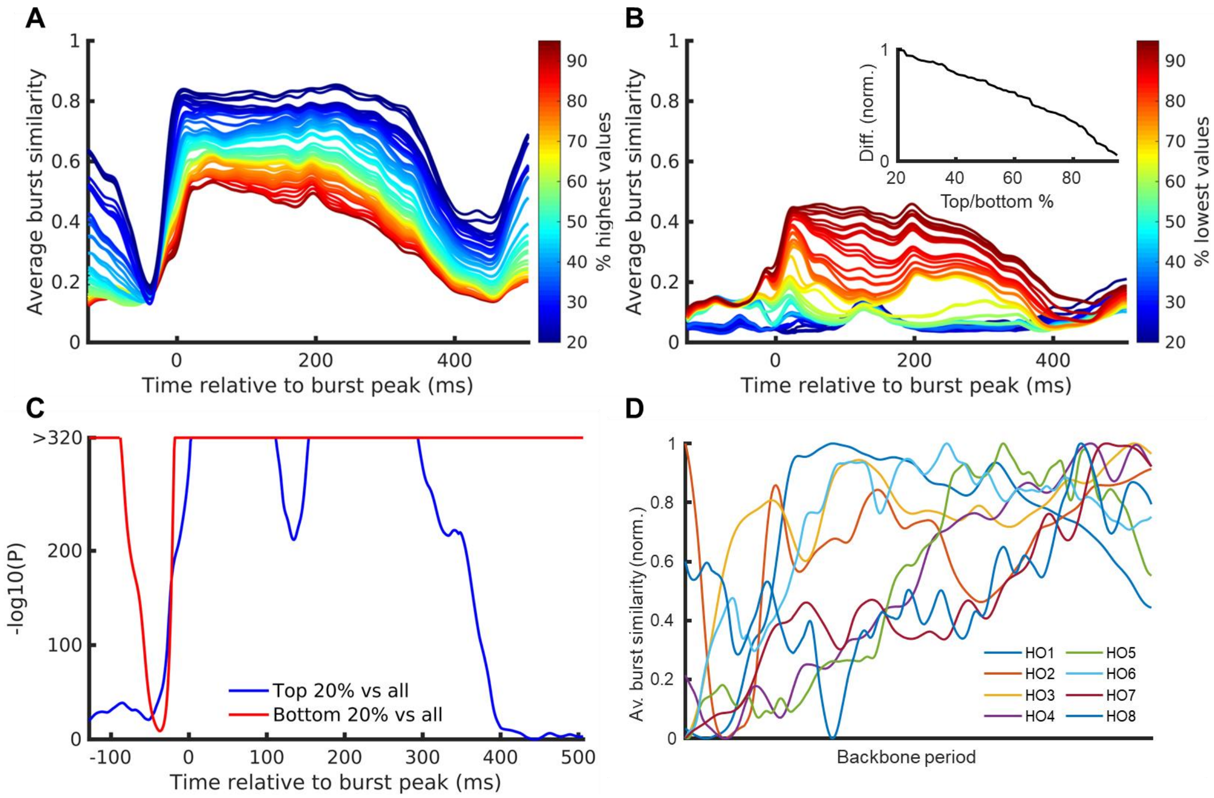


**Supplementary Figure 19. Burst similarity for correlated units.** (**A**) The top 20th percentile of units based on their average pairwise correlation generated higher cosine similarity scores compared to larger percentile ranges, highlighting that the burst similarity score is driven by the most correlated units. (**B**) Greater than 80% of the units with the lowest average pairwise correlation was required to get similar burst similarity score traces as all units together. The burst similarity score increase during the backbone period was not present when only the 20-40% units with the lowest average pairwise correlation was considered. The inset shows the difference in the area under the curve for the top x% of units compared to the bottom x% of units. (**C**) The -log10(*P*) values of the paired sample t-tests comparing the burst similarity scores of the top or bottom 20% units based on their average pairwise correlation to the burst similarity scores computed over all units. A paired sample t-test is performed at every frame relative to the burst peak, comparing the distribution of burst similarity scores for all possible burst pairs between the two unit selections. (**D**) Average burst similarity computed over all units for each of the 8 organoids over their respective backbone periods. A backbone period is defined as the time of the first median firing-rate peak time until the last median firing rate peak time relative to the burst peak over all backbone units. The burst similarity scores within the backbone period are normalized between 0 and 1 for each organoid. Note the sharp increase from the start of the backbone period for all organoids.

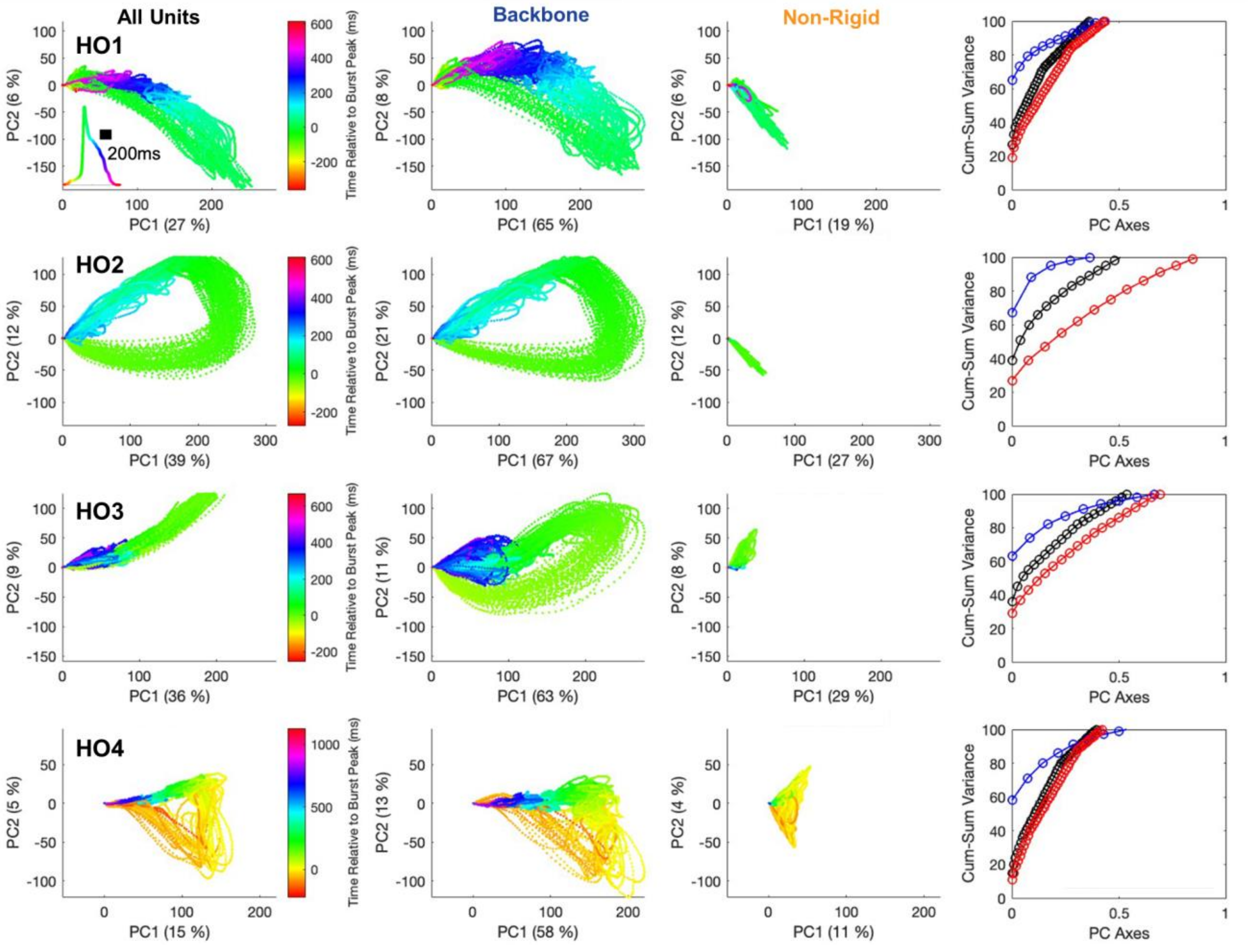


**Supplementary Figure 20. PCA manifolds for different organoid recordings.** Every row contains data from a different organoid. A PCA was performed on the firing rates per single unit for all units in the recording combined. Only spikes that occurred during bursts were included in the firing rate computations. Each dot represents a recording frame and is colored by the time point relative to the closest burst peak. First column, population activity projected onto its first two principal components. Note the consistent circular trajectory reflecting the burst manifold. The inset shows the burst-peak-centered average population rate colored in the same way as the PCA trajectories to indicate which parts of the burst correspond to the different coloring. Second column, same as first column but only including the backbone units. Note the similarity in the low dimensional manifold representations between the first and second column, indicating the strong contribution of the backbone units to the low dimensional activity of the whole population. Third column, same as first column but only including the non-rigid units. The low-dimensional manifold is not present anymore and the variance explained by the first to principal components is notably lower, reflecting the higher dimensional activity of the non-rigid units.

Last column, the cumulative sum of the variance explained per principal component as a function of the number of first principal components included in the sum. The explained variance for backbone units only (blue) required fewer PCs compared to all units (black). Meanwhile, the opposite is true for the non-rigid units only (red).

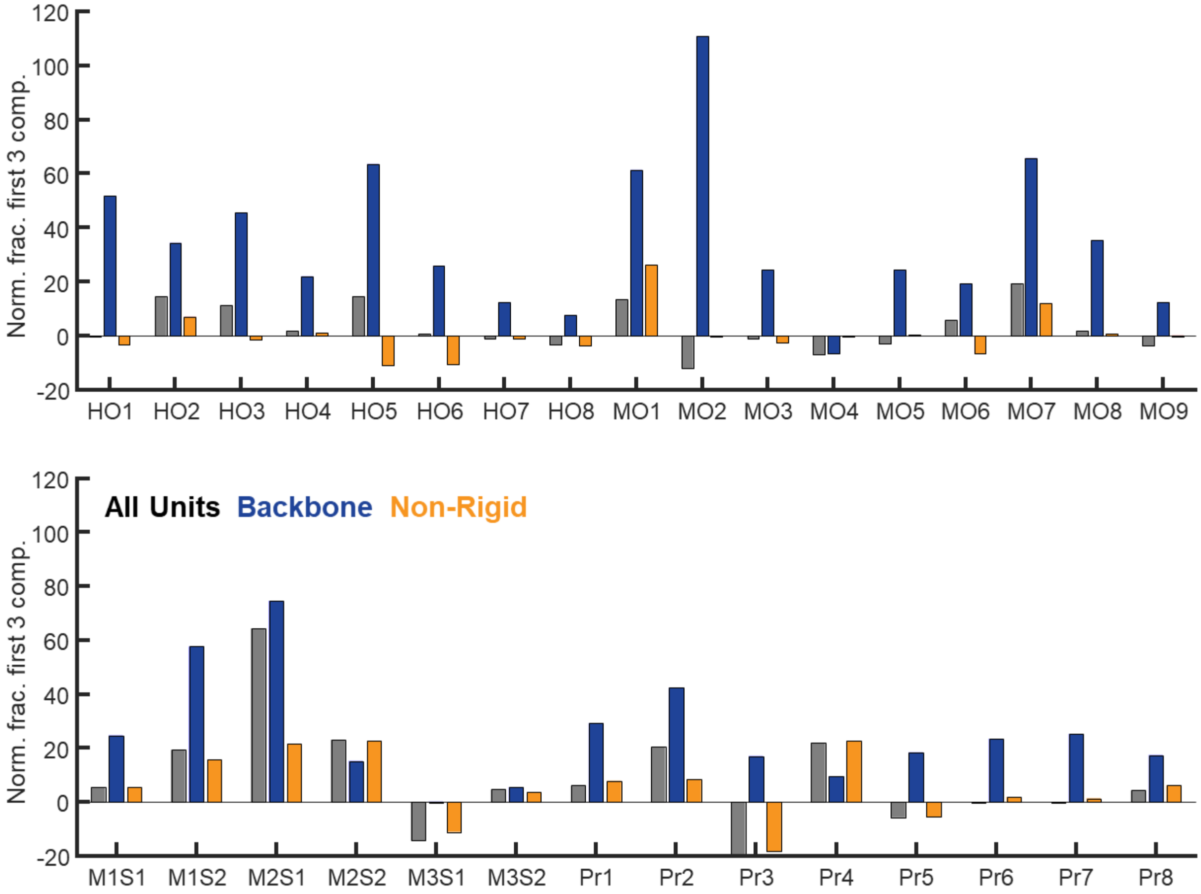


**Supplementary Figure 21. Normalized sum of explained variance of first 3 components of PCA manifold.** The normalized fraction of the variance explained, summed over the first three principal components for the PCA manifolds per recording. To account for differences in the total number of principal components per category, the summed explained variance for the first three principal components is divided by the summed explained variance of the first *X* principal components, where *X* is the lowest number of total principal components from the three categories: all units, backbone and non-rigid. *X* is defined separately per recording due to variation in the number of detected units. This summed variance is computed for the original data and the shuffled data after which the shuffled results are subtracted from the original results to get the final value. See **Fig. 7E** for the results of the statistical comparison.

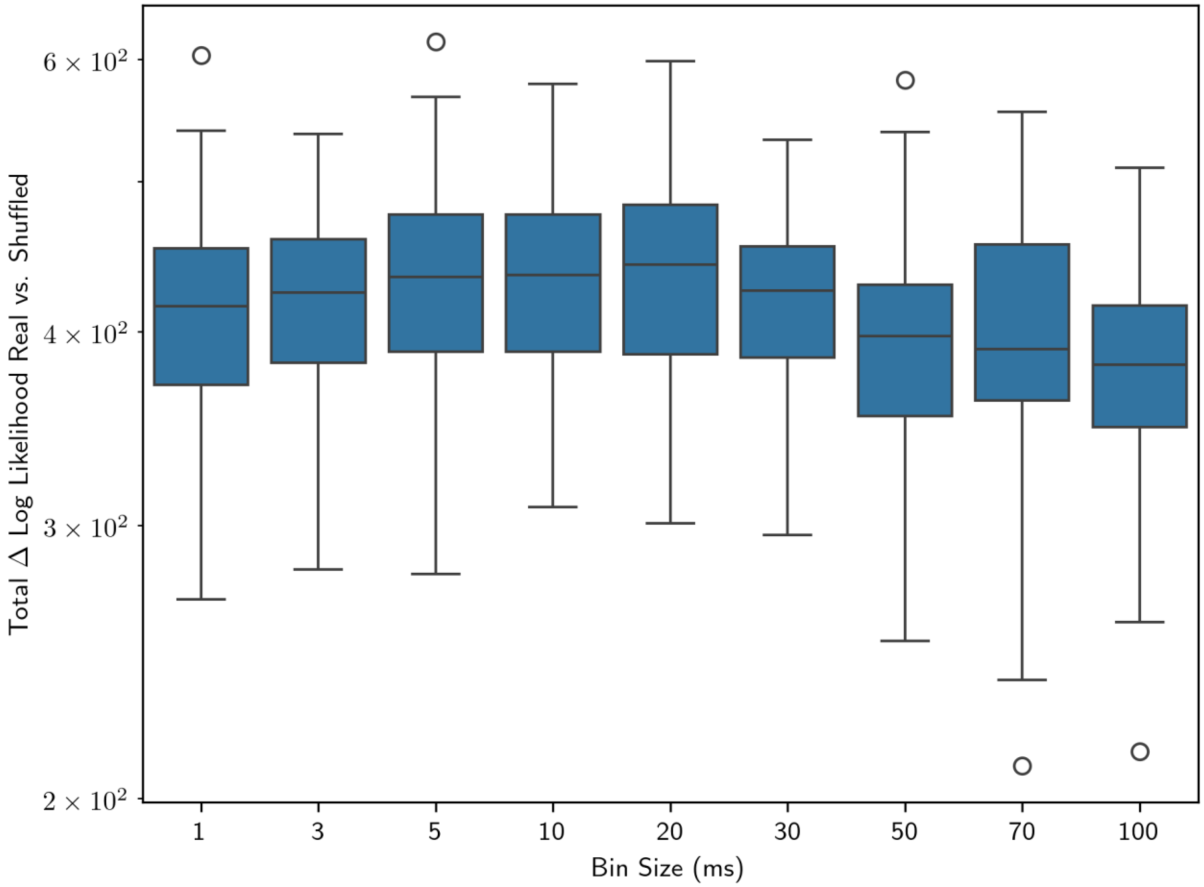


**Supplementary Figure 22. Model performance with bin size**. Distribution of the difference in posterior log likelihood between held-out validation data and shuffled data across 5 random cross-validation folds are shown here for Organoid 1. The number of states ranges from 10 to 30, for a total of 31 data points per bin size. Performance is relatively insensitive to bin size, just as it is insensitive to model size but depends substantially on the biological sample itself (Supplementary Figures 16 and 17). Note that in this and other supplementary figures depicting this metric, the difference in log likelihood is itself shown on a log scale.

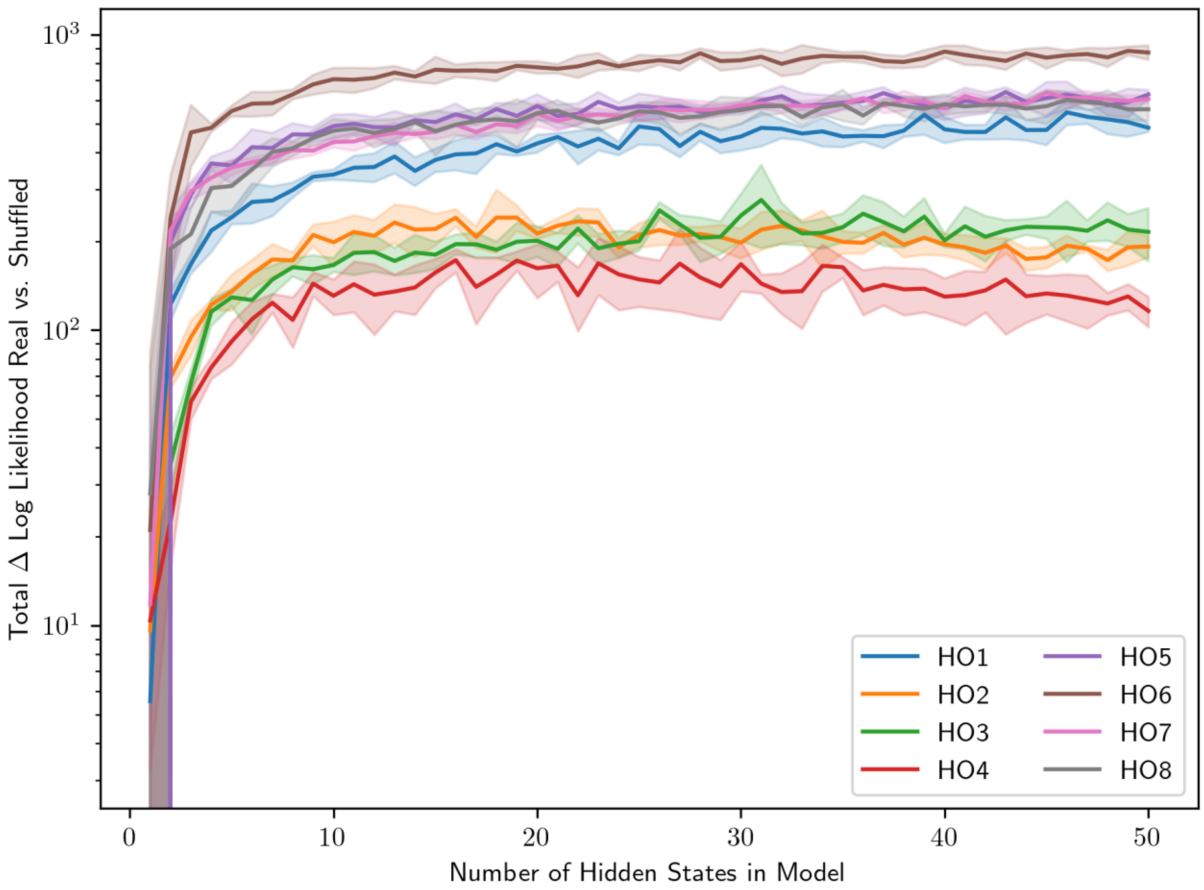


**Supplementary Figure 23. posterior log likelihood plateaus around 10 hidden states**
Difference in posterior log likelihood between the held-out validation set and randomized data, for models with 30 ms time bins, as a function of the number of hidden states. Curves indicate mean values, with shaded area for ±1 STD, as calculated across 5 random cross-validation folds as described in Methods. The posterior log likelihood of each human brain organoid recording plateaus at a different value, but this is highly consistent across multiple training runs and multiple CV folds.

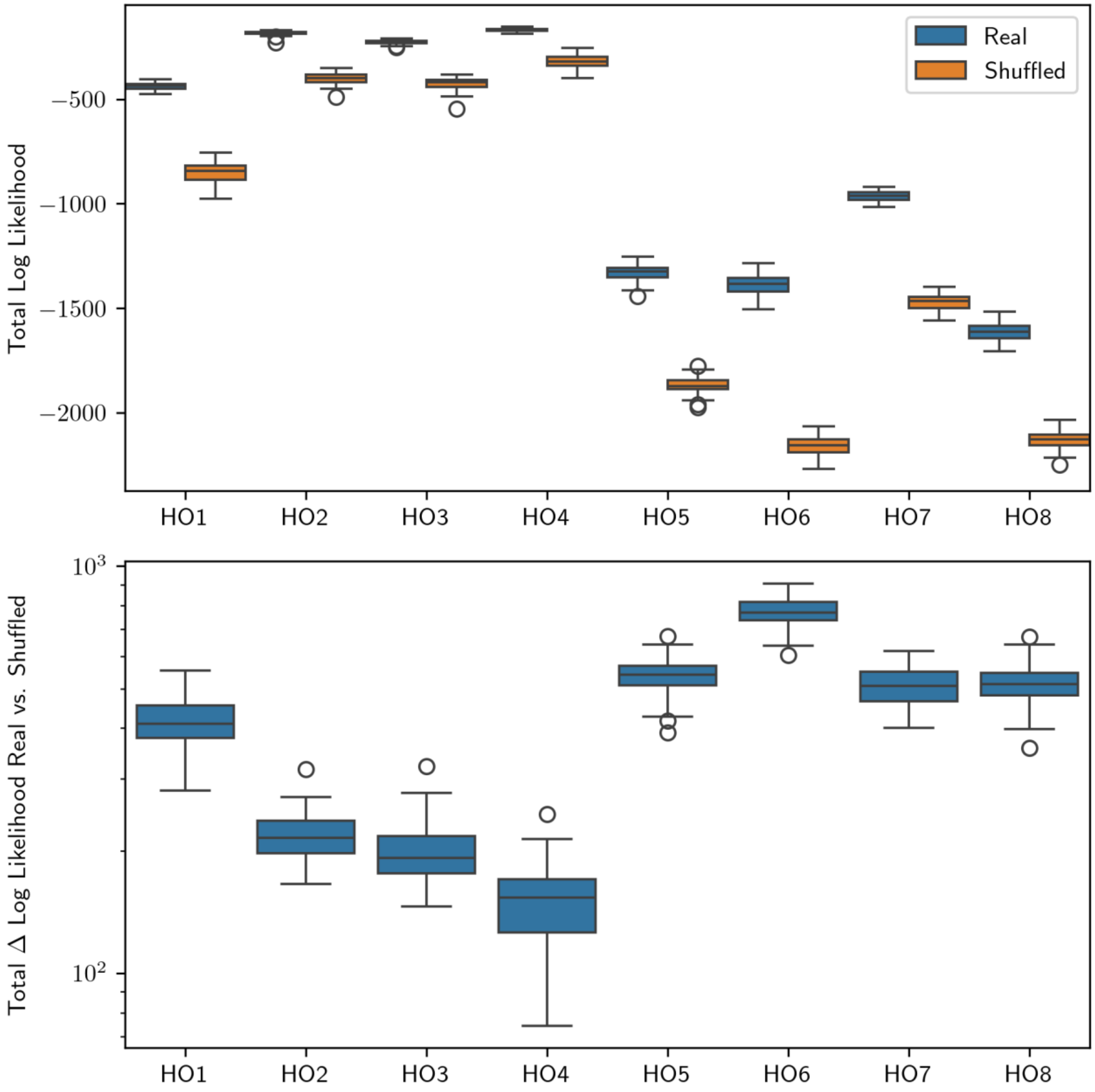


**Supplementary Figure 24: Log likelihood is consistently higher for real compared to randomized data.** Difference in posterior log likelihood between real and randomized data as a distribution across 5 cross-validation folds for models trained on 30 ms time-binned rasters of each of the human brain organoid recordings, with the number of hidden states ($K$) ranging from 10 to 30. The figure compares the value of the log likelihood between the real data and the shuffled data control (top), as well showing the distribution per organoid of the difference in log likelihood for each value of $K$ (bottom). Log likelihood is always greater for real than for randomized data, with no overlap between the distributions for real and shuffled data. However, the value, as well as the magnitude of the difference, varies systematically between recordings.

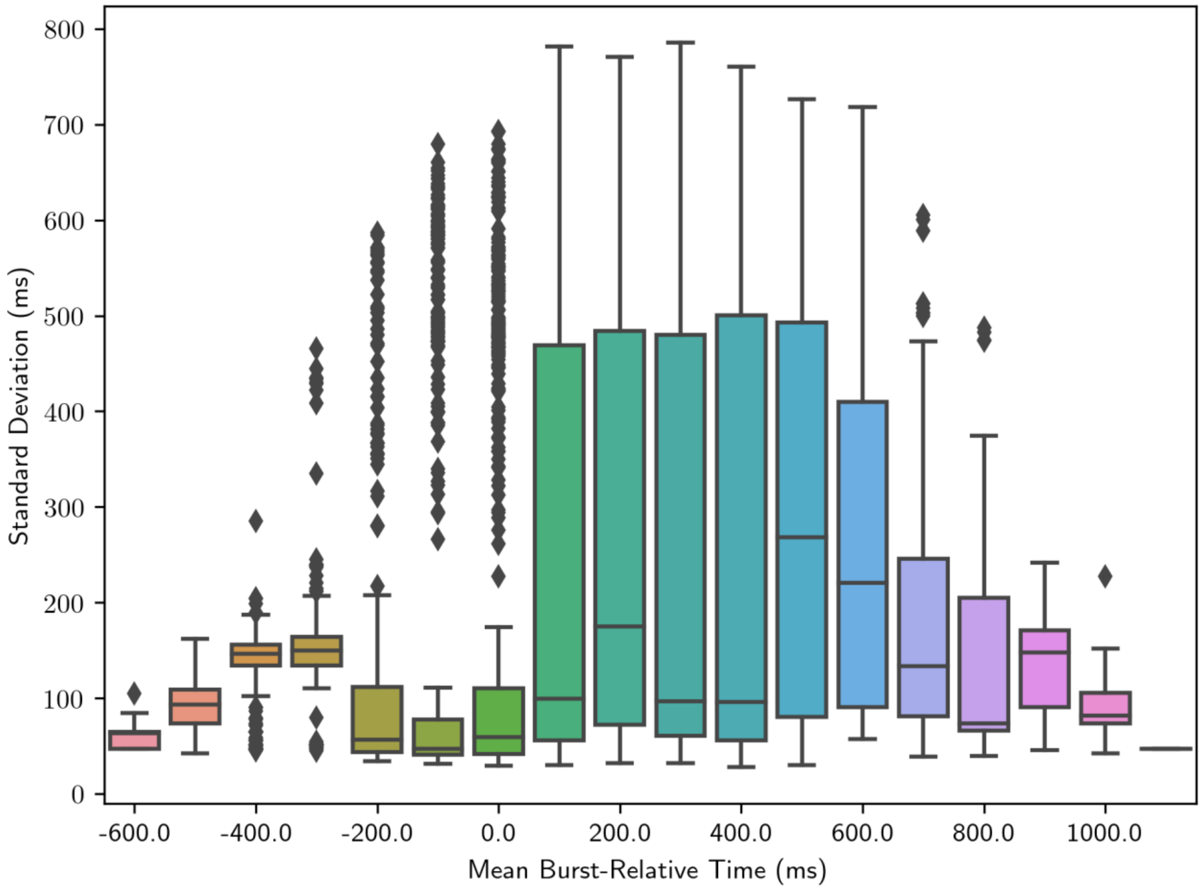


**Supplementary Figure 25. Later hidden states have greater temporal spread within a burst**. We calculated the histogram of burst-relative times of realizations of each hidden state of each HMM fitted to each organoid recording with bin size $T = 30$ ms (see Methods), and fitted a Gaussian distribution to each histogram, parametrized by the mean and standard deviation of burst-relative times. We attempted to restrict our attention to states which are well described by the mean/STD characterization (as opposed to, e.g., having a multimodal distribution with significant weight both before and after the burst). To do this, we selected hidden states via a chi-square test, including only those which failed to reject the Gaussian fit at the 1% significance level (i.e., all states with $P > 1\%$). The selected states were then grouped according to their mean occurrence time with 100 ms resolution.

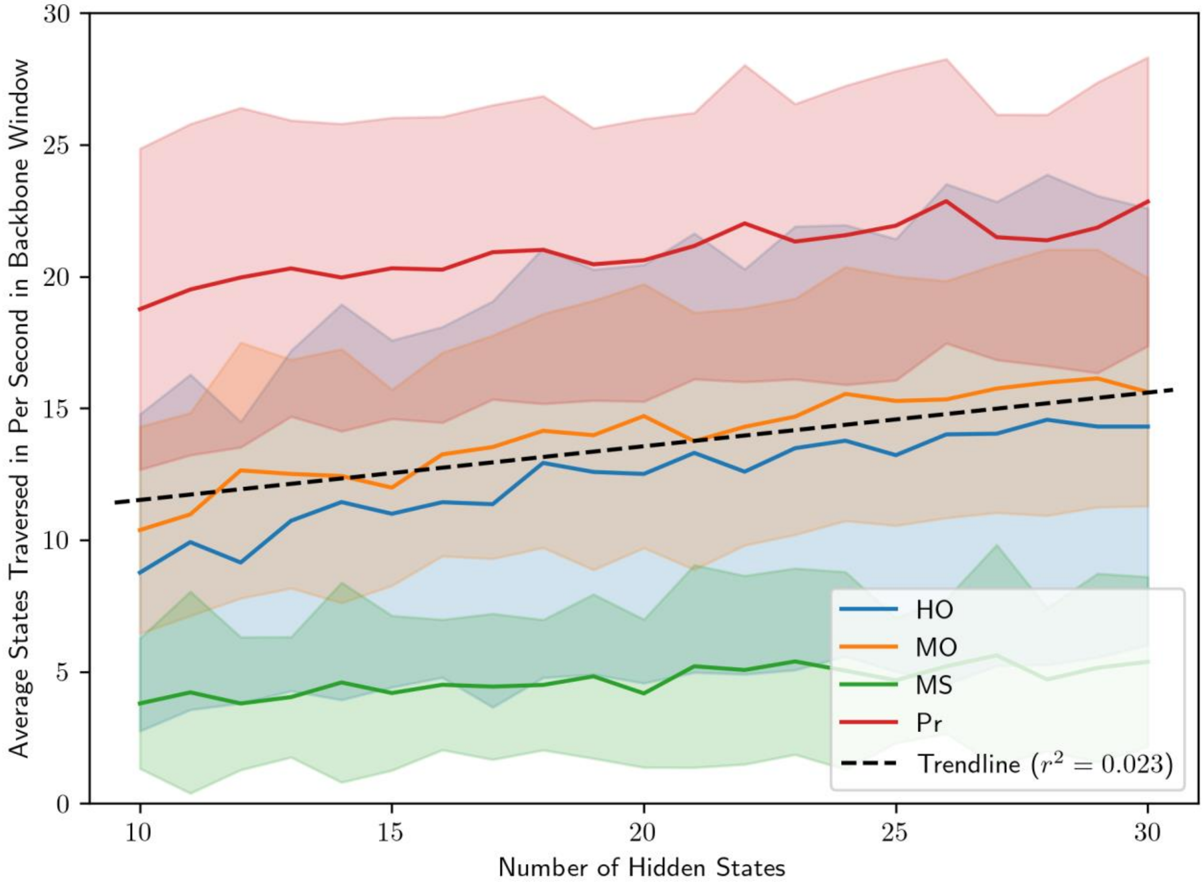


**Supplementary Figure 26. State traversal rate varies little with the number of hidden states.** For each hidden Markov model, fitted to a given experiment, we averaged across bursts the number of *distinct* hidden states traversed during the backbone interval (see Methods) divided by its duration. An overall trendline is drawn indicating the effect of the number of hidden states on the state traversal rate. Although it is statistically significant (*P* < 0.01%), the effect size is small in comparison to the variability within and between samples.

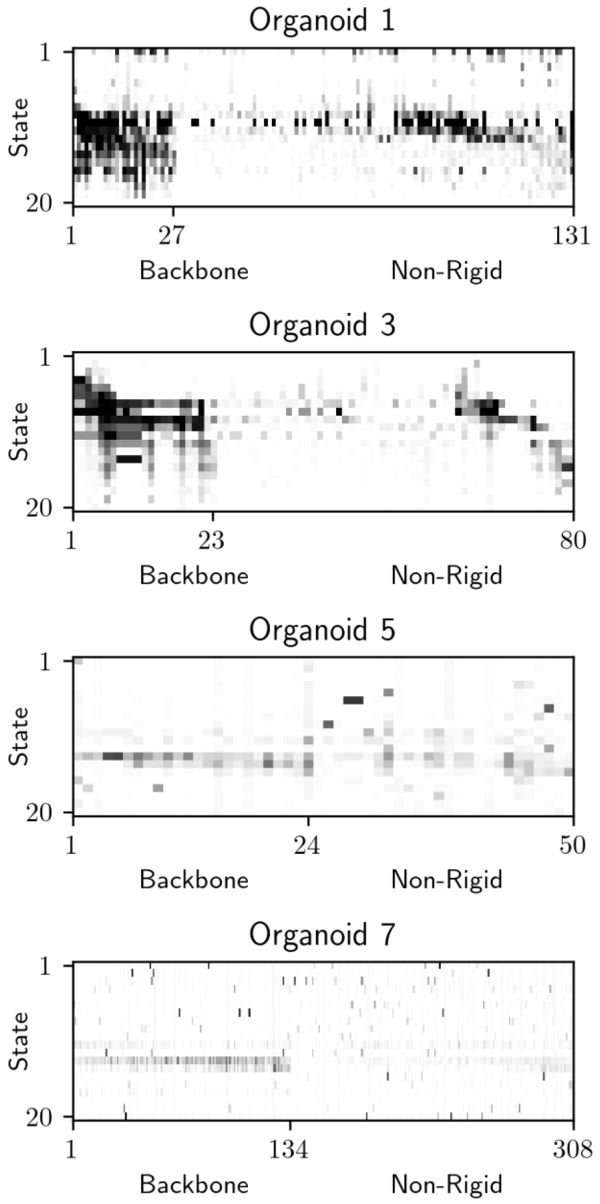


**Supplementary Figure 27. Backbone and non-rigid units differ in their consistency by state.** Each panel represents, for one brain organoid recording, the probability that each unit fires at least once in each of the 20 hidden states. The color scale as well as the meanings of all quantities and labels are identical to **Fig. 5D**.

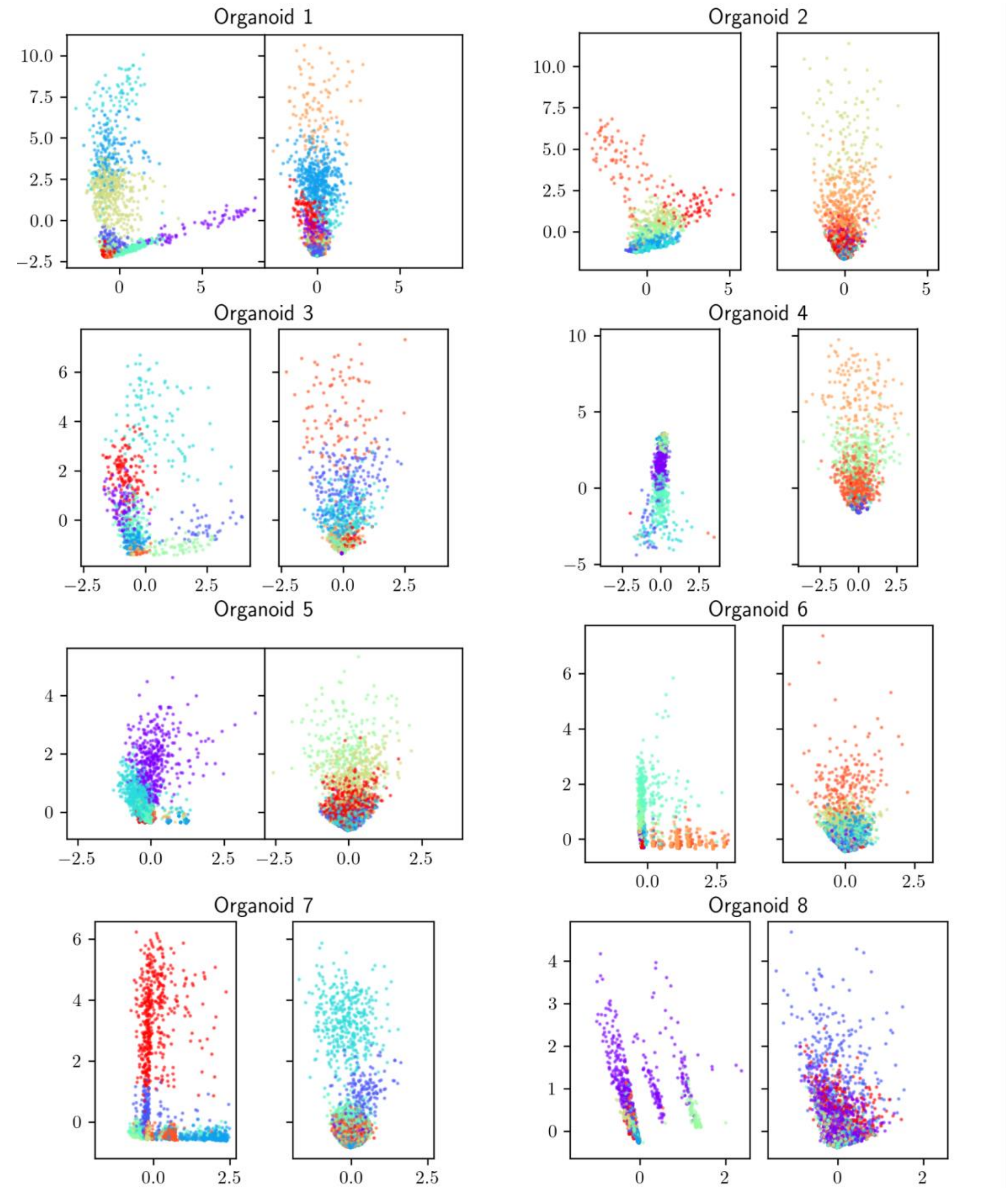


**Supplementary Figure 28. Randomized but not real data is explained by population rate**
HMM-based reduced-dimension view of time-binned spike matrix data (see *Methods: dimensionality of a fitted HMM*), with points colored according to the most probable hidden state according to a trained HMM with $K = 10$ hidden states. For each organoid, the left panel shows real data and the right panel shows randomized data. Note that in each case, for real data, hidden states clearly occupy different regions in the two-dimensional space, whereas they are predominantly vertically stratified in the randomized data. This vertical stratification must correspond at least approximately to the population rate, as population rate is the only temporally varying information left intact by randomization.

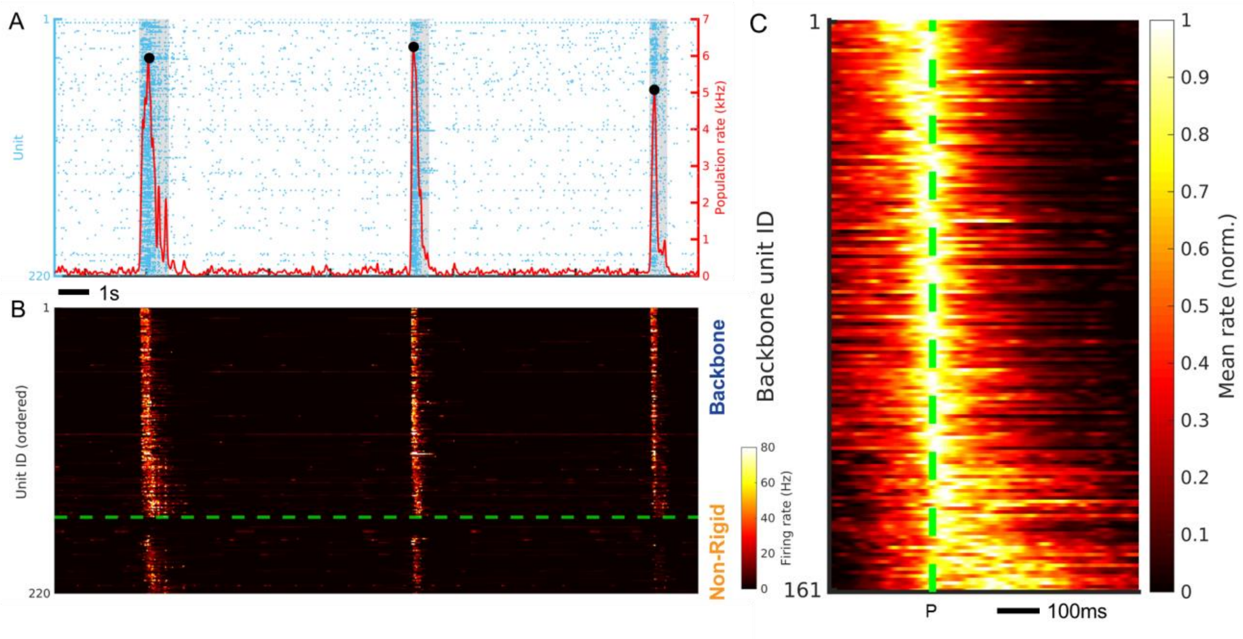


**Supplementary Figure 29: Consistent results in brain slices from different animals.** (**A**) Raster plot visualization of single-unit spiking (blue dots) measured across the surface of a murine neonatal cortical slice from a different animal (M3S1) dissected at P13, positioned on top of the same type of microelectrode array as used for the recordings included in the main figures. The population firing rate is shown by the red solid line. Population bursts are marked by sharp increases in the population rate. Burst peak events are denoted by local maxima (black dots) that exceed 4x-RMS fluctuations in the population rate. The shaded gray regions denote the burst duration window as defined by the time interval in which the population rate remains above 10% of its peak value in the burst. (**B**) The instantaneous firing rate of single-unit activity from panel A after reordering. The backbone units are plotted above the dashed line while non-rigid units are plotted below the dashed line. In each category, units are ordered based on their median firing rate peak time relative to the burst peak, considered over all bursts in the recording. (**C**) The average burst peak centered firing rate measured across all burst events for the example recording of which part is shown in **A**. The burst peak is indicated by the dotted line. The unit order is the same as **B**. Note the progressive increase in the firing rate peak time relative to the burst peak, as well as a spread in the active duration for units having their peak activity later in the burst. The average firing rate is normalized per unit to aid in visual clarity.

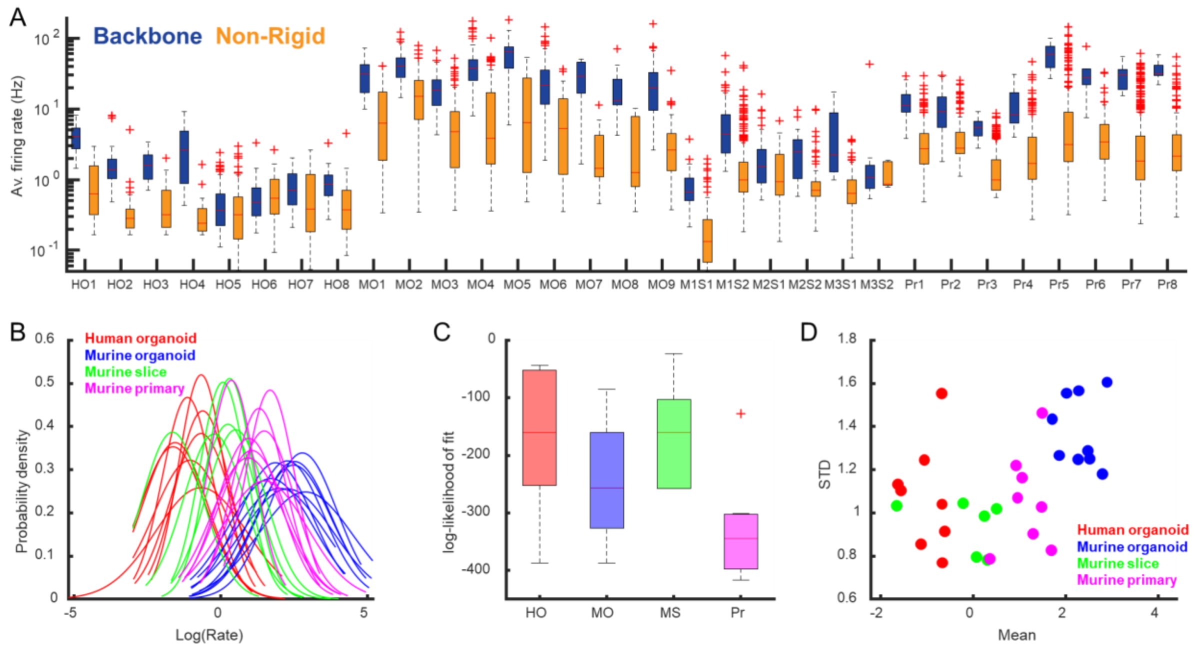


**Supplementary Figure 30. Backbone units have a higher average firing rate in each individual recording.** (**A**) The distribution of average firing rates per analyzed recording for backbone and non-rigid units separated. In all cases, the backbone units populate the tail of the skewed average firing rate distributions. Or = organoid, MS = Murine Slice, Pr = Murine Primary culture. See Fig. 7A for the results of the statistical comparison. (**B**) Normal distributions fitted to the distributions of the log of the firing rate per unit for all recordings. (**C**) Log-likelihood for the fits in **B**. (**D**) The mean and STD for the fits in **B**.

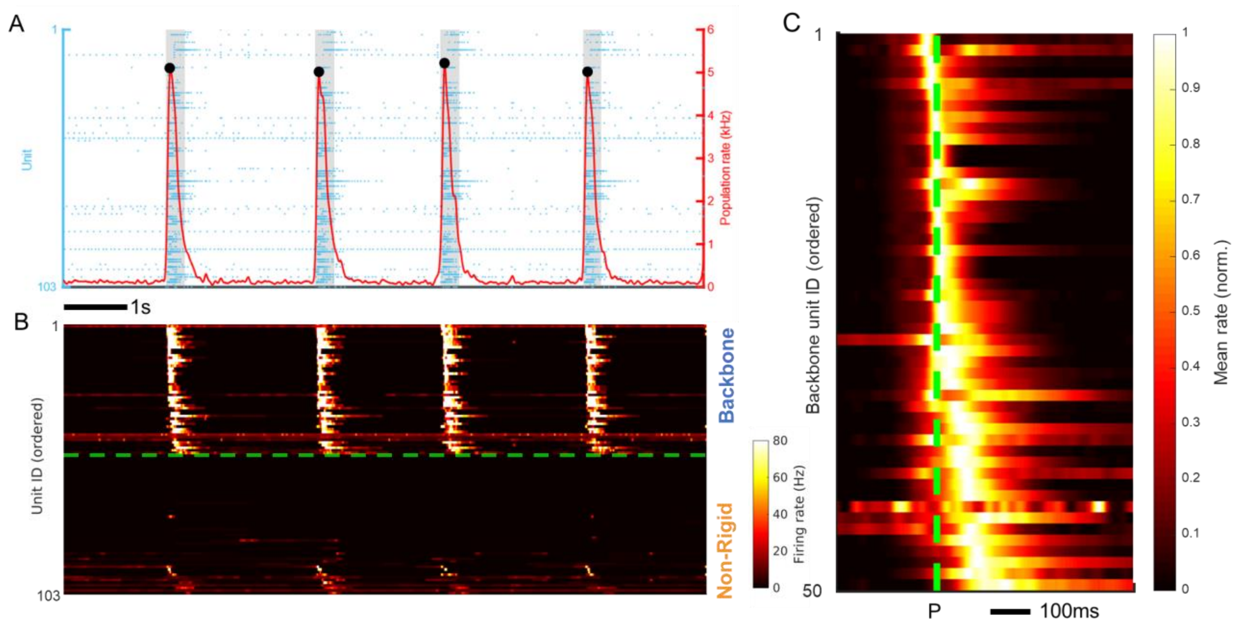


**Supplementary Figure 31. Backbone sequences in spontaneous population bursts in murine organoids.** (**A**) Raster plot visualization of single-unit spiking (blue dots) measured across the surface of a murine organoid (MO1) recorded at 42 DIV, positioned on top of the same type of microelectrode array as used for the recordings included in the main figures. The population firing rate is shown by the red solid line. Population bursts are marked by sharp increases in the population rate. Burst peak events are denoted by local maxima (black dots) that exceed 4x-RMS fluctuations in the population rate. The shaded gray regions denote the burst duration window as defined by the time interval in which the population rate remains above 10% of its peak value in the burst. (**B**) The instantaneous firing rate of single-unit activity from panel A after reordering. The backbone units are plotted above the dashed line while non-rigid units are plotted below the dashed line. In each category, units are ordered based on their median firing rate peak time relative to the burst peak, considered over all bursts in the recording. (**C**) The average burst peak centered firing rate measured across all burst events for the example recording of which part is shown in **A**. The burst peak is indicated by the dotted line. The unit order is the same as **B**. Note the progressive increase in the firing rate peak time relative to the burst peak, as well as a spread in the active duration for units having their peak activity later in the burst. The average firing rate is normalized per unit to aid in visual clarity.

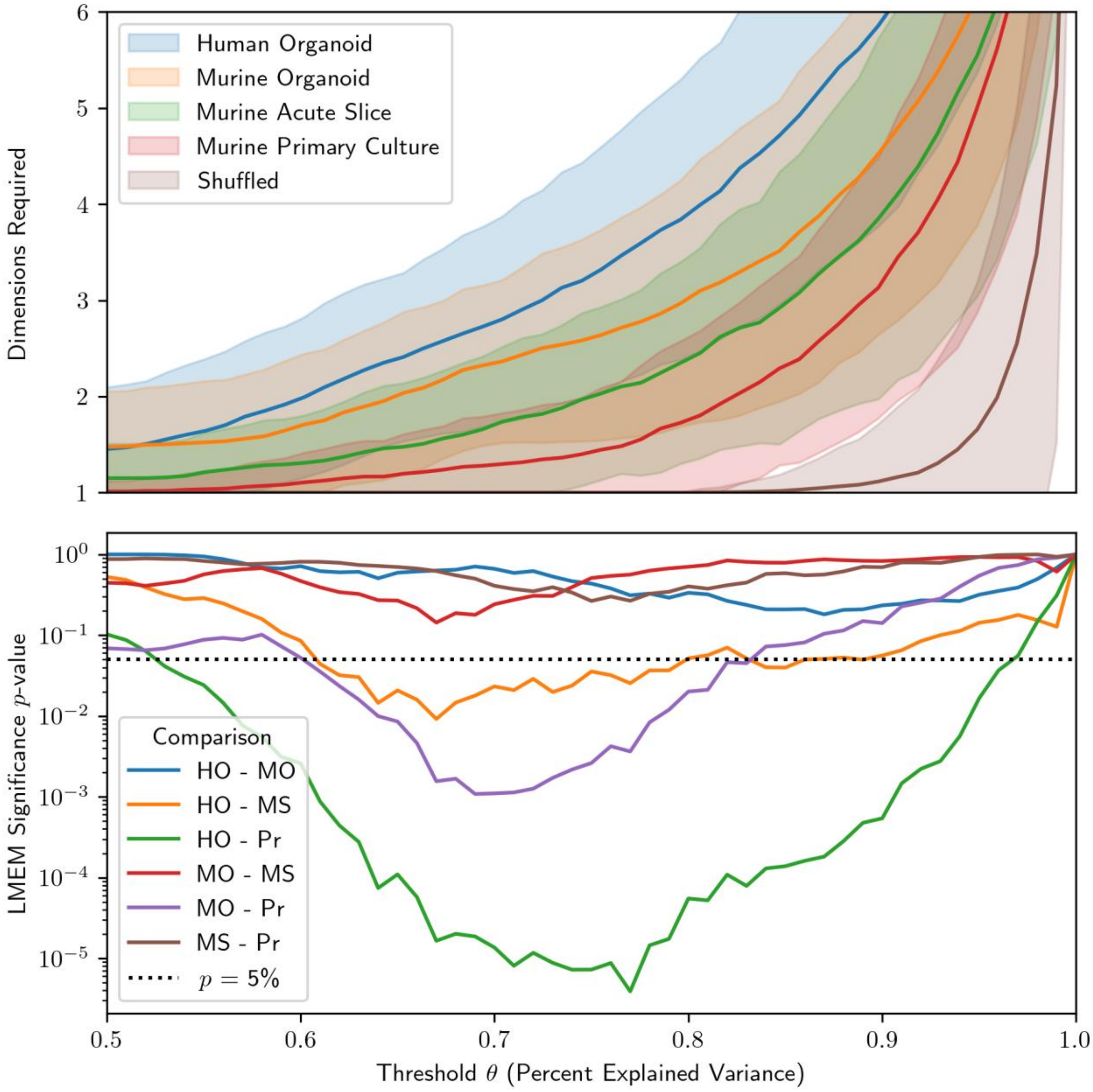


**Supplementary Figure 32. Threshold Dependence of Figure 7F.**
Top: mean ± one standard deviation of the dimensions required to explain a given fraction $\theta$ of the variance between HMM states, as a function of $\theta$. Bottom: statistical significance of the difference between groups in the top panel, according to a generalized linear mixed-effects model (see *Methods: Dimensionality of a Fitted HMM*).

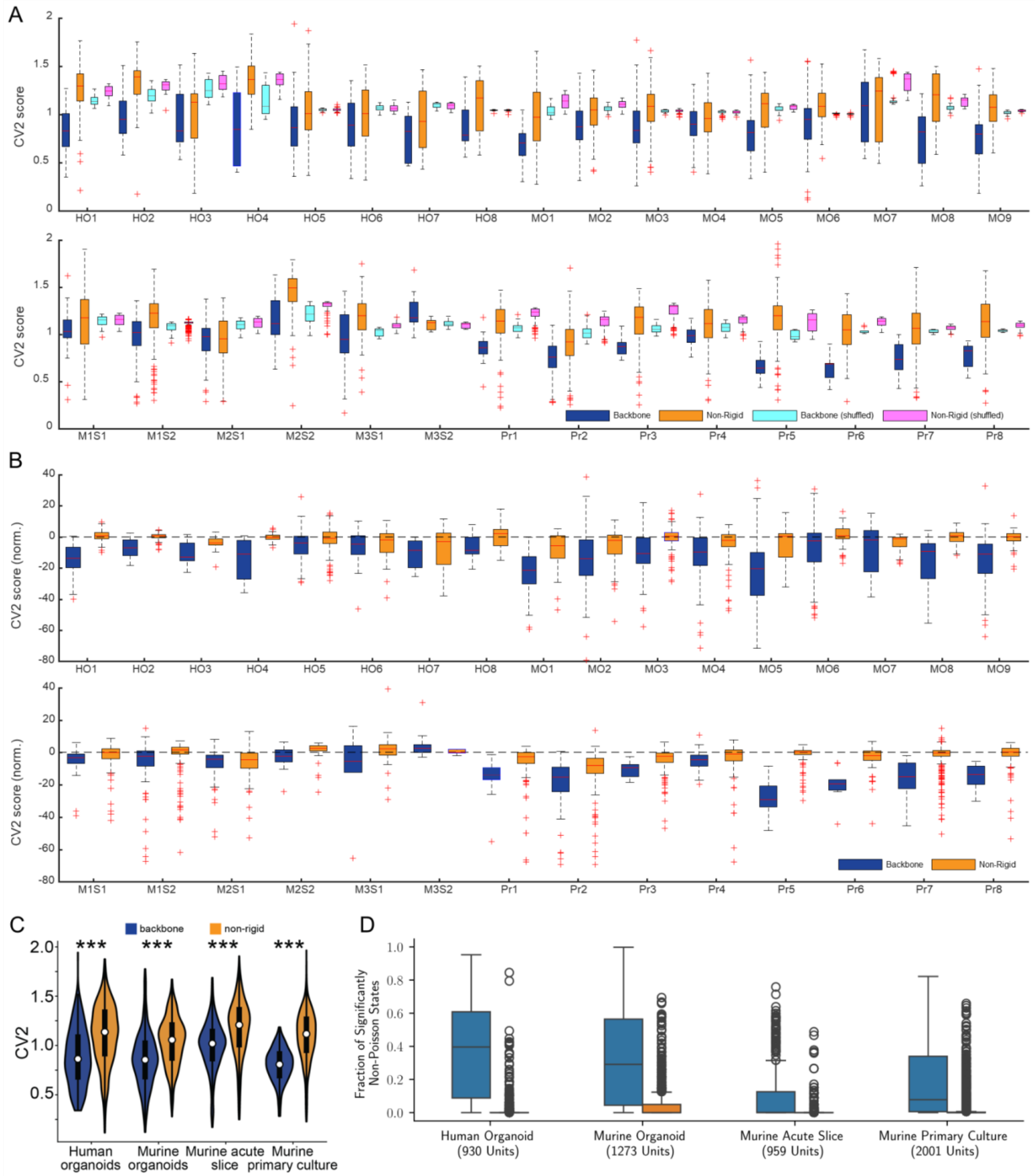


**Supplementary Figure 33. Spiking variability of backbone and non-rigid units.** (**A**) The CV2 scores of the spiking activity of each individual unit as a measure for spiking variability. The same CV2 computations were performed on 100 different shuffled spike matrices and the average over all 100 shuffled datasets are reported as well. (**B**) The CV2 scores shown in A after z-score normalization using the mean and STD of the CV2 computations applied to the 100 shuffled spike matrices. (**C**) The CV2 scores are significantly larger for non-rigid units compared to backbone units, reflecting a more Poisson-like firing of non-rigid units ($P < 10^{-16}$, linear mixed-effect model). (**D**) Distribution of the fraction of all hidden states across all fitted HMMs in which each unit exhibited lower firing variability than Poisson (according to a chi-square test described in *Methods: non-Poisson Units*). Whereas most backbone units have low variability in a sizable fraction of hidden states, most non-rigid units are indistinguishable from Poisson across almost all states.

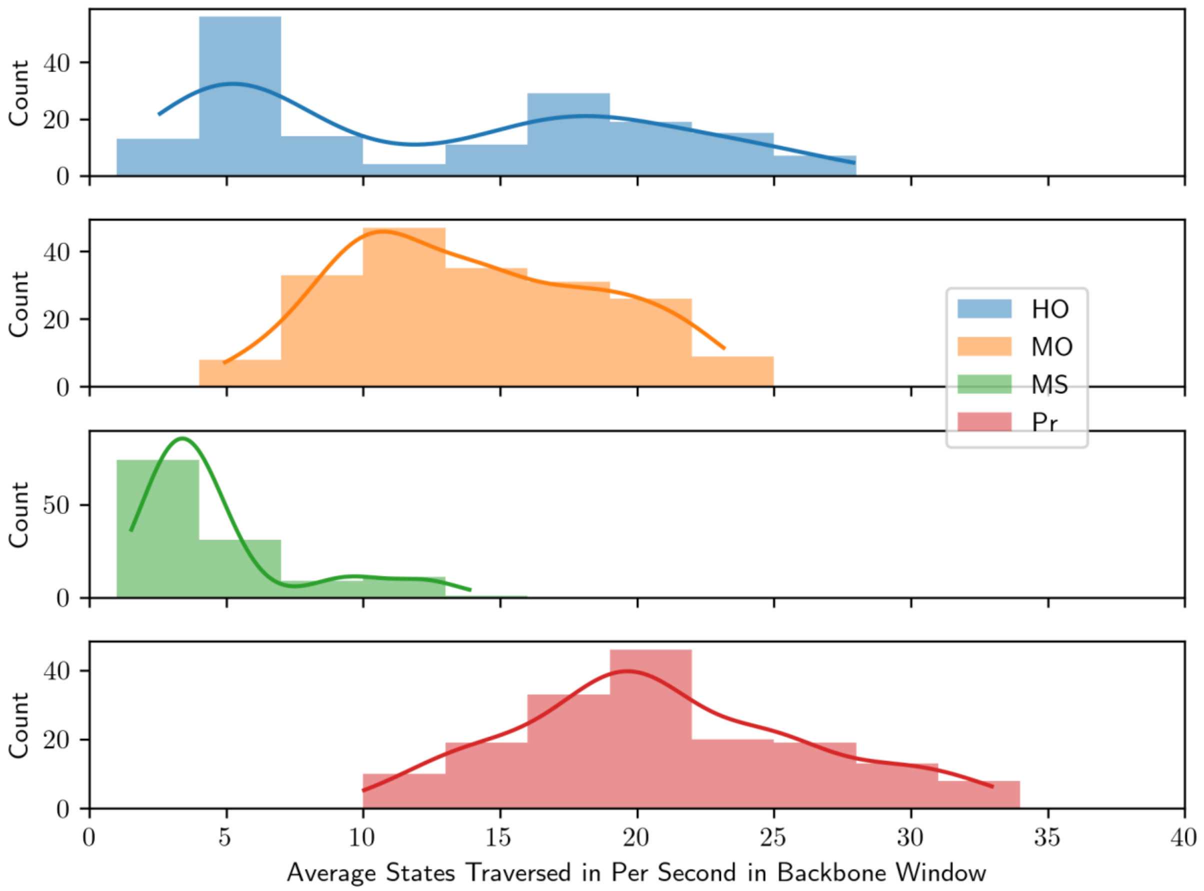


**Supplementary Figure 34. State traversal rate differs significantly between models.** For each hidden Markov model fitted to a given experiment, we averaged across bursts the number of *distinct* hidden states traversed during the backbone interval (see Methods) divided by its duration. The distributions appear substantially different, but not all are significant according to a linear mixed-effects model given the large inter-sample effects. In particular, human and murine organoid cannot be distinguished ($P = 93\%$), whereas the higher state traversal rate of murine primary cultures is significantly different from that of human organoids ($P = 2.1\%$) and murine acute slices ($P < 0.01\%$) but not significant relative to murine organoids ($P = 6.4\%$).

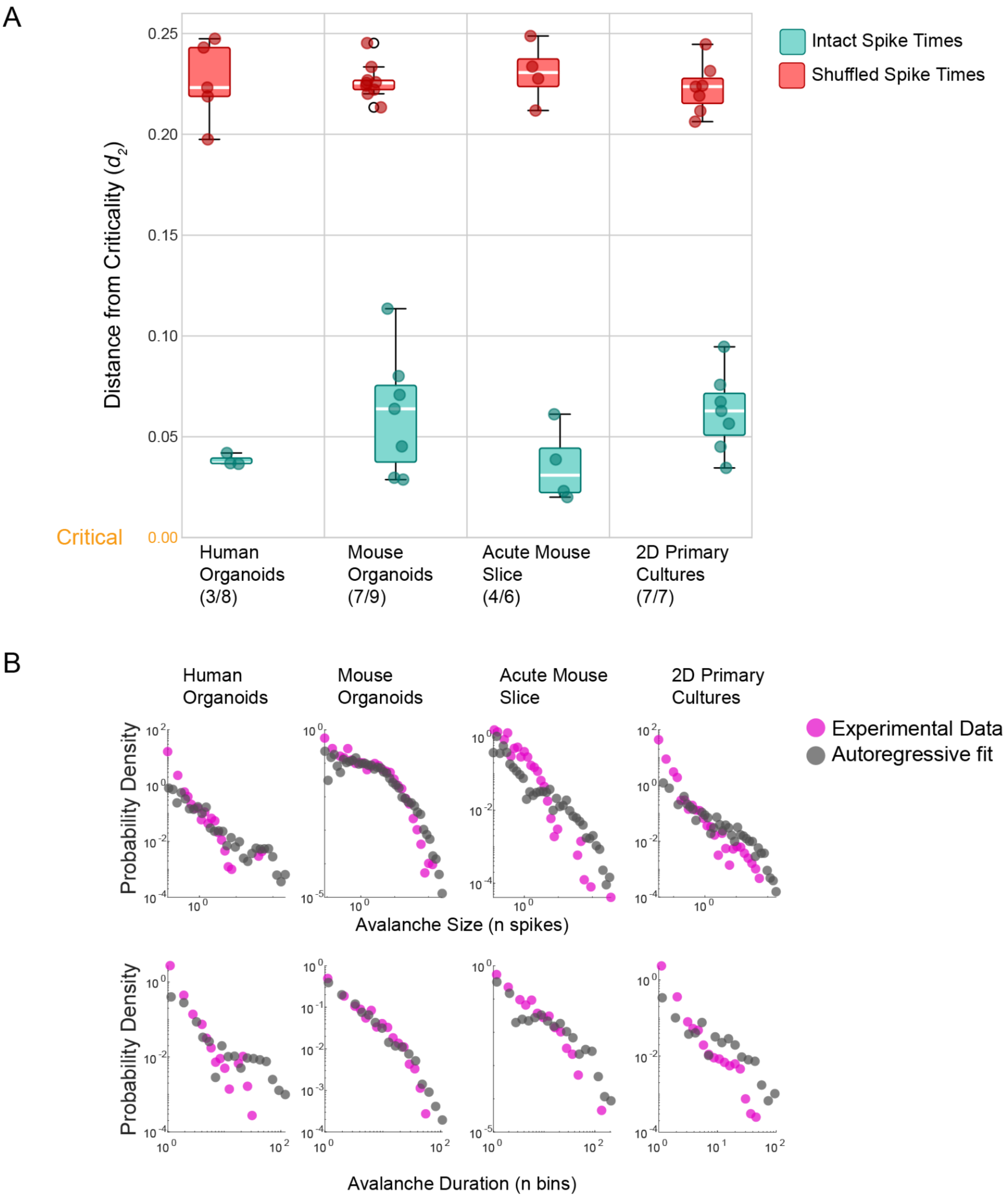


**Supplementary Figure 35. All preparations are capable of generating scale invariant dynamics consistent with a near-critical regime.** (**A**) The distance metric $d_2$ quantifies how close a system is to temporal criticality. Values of $d_2$ between ~0.0-0.1 indicate near-critical dynamics with temporal structure spanning multiple timescales. Random (shuffled) data typically yield $d_2 \geq 0.2$. A subset of preparations from each model system generated activity well-described by the autoregressive model underlying $d_2$ calculation and exhibited near-critical dynamics (teal box plots: 3/8 human organoids, 7/9 mouse organoids, 4/6 acute mouse slices, and 7/7 2D cultures). Shuffling spike times (red box plots) abolished temporal criticality (Red box plots, $P = 2.84 \times 10^{-31}$ compared to intact. 5/8 human organoids, 9/9 mouse organoids, 4/6 acute mouse slices, and 7/8 2D cultures). (**B)** Representative examples from each preparation type showing empirical data fits to the autoregressive model used for $d_2$ calculation. Activity patterns are analyzed as avalanches - bursts of spikes that are quantified by their duration (number of consecutive time bins with activity) and size (total number of spikes). Probability distributions of avalanche sizes (top row) and durations (bottom row) are shown for both experimental data (magenta) and the corresponding best-fit autoregressive model (gray). Mouse organoid data demonstrates particularly good agreement between empirical measurements and model predictions.
